# Estrogen and sex-dependent loss of the vocal learning system in female zebra finches

**DOI:** 10.1101/2020.03.28.011932

**Authors:** Ha Na Choe, Jeevan Tewari, Kevin W. Zhu, Matthew Davenport, Hiroaki Matsunami, Erich D. Jarvis

**Affiliations:** Department of Neurobiology, Duke University Medical Center, Durham, NC, 27710 USA; Department of Molecular Genetics and Microbiology, Duke University Medical Center, Durham, NC, 27710 USA; Laboratory of Neurogenetics of Language, The Rockefeller University, New York, NY, 10065 USA; The Howard Hughes Medical Institute, Chevy Chase, MD, 20815, USA

**Keywords:** specialized gene expression, transcriptomics, plumage, juvenile, development, aromatase inhibitor, sex hormones, exemestane, birdsong

## Abstract

Sex hormones alter the organization of the brain during early development and coordinate various behaviors throughout life. In zebra finches, song learning is limited to males, and the associated song learning brain pathway only matures in males and atrophies in females. This atrophy can be reversed by giving females exogenous estrogen during early post-hatch development, but whether normal male song system development requires estrogen is uncertain. For the first time in songbirds, we administered exemestane, a potent third generation estrogen synthesis inhibitor, from the day of hatching until adulthood. We examined the behavior, brain, and transcriptome of individual song nuclei of these pharmacologically manipulated animals. We found that males with long-term exemestane treatment had diminished male-specific plumage, impaired song learning, but retained normal song nuclei sizes and most, but not all, of their specialized transcriptome. Consistent with prior findings, females with long-term estrogen treatment retained a functional song system, and we further observed their song nuclei had specialized gene expression profiles similar, but not identical to males. We also observed that different song nuclei responded to estrogen manipulation differently, with Area X in the striatum being the most altered by estrogen modulation. These findings support the hypothesis that song learning is an ancestral trait in both sexes, which was subsequently suppressed in females of some species, and that estrogen has come to play a critical role in modulating this suppression as well as refinement of song learning.

## 1. Introduction

Sexually dimorphic behavior is widespread in the animal kingdom, including but not limited to courtship, mate choice, predator avoidance, and parental care, all necessary for a species’ survival (Breed and Moore, 2010). These behaviors are often highly stereotypic, suggesting that they utilize “hard-wired” pathways within the brain that are established during early development (Carrer and Cambiasso, 2009; Kurian et al., 2010; McCarthy and Arnold, 2011; Wu et al., 2009). One such behavior is vocal learning, the ability to imitate sounds heard. In its advanced form, this rare trait is found in only three groups of birds (songbirds, parrots, and hummingbirds) and five groups of mammals (humans, bats, cetaceans, pinnipeds, and elephants), and is one of the most critical components for spoken language in humans (Jarvis, 2019). The most commonly studied non-human vocal learner is the zebra finch, where the trait is highly sexually dimorphic (Nottebohm and Arnold, 1976). Like many other songbirds, male zebra finches learn their father’s/tutor’s song and use it to attract female mates; female zebra finches do not learn to sing, although like most other vertebrates, the females can produce innate vocalizations and form auditory memories of sounds heard, which she uses to evaluate the male’s song (Riebel, 2009).

This sex difference in vocal learning behavior is associated with sex differences in the zebra finch brain. Male zebra finches (and songbirds generally) have a forebrain vocal learning system comprised of seven telencephalic nuclei and one diencephalic nucleus (**Fig. 1A**) with parallels to brain pathways for spoken language in humans (Jarvis, 2019; Pfenning et al., 2014). Four of these nuclei are connected into a posterior vocal pathway (e.g. HVC and Robust nucleus of the Arcopallium [RA]) necessary for producing learned vocalizations and an anterior vocal pathway (Area X and Lateral magnocellular nucleus of the anterior nidopallium [LMAN]) necessary for imitating vocalizations (**Fig. 1A**). During development, both male and female brains start out with all four of these telencephalic song learning nuclei that are first seen between 5-15 days post hatch (PHD5-15). However, after PHD15, the male song nuclei continue to grow while the female nuclei atrophy and become barely visible, if at all, in adulthood at >PHD90 (**Fig. 1B**) (Bottjer et al., 1985; Garcia-Calero and Scharff, 2013; Konishi and Akutagawa, 1985; Nordeen and Nordeen, 1988; Nottebohm and Arnold, 1976; Shaughnessy et al., 2018). Additionally, around PHD30, a robust axon tract that extends from HVC to RA innervates RA in males, but not the atrophying RA in females (**Fig. 1A,B**) (Holloway and Clayton, 2001; Konishi and Akutagawa, 1985; Mooney and Rao, 1994).

**Fig. 1.**
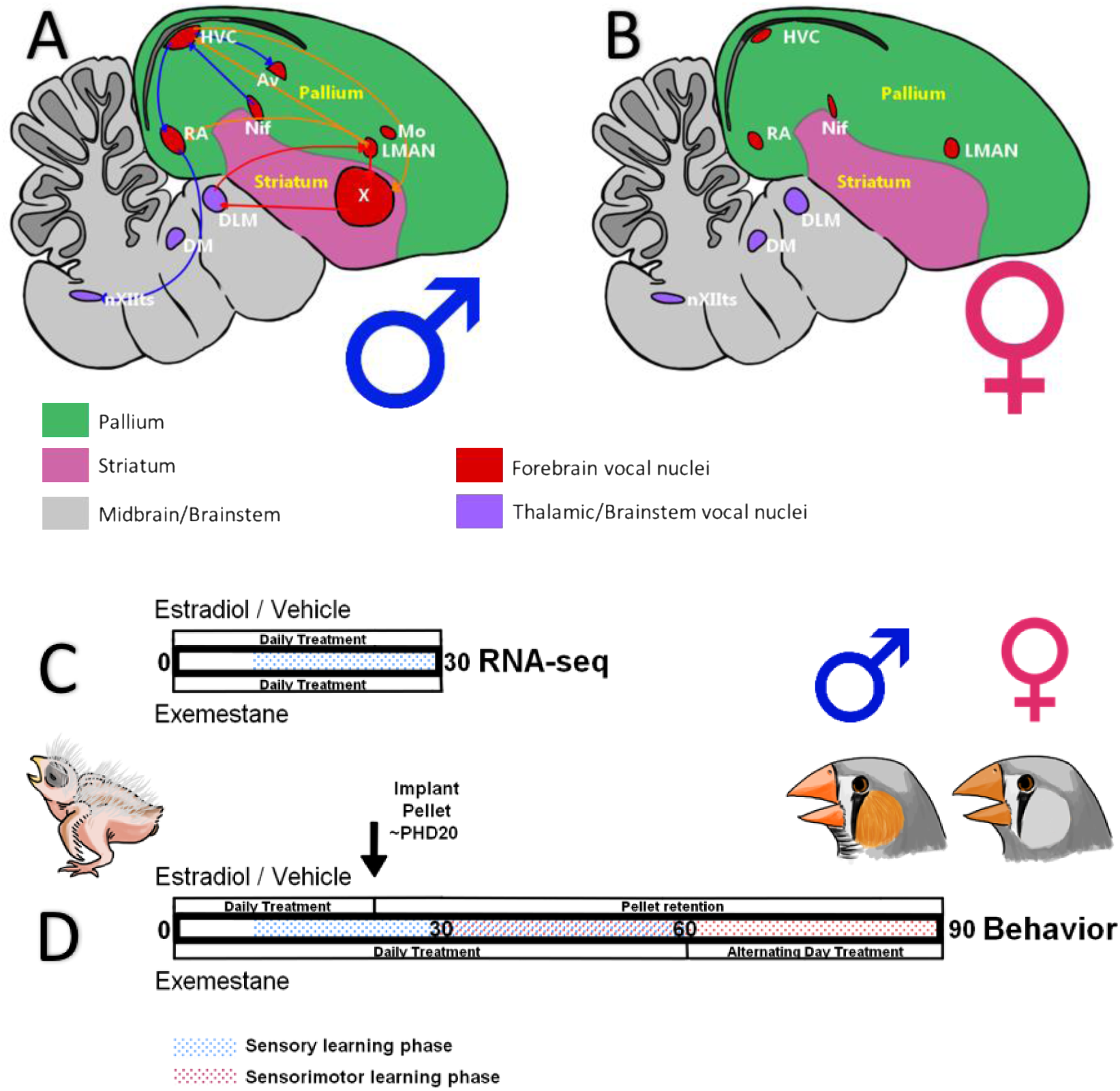
Brain organization sex differences and experimental paradigm. (A) Adult male zebra finch brain. (B) Adult female zebra finch brain. There are 7 telencephalic song nuclei: HVC, RA, LMAN, Area X, Av, Nif, and MO. Additionally, there are 3 vocal nuclei in the thalamus (DLM) and brainstem (DM and the XIIts motor nucleus). The male brain (A) has all 7 song nuclei, and the female (B) 4 confirmed song nuclei, but is missing Area X, and Av. HVC and RA are also smaller in females. LMAN is not visually different between males and females. Some connections of male song nuclei are shown; medial MAN projects to HVC. (C) PHD30 experimental timeline. Vehicle, estradiol, and exemestane treated animals were dosed daily until PHD30 and taken for brain transcriptome analysis. (D) PHD90 experimental timeline. Estradiol and vehicle animals were dosed daily until PHD20, whereupon they received silastic implants with estradiol or vehicle until sacrifice at PHD90. Exemestane animals were dosed daily until PHD60, and then every other day until behavior collection and sacrifice at PHD90. HVC, High Vocal Center; RA, Robust nucleus of the Arcopallium; LMAN, Lateral Magnocellular nucleus of the Anterior Nidopallium; Area X, proper name; Av, Avalanche; Nif, Nucleus Interface of the Nidopallium; MO, Oval nucleus of the Mesopallium; DLM, Dorsal Lateral nucleus of the Medial thalamus; DM, Dorsal Medial nucleus; XIIts, 12^th^ motor nucleus, tracheosyringeal part.

Remarkably, even a brief treatment with estrogen a week before PHD30 can cause zebra finch females to retain their vocal learning systems into adulthood, which allows these masculinized females to produce courtship song similar to males, suggesting that estrogens are potent masculinizing agents in the song system of zebra finches (Gurney, 1982; Simpson and Vicario, 1991a). In contrast, androgens are far less effective at masculinizing the female song system (Grisham and Arnold, 1995).

The reverse scenario with attempts at demasculinizing the male song system by blocking estrogens or androgens has yielded inconclusive results. Prior studies that have used fadrozole, a drug that inhibits aromatase from converting androgens to estrogens, did not reveal any impacts on male song development either behaviorally or anatomically (Merten and Stocker-Buschina, 1995; Wade and Arnold, 1994). However, fadrozole was later found to effect song behavior and aromatase activity only briefly within 30 minutes of injection; the effects were lost after 4 hours (Alward et al., 2016). It was also found that fadrozole can stabilize the aromatase protein, which is then followed by increased estrogen synthesis after drug clearance, causing a “rebound effect” (Harada and Hatano, 1998). Another drug, tamoxifen, originally thought to act only as an estrogen receptor antagonist, was found rather to enhance the song systems in zebra finches (Mathews and Arnold, 1990, 1991; Mathews et al., 1988). Later it was discovered that when tamoxifen binds to the estrogen receptor, competing with estrogen, it can either behave as an agonist or antagonist, depending on the tissue type (Martinkovich et al., 2014; McDonnell, 2005; Wardell et al., 2012; Wardell et al., 2014), behaving as a super-estrogen in the brain (Mathews and Arnold, 1991). Treatment of males with flutamide, a non-steroidal androgen receptor antagonist, failed to demasculinize any aspect of the song system, and in fact hypermasculinized RA (Schlinger and Arnold, 1991). Lastly, studies that have used selective estrogen receptor disruptors (Bender and Veney, 2008) and G-protein coupled estrogen receptor (GPER) antagonists (Tehrani and Veney, 2018) found small decreases in male song nuclei soma, decreases in HVC and Area X sizes, but possible effects on song behavior was not reported. To further complicate the issue, many studies blocked estrogen only during a limited period of development and not across the entire PHD2-PHD45 vocal learning, estrogen-responsive, critical period (Gobes et al., 2017; Konishi and Akutagawa, 1988; Pohl-Apel and Sossinka, 1984). In sum, the effects from pharmacologically blocking estrogen during development of the sexually dimorphic song system remains enigmatic.

The current hypothesis regarding the sexual dimorphism seen in songbirds is that estrogen is not critical for the development of the song system, and that the interplay between excess Z chromosome gene products and sex hormones is responsible for male-typical brain development, which is absent in the hemizygous female. Further refining this hypothesis, recent findings from Odom et al. (2014) suggest that vocal learning may have evolved in both sexes in the ancestor of all songbird species, which was then lost in the females of some species due to evolutionary pressure (Jarvis, 2004). These and the above reviewed findings lead us to suggest two alternative hypotheses: 1) That estrogen is required for development of the song learning system in both sexes of zebra finches, but prior methods missed discovering this because they did not sufficiently block it in males; or 2) Post evolution of song learning in both sexes, females of some species, such as the zebra finch, lost the trait and reversal of this sex-dependent loss is dependent on the gene-modulatory effects of estrogen signaling.

In this study, we test the plausibility of these hypotheses by utilizing exemestane, a third generation steroidal aromatase inhibitor, that has been shown to lack the “rebound” effect (Wang and Chen, 2006). Exemestane is the current gold standard for adjuvant therapy in treating estrogen sensitive cancers in the clinic due to its efficacy and specificity (Lonning and Eikesdal, 2013). We treated male and female zebra finch chicks daily with either exemestane or estradiol from the day of hatching until sacrifice, either at the beginning of the sensorimotor learning period (PHD30) or into adulthood and confirmed manipulation of estrogen levels in the blood and brain. We then conducted behavioral and anatomical comparisons, and RNA-seq analyses of song nuclei gene expression specializations. We found estrogen exerts un-equal effects in a sex-dependent, brain region-dependent manner, supporting hypothesis 2, while further revealing a requirement of estrogen for normal male plumage development, normal song learning development, and massive gene expression specializations in Area X of females during early development.

## 2. Materials and methods

### 2.1 Husbandry for hormone treated chicks

Zebra finches from our breeding colony at Duke University were kept on a 12:12 light/dark cycle between 23-29°C and 30-70% humidity. Fortified finch seed mixture (Kaytee), enriched grit (Higgins), poultry feed (Purina), cuttlefish bone and water were provided *ad-libetum*. This normal diet was supplemented with hardboiled eggs and oranges given twice weekly. Water was also supplemented with liquid calcium borogluconate (Morning Bird). Baths were given weekly with cage pan changes, and cage enrichments were provided and exchanged during cage changes (every 3 months). Breeding animals were kept in pairs, and native or foster offspring were kept with their parents until PHD60 (or sacrifice at PHD30) when the offspring have completed their sensory learning period and they are able to feed on their own. Nest boxes were cleared out and replenished with fresh nesting material between clutches.

To synchronize embryo development from clutches of multiple parents, eggs were collected daily from nest boxes during a 2-week period and placed in developmental stasis at 15°C with 80% humidity on a 30° angle rotator set to rotate once every 2 hours, in a P-008A BIO incubator (Showa Furanki Corp). Eggs were kept in stasis for no longer than 3 weeks. Egg collection from nesting pairs chosen for fostering was ceased 3 days prior to artificially incubating the synchronized eggs, permitted these foster parents to brood a clutch of 3-4 eggs. After synchronizing a cohort of eggs in the low temperature incubator, they were then moved to a higher temperature incubator at 37.5°C with ∼50% humidity on a 30° angle rotator set to rotate once per hour and incubated for up to 14-15 days. When chicks started to pip (crack the eggshell), usually beginning around incubation day 13, all eggs were transferred from the incubator’s egg rotators to the hatch plate. Within 16 hours of hatching, animals were tagged and transferred to the chosen foster nests in the aviary. To tag them, a distal toe joint of the newly hatched animals were removed with sterile forceps and a scalpel; the removed sample was used to determine the sex of the animal by PCR-genotyping with degenerate P2 (TCTGCATCGCTAAATCCTTT) and P8 (CTCCCAAGGATGAGRAAYTG) primers as previously described by Adam et al. (2014). Chicks born natively under their biological parents had their downy feathers removed in unique patterns for identification and were later sexed using cells collected from buccal swabs during banding (∼PHD 10). Nests were limited to no more than 5 chicks to reduce nestling mortality. Animals were separated into two colonies: one for animals treated with exemestane or estradiol, and the other with the main colony for those treated with vehicle. To prevent cross contamination of drugs by contact between animals, exemestane and estradiol treatments were not done concurrently and treatment with the opposing drug was not conducted until 48 hours had passed following the removal of the active compound and the cage was replaced with a cleaned and sanitized one.

### 2.2 Pharmacological manipulations and experimental timelines

Exemestane is trademarked by Pfizer as Aromasin (Pfizer, 2018), which irreversibly binds to the testosterone binding site of aromatase and alters the confirmation of the protein for ubiquitin mediated degradation, pharmacologically removing the sole enzyme responsible for estradiol and estriol synthesis (Lonning and Eikesdal, 2013). Exemestane (Sigma PHR1634) was dissolved in DMSO (PanReac Applichem 191954) at a concentration of 100mg/mL, which was then suspended in olive oil (Sigma 75343) for a final concentration of either 10mg/mL or 20mg/mL, which is needed for prolong absorption as done with sesame oil in quail (Çiftci, 2012) and rat (Theodorsson et al., 2005). Vehicle was the same solution without exemestane. Exemestane or vehicle treatments were given daily via subcutaneous injection with a 28.5-gauge needle from PHD0 until sacrifice at PHD30 (**Fig. 1C**), or until ∼PHD60, after which the treatments were given every other day until sacrifice at ∼PHD90 (**Fig. 1D**). Doses were given between 10 and 60 ug/g body weight (between 5uL and 30uL) and not exceeding 100ug/g, as doses over 125 ug/g (mg/kg) in mammals are toxic (Pfizer, 2018).

Estradiol (E2) is the most potent form of estrogen. As with exemestane above, estradiol (Sigma E1024-1G) was also dissolved in DMSO at a concentration of 100mg/mL, which was further suspended in olive oil to a final concentration of 1mg/mL. Initially in a pilot experiment, we treated chicks with daily subcutaneous estradiol injections at 20 ug/g body weight, but this resulted in high mortality even after lowering the dose to 5 ug/g body weight. Therefore, we later transitioned to daily topical treatments with one drop (∼30-50uL) of the 1mg/mL solution applied near the flank as this was the easiest and least invasive route of treatment. Identical topical daily treatments with the estradiol solution or vehicle were given at doses no higher than 50ug of estradiol (or equivalent vehicle volume) until ∼PHD14, and then the solution was given every other day until time of sacrifice for animals taken at PHD30 (**Fig. 1C**), or until PHD20 for animals intended for sacrifice at PHD90 (**Fig. 1D**). For the latter group, at ∼PHD20, instead of topical application, silastic pellets were implanted. These implants were made by mixing medical grade silicone adhesive (Nusil MED-1037) with estradiol dissolved in DMSO 100mg/mL or adhesive mixed with DMSO alone as a vehicle control. The mixture was extruded from syringes into ropes that were cured overnight. The resulting pellets were cut, weighed, and kept in sterile conditions at 4°C until use (Gurney, 1982; Sahores et al., 2013; Simpson and Vicario, 1991a). Each implant carried approximately 150-200ug of estradiol, and vehicle implants were size matched. The silastic pellets were surgically placed subcutaneously on the flank under the wing and retained until sacrifice date at ∼PHD90. PHD20 was the earliest the birds could reliably hide the surgical site from their parents. This approach was less detrimental for animal health, as we found that daily or alternating daily topical application past PHD30 resulted in animals with bones too fragile to properly fly or walk.

To carry out the implant surgeries, prior to surgery, the animals were given meloxicam (Metacam NDC 0010-6013-01, 5mg/mL) intramuscularly at 0.3mg/kg body weight. They were then initially anesthetized with 3-4% isoflurane (Isothesia NDC 11695-6776-1) in 100% oxygen, and then sustained with 1.5-2% isoflurane for the duration of the procedure. The surgical site was plucked, and the exposed skin was scrubbed with 70% ethanol and 10% povidone-iodine prior to the creation of a shallow incision. A pocket was created under the skin using a blunt hemostat, and the implant was placed in the pocket. The incision was sealed with veterinary adhesive (3M Vetbond, 1469SB or Henry Schein Vetclose 031477), and bupivacaine (Hospira NDC 0409-1159-01, 0.25%) was applied topically afterwards. The animals were observed continuously for the first 2 hours, and daily afterwards. The animals were also given additional intramuscular meloxicam 24 hours and 48 hours after surgery to manage pain. When handling estradiol, in addition to normal standard lab protective clothing, experimenters wore N-95 respirator masks to avoid inhaling any possible estradiol-contaminated airborne particles.

On the day of sacrifice, all experimental animals were kept in the dark for approximately 1 hour to return most brain gene expression activity to baseline levels (Jarvis and Nottebohm, 1997; Whitney et al., 2014). For animals collected at PHD30, they were separated from their parents for 1 hour to rest in the dark. The conditions of adults are described below for vocalization analyses. For both adults and juveniles, the animals were rapidly anesthetized via isoflurane inhalation overdose followed by rapid decapitation when the animals were unresponsive, to reduce possible stress-induced brain gene expression. The brains were quickly dissected and embedded in OCT before snap freezing in a slurry of dry ice and ethanol. Trunk blood was collected from the neck at sacrifice and left at room temperature for approximately 20 minutes to clot before centrifuging at 20,000g for 3 minutes to separate the cells from serum. Gonads were examined post-mortem to confirm sex.

### 2.3 Vocalization behavior analyses

For animals sacrificed at ∼PHD90, vocalization behavior was recorded starting at PHD60, when the animals were weaned and past the sensory learning period of song development (Kojima and Doupe, 2007). They were recorded continuously in sound isolation chambers where they could neither see nor hear other animals. At ∼PHD90, experimental animals (males and females) were provided with a novel female to collect directed songs (males and estradiol treated females) and other vocalizations for over 2 hours, regardless of sex or treatment. After this period of directed vocalization, the novel females were returned to their home-cage and the experimental animals were kept in the dark for approximately 1 hour to fast and return brain activity to baseline. The animals were then sacrificed.

One estradiol treated PHD90+ female underwent audio/visual recording with a novel control female as a visual stimulus. Both animals were able to hear the other but were separated by an electrochromic glass plate that could be turned opaque or transparent remotely. This estradiol treated female underwent several sessions for 5 hours, with 30-minute intervals of seeing and 1-hour of not seeing the stimulus female. This estradiol treated female was later returned to her sound isolation home cage overnight before conducting recording behavior the same as all other experimental animals.

Recordings were collected using (Earthworks SR69 or SRO) microphones and an Aardvark 24/96 Pro pre-amplifier, connected to a computer operating Windows XP sp3. Avisoft recording software was used to gate and record sounds that were due to vocalizations, namely with an energy threshold of >1%, entropy threshold of <70%, and duration of >3 milliseconds. Recordings included 500 milliseconds before and after the triggering event. Avisoft Recorder v4.2.18 (Avisoft Bioacoustics) generated sound files were opened in Raven Lite 2.0 interactive sound analysis software (Cornell lab of Ornithology) and examined manually by blinded personnel to identify sound files with excessive cage noise or sound files with suitable vocalizations for automated thresholding in subsequent analysis. Syllable characterization and quantification was performed in Sound Analysis Pro 2011 (SAP2011) (Tchernichovski et al., 2000). Syllables were segmented and had their feature characteristics tabulated through automated batch analysis functions included in SAP2011. These characteristics were tabulated across development for all individuals. Syllables were then visualized through Nearest Neighbor Hierarchical clustering functions provided in SAP2011, and the discrete clusters separated in Euclidian space were identified manually by 2 blinded evaluators.

### 2.4 Protein modelling of aromatase binding site

The complete amino acid sequences of human aromatase (UniprotKB: P11511) and zebra finch aromatase (UniprotKB: Q92112) were taken from the UniProtKB database and aligned using the UniProt alignment tool (https://www.uniprot.org/align). The model of zebra finch aromatase was built via homology modeling using Modeller (v9.23) based on a mono template of the human experimental structure bound to exemestane and the heme co-factor (Protein Data Bank (PDB) ID: 3S7S). Visualization of aromatase bound to exemestase and the heme co-factor was performed on the Visual Molecular Dynamics (VMD v1.9.3) program.

### 2.5 Steroid panel assay with high performance liquid chromatography and tandem mass spectroscopy

To test the efficacy of exemestane in zebra finches, adult animals were subcutaneously injected with exemestane or vehicle (60-40ug/g body weight: 600-800ug) for three days, and then sacrificed 24 hours after final treatment. Animals were euthanized in the same fashion as outlined above. Serum and whole brain samples were submitted to the metabolomics core facility at Duke university. Tissue homogenization and sterol extraction was performed and these samples were assayed for a complete steroid panel analysis (Cortisol, Cortisone, 11-Deoxycortisol, 17α-Hydroxyprogesterone, Progesterone, Aldosterone, Corticosterone, 11-Deoxycorticosterone, Estradiol, Estrone, Androstenedione, Androsterone, Dehydroepiandrosterone, Dehydroepiandrosterone sulfate, Dihydrotestosterone, Etiocholanolone, and Testosterone), using the AbsoluteIDQ Stero17 kit (BioCrates) on the Xevo TQ-S MS UPLC/MS/MS instrument (Waters Corporation). The full extraction and assay protocol is included in **Supplemental Note 1**.

### 2.6 ELISA assays to measure blood estrogen levels

Collected serum was serially diluted in ultrapure water (Invitrogen 10977015) and used directly in the estradiol enzyme immunosorbent assay (EIA, Cayman Chemical 582251) following the manufacturer’s protocol (Vedder et al., 2014). A total of 4 dilutions were run in triplicate for each exemestane treated animal (1:5, 1:10, 1:20, 1:40) and for each vehicle or estrogen vehicle treated animal (1:10, 1:20, 1:40, 1:80). Results were obtained using a SpectraMax M3 micro-plate reader (Molecular Devices) with Softmax Pro software v6.2.1 on a computer operating Windows 7 professional.

### 2.7 Cresyl violet staining histology

The left hemisphere of brains frozen in OCT were sagitally sectioned over 9 serial slides. These sections were cut at 14-16um thickness on a microtome cryostat (Leica CM1850), thaw mounted onto charged borosilicate slides (Fisherbrand Superfrost #12-550-15) and stored at -80°C until use. One series was dehydrated and rehydrated in graded ethanols (0%, 50%, 70%, 95% 100%), stained in 0.3% cresyl violet acetate (Sigma C5042), defatted in mixed xylenes (Fisher X5), cover-slipped with permount (Fisher SP15) mounting media and cured for one week in a chemical cabinet prior to imaging on a stereomicroscope (Zeiss Stemi 305) equipped with a color camera (Zeiss Axiocam 105). Images were obtained on a computer operating Windows 7 using Zeiss Zen Blue 2.0 software.

### 2.8 Chromogenic in-situ hybridization

Previously cloned *CADPS2* plasmids (Accession: DV955943) were grown from bacterial stock and collected via miniprep columns (Qiagen 27104). *CADPS2* was verified for sequence identity and orientation using Sanger sequencing services provided by Eton Biosciences or GeneWiz. We performed a modified version of the *in-situ* hybridization protocol as first described by Takatoh et al. (2013)(Biegler et al., in preparation). Template DNA was PCR amplified from plasmids using M13 forward and reverse primers and Phusion high-fidelty DNA polymerase (Thermofisher F530S). The target product was gel purified using the NucleoSpin mini-spin columns (Machery-Nagel 740609.50). DIG-labelled (Roche 11277073910) RNA probes were transcribed via the T3 promotor from 1ug of purified DNA template and cleaned via ethanol-salt purification with GenElute linear polyacrylamide (Sigma 56575-1ML) as a neutral carrier. The probe pellet was rehydrated in 100uL of 90% formamide, and frozen in 5uL aliquots at -80°C. Probes were only freeze-thawed once to prevent RNA degradation.

Slides with fresh-frozen sections were fixed in freshly prepared 4% PFA/1X PBS for 5 minutes at room temperature. The slides were then washed in 1X PBS and acetylated in 0.1M triethylamine + acetic acid for 10 minutes before they were dehydrated in a series of graded ethanols. The opaque tissue sections were outlined with hydrophobic marker (Thermofisher 008899) and then prehybridized (50% formamide [ThermoScientific 15515026], 5X SSC [ThermoScientific AM9763], 1X denhardt’s solution [Sigma D2532], 250ug/mL Brewer’s yeast tRNA [Roche 10109495001], 500ug/mL herring sperm DNA [ThermoScientific 15634017]) with parafilm coverslips at room temperature for 1 hour in a humidified chamber. The hybridization-probe solution was made with probe diluted 1:100 (from frozen aliquots outlined above) in hybridization buffer (300mM NaCl, 20mM Tris-Hcl pH 8.0, 5mM EDTA, 10M Na2HPO4 pH 7.2, 10% dextran sulfate, 1X denhardt’s solution [Sigma D2532], 500ug/mL Brewer’s yeast tRNA [Roche 10109495001], 200ug/mL Herring sperm DNA [ThermoScientific 15634017], 50% formamide [ThermoScientific 15515026]) hydrolyzed at 80°C for 6 minutes before being chilled on ice before application. After prehybridization was complete, the parafilm coverslips were removed, the excess prehybridization buffer was removed with a kimwipe. The prepared hybridization-probe solution was applied and then the slides were coverslipped with glass coverslips (VWR 48393106). The slides were then incubated at 65°C in a hybridization oven overnight (>16 hours) in a humidified chamber. Humidification was done using either RNase free water or a solution of 5X SSC, 50% formamide.

After overnight hybridization, the coverslips were gently removed in room temperature 5X SSC. The hydrophobic pen residues were wiped off using kimwipes. The slides were then washed in 5X SSC at 68°C for 10 minutes, and then washed 4 times for 30 minutes per wash in 0.2X SSC at 68°C. After the final wash, the container holding the 0.2X SSC and slides were allowed to cool to room temperature. The slides were washed once more in fresh 0.2X SSC at room temperature.

After the high stringency washes, the slides were washed in Buffer B1 (0.1M Tris pH=7.5, 0.15M NaCl) for 5 minutes at room temperature before being transferred to slide mailers. The slides were then incubated in blocking Buffer B2 (10% Sheep serum [Sigma S3772] in Buffer B1) at room temperature for a minimum of one hour. After blocking, the slides were incubated in an AP-conjugated anti-DIG antibody solution ([Roche 11093274910] diluted 1:2000 in 1% sheep serum in Buffer B1) overnight at 4°C.

Following antibody incubation, the slides were washed 3 times for 10 minutes per wash in Buffer B1 at room temperature 3 times and then equilibrated in 100mM Tris-HCl pH-9.5 for 5 minutes. The slides were then incubated in a working solution of NBT/BCIP (Vector labs sk-5400), prepared according to the manufacturer’s protocol, for 16 hours in the dark at room temperature.

When the NBT/BCIP signal was optimal, the reaction was stopped in 1X PBS, and the slides were washed 3 times for 5 minutes per wash in 1X PBS at room temperature. The slides were then rinsed in diH_2_O and counterstained in a 1:3 diluted solution of Nuclear fast red (Vector labs H-3403). Counterstaining was done for no longer than 3 minutes. The slides were then rinsed with diH_2_O and dipped in 100% histology grade ethanol (no more than 10 dips) before being left to airdry. Once the dried sections were opaque, the slides were mounted with Vectamount permanent mounting solution (Vector labs H-5000) and dried overnight at room temperature in a dark, dry location before imaging.

### 2.9 Imaging and area calculations

Cresyl violet and *in-situ* hybridized sections were imaged on a stereomicroscope (Zeiss Stemi 305) equipped with a color camera (Zeiss Axiocam 105). Images were obtained on a computer operating Windows 7 using the Zeiss Zen Blue 2.0 software. Images were saved as .czi files (Zeiss) and area size values were obtained using the “region of interest” tools available in Zen Blue 3.0 (Zeiss). The area of song nuclei (Area X, HVC, LMAN and RA) and several brain subdivisions (striatum, arcopallium and mesopallium) were obtained from sections stained with either Cresyl violet or *CADPS2* as a marker gene. Sections were selected based on anatomical landmarks to compare across individuals. Song nuclei and brain subdivision areas were divided by the area of the whole telencephalon within each respective section. Brain subdivisions were determined according to the online zebra finch histological atlas from the Mello lab (http://www.zebrafinchatlas.org) and molecular markers established by Feenders et al. (2008) and Jarvis et al. (2013).

### 2.10 RNA-Seq data generation

Following a modified protocol of Whitney et al. (2014), the right hemisphere of brains frozen in OCT were sectioned coronally for regions of interest at 14um and thaw mounted onto polyethylene naphthalate (PEN) membrane slides (Applied Biosystems LCM0522). As soon as the sections dried, the slides were promptly stored at -80°C until further use. A series of adjacent sections on regular glass slides were stained with cresyl violet stained and used as reference to identify sections with song nuclei (**Supplemental Fig. 1**). Processing one PEN slide at a time, sections containing the target song nucleus and its immediate surround were taken out of -80°C and submerged at -20°C in 75% ethanol. The slide then was dehydrated in ice cold graded ethanols: 75%, 95%, 95% (second rinse), 100%, 100% ethanol (second rinse), for 10 dips each, before incubating in fresh xylenes twice for 5 minutes each incubation. After the second xylene incubation, the slide was air dried and the desired regions were collected one brain region at a time across multiple sections, using laser capture microscopy (LCM) on an ArcturusXT LCM system (Nikon) with CapSure Macro LCM caps (Applied Biosystems LCM0211). These regions were Area X, the medial striatum (MSt) ventromedial to Area X, HVC, HVC shelf ventral to HVC, LMAN, anterior nidopallium lateral to LMAN (LANido), RA, and lateral intermediate arcopallium (LAi) lateral to RA. Care was done to protect samples from RNase degradation by using only RNase-free materials/reagents and cleaning all reusable equipment with RNase Zap (Invitrogen AM9780). All collection was done within 35 minutes upon exposure to open air.

RNA was isolated from the LCM collected tissues using the Arcturus Picopure kit (Applied Biosystems KIT0204) following manufacturer’s instructions. RNA quality was determined using an Agilent 2100 bioanalyzer with the high sensitivity RNA 6000 pico kit (Agilent 5067-1513). Only samples with RIN numbers higher than 5 were used for further processing.

cDNA was synthesized using the SMART-Seq v4 Ultra Low input RNA Kit (Takara 634892) following the manufacturer’s protocol. The cDNA product was validated using an Advanced Analytical fragment analyzer (Agilent) with the HS NGS 1-6000 fragment kit (Agilent DNF-474). Sequencing libraries were created using the NEBNext Ultra II DNA Library Prep kit for Illumina sequencing (New England Biolabs E7645L). All cDNA and library clean-up was done using SPRIselect beads (Beckman Coulter B23317).

Sequencing services were conducted by Novogene Co., Ltd. on the Novaseq 6000 platform (Illumina) via the s4 flow cell for 150bp paired-end reads. The resulting reads were aligned to the TaeGut 3.2.4 zebra finch genome assembly (GCA_000151805.2) using two independent approaches in custom designed pipelines: Kallisto (Bray et al., 2016) with reads aligned to cDNA transcripts from Ensembl; and the splice aware STAR (Dobin et al., 2013) to the annotated zebra finch genome in NCBI (Warren et al., 2010). In both analyses, the output of read counts for all genes were called. The read counts were comparable with both methods, but we saw some differences for a small number of genes. We only included in our analyses genes with consistent read count differences using both methods.

### 2.11 Statistics

Estradiol measurements did not fit a normal distribution and were thus analyzed using the non-parametric Kruskal-Wallace tests with post-hoc Steel-Dwass tests for each pair comparison or Wilcoxon rank sum tests. Song-syllable cluster data was normally distributed, and thus analyzed using standard least squares models for a 2-way ANOVA with post-hoc Tukey’s Honest Significant Difference (HSD) tests for each pair comparison. Song nuclei:surround or song nuclei:telencephalon area ratio measurements were analyzed using Aligned-ranks transformed ANOVA (Wobbrock et al., 2011) with post-hoc Wilcoxon tests for each pair.

For the RNA-seq analyses, to compare expression across all genes in all samples, including those without formal gene names, we normalized the data by via variance stabilization transformation and then performed principle component analysis (PCA). To identify significant differentially expressed genes (DEGs) between the song nuclei and their surrounding brain subdivisions, we took the Kallisto and STAR generated read counts and applied them to the DESeq2 package (Love et al., 2014) in R and performed two types of DESeq2 analysis using the Wald test: unpaired group comparisons between brain regions for each of the six groups of birds (Area); and paired comparisons between brain regions for each bird (Subject + Area). Genes with FDR ≤ 0.05 were considered significantly DEG. DEG log2-fold change values greater than 2 were binned at |2| for generating heatmaps. We also removed DEG genes identified as singing-regulated in each song nucleus within 1 hour of singing (40-67 genes) by Whitney et al. (2014), to remove potential differences associated with vocalizing behavior in the hours before sacrifice as opposed to real baseline differences. Scripts for DESeq2 and downstream analysis are in **Supplemental Note 2**. All statistical functions not included in the default DESeq2 software as-is, were carried out using one of two software/programming packages: JMP 13 or R Studio with various packages (**Supplemental Note 3**).

## 3. Results

Our experiments were designed to test the plausibility of two alternative hypotheses: 1) That estrogen is required for development of the song learning system in both sexes of zebra finches; or 2) Post evolution of song learning in both sexes, female zebra finches lost the trait and reversal of this sex-dependent loss has become dependent on estrogen signaling. We first determined whether exemestane can inhibit estrogen production in zebra finches and then assessed the potential effects of this inhibition on development of vocal learning behavior, vocal learning brain regions, and examined associated genes.

### 3.1 Exemestane is a potent blocker of estrogen synthesis in zebra finch blood and brain

In mammals, exemestane is a well-established inhibitor of aromatase, the sole enzyme responsible for synthesizing estradiol and estrone from androgens, but this drug’s efficacy has not been demonstrated in birds. When we compared the human and zebra finch aromatase amino acid sequences, they were 75% identical (**Fig. 2A**). Structural homology modelling revealed that the binding site of testosterone/exemestane in aromatase was highly conserved between human and zebra finch (**Fig. 2B**), with 11 of the 12 amino acids in the site being identical (**Fig. 2A**). The single non-conserved amino acid, at position 373 in the alignment, was a valine in humans and an isoleucine in zebra finches, both being two of only three branch chain amino acids. The high amount of conservation and nominal discrepancy in the binding site structure of human and finch aromatase strongly suggested that exemestane should bind with and inhibit aromatase similarly in the two species.

**Fig. 2.**
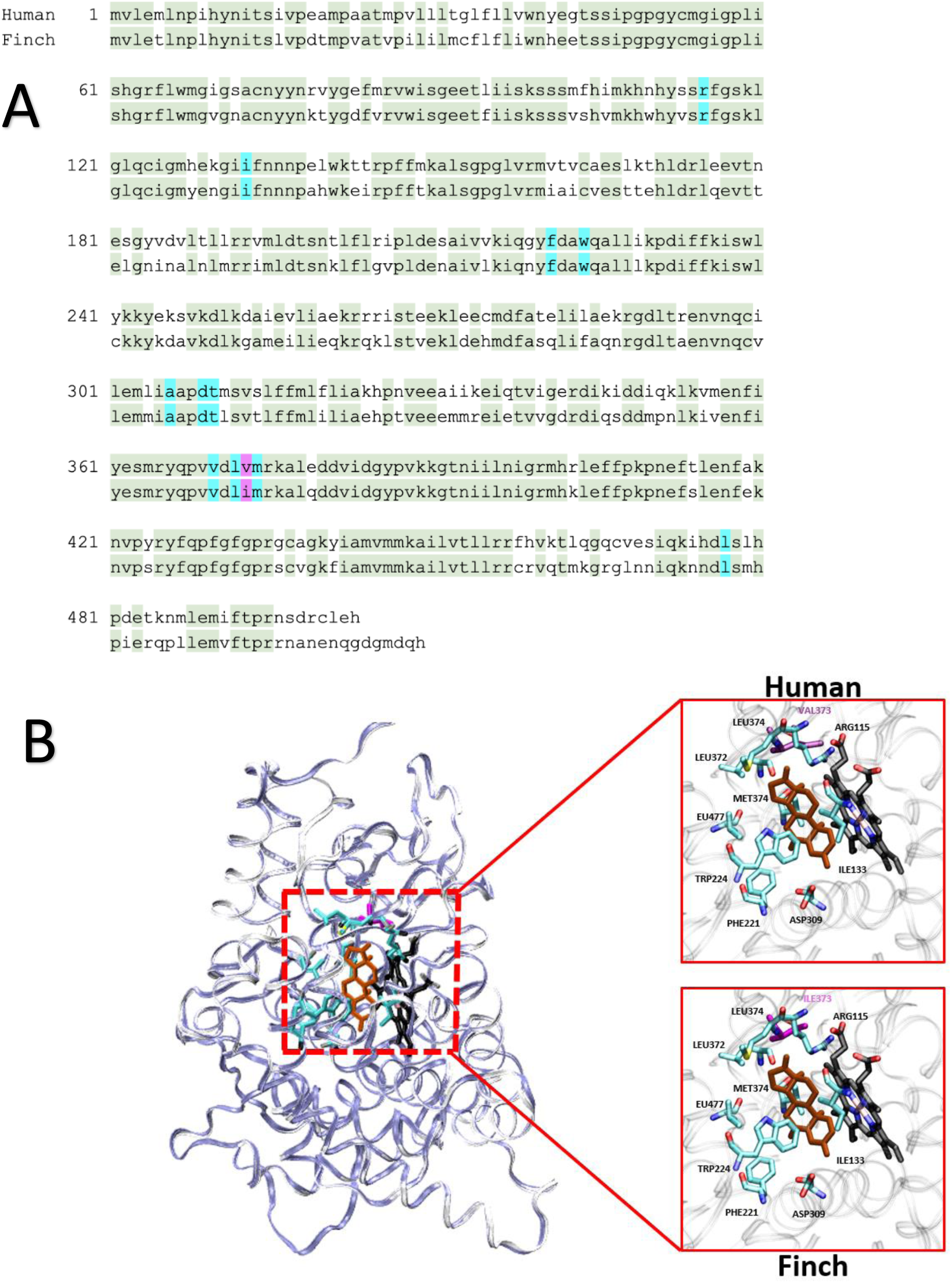
Exemestane binding site in human and zebra finch aromatase. (A) Primary protein sequence alignment between human and zebra finch aromatase protein (CYP19A1). Light green, conserved amino acids outside the ligand binding site. Cyan, conserved amino acids in the ligand binding site. Magenta, non-conserved amino acid in the binding site. (B) Superimposed tertiary structure of human (PDB ID: 3S7S – white ribbon) and finch (built by homology modeling – iceblue ribbon). Binding sites are magnified in right insets. Cyan, conserved residues between human and finch. Magenta, non-conserved amino acid. Black, the cofactor heme (HEM) group. Orange, exemestane molecule.

To test the effectiveness of exemestane in modulating estrogen levels in the zebra finch *in vivo*, we treated healthy adult male and female breeding animals (n = 5 each) for three days with either exemestane or vehicle and examined steroid levels from serum and brain tissue using an uHPLC-MS/MS assay. We found that in all birds treated with exemestane, there was no detectable estradiol in the brain (**Fig. 3A**) and very low levels in serum (**Fig. 3B**), whereas in vehicle treated animals some of them had moderate to high estradiol levels (full statistics in **Supplemental Table 1)**. There were no detectable differences in estradiol levels between adult males and females. Analyses of seven other steroids showed a small but statistically significant decrease of aldosterone in female brains and progesterone in male serum, alongside an increase in cortisone in male serum, but no other changes (**Supplemental Fig. 2; Supplemental Table 1)**. These findings indicate that exemestane is a potent inhibitor of estrogen synthesis in zebra finches, with limited off target effects.

**Fig. 3.**
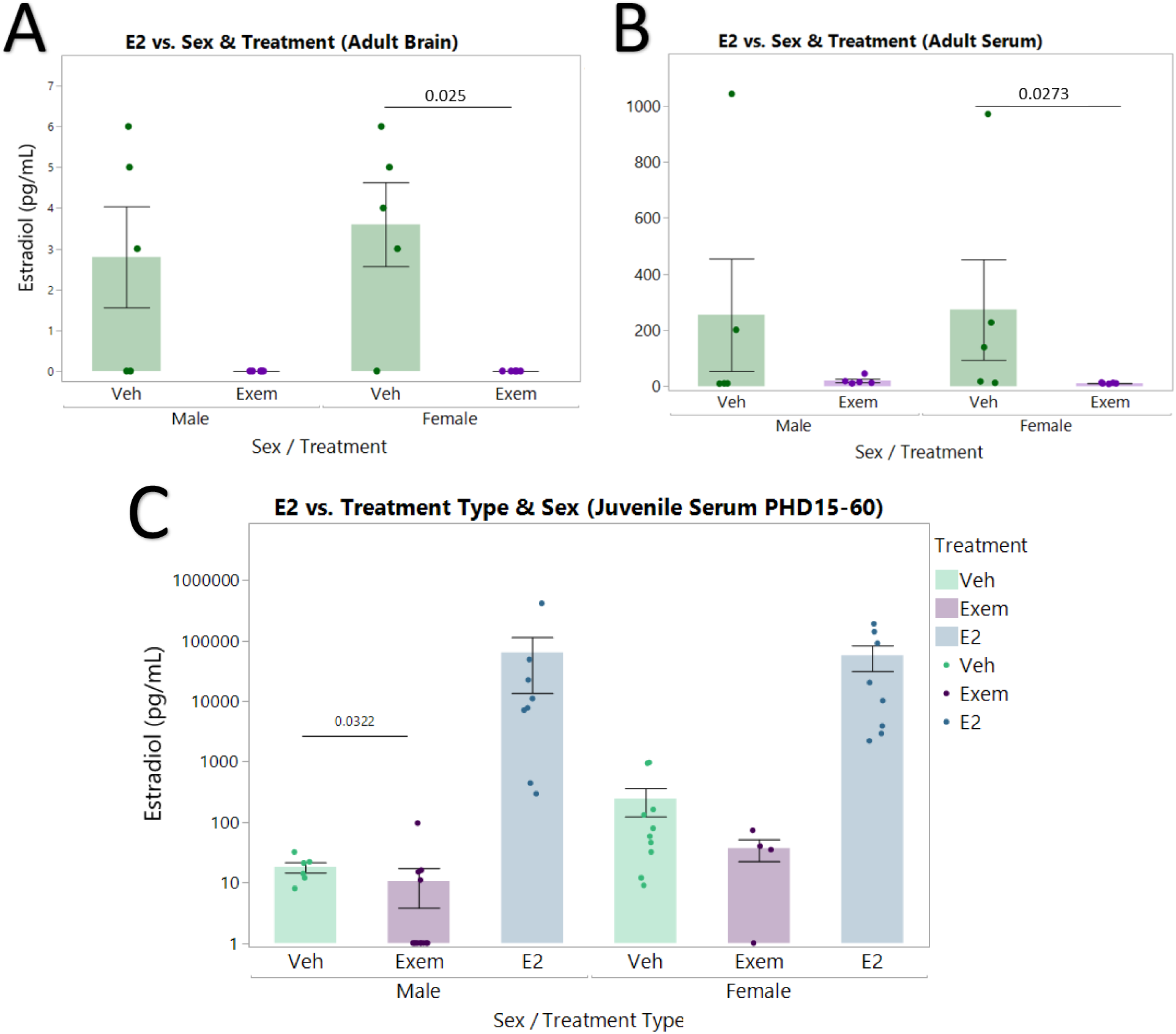
Brain and blood estradiol levels in treatment groups. (A) Adult brain uHPLC-MS/MS. None of the exemestane treated animals have detectable E2 levels in the brain. Wilcoxon test (n = 5) (B) Adults serum uHPLC-MS/MS. All of the exemestane treated animals have barely detectable E2 levels in the serum. Wilcoxon test (n = 5) (C) Juvenile serum EIA. E2 treated animals had significantly higher serum E2 levels when compared to vehicle treated animals, for nearly all animals. Exemestane treated animals had significantly lower serum E2 levels when compared to vehicle treated animals, for all animals, in males alone, but not in females alone. Kruskal-Wallis test with Steel-Dwass post-hoc (n = 4 to 14). Full statistics in **Supplemental Table 1**.

In juvenile (∼PHD15-60) experimental animals that underwent long-term daily treatments with exemestane, estradiol, or vehicle (n > 4 each group), we measured serum estradiol levels by EIA assays. The EIA assays appeared to be more sensitive at detecting differences in serum estradiol levels, requiring a log-scale analyses (**Fig. 3C, Supplemental Table 1**). We found that juvenile females had on average 10 times more estradiol than males, with overlapping ranges. Most exemestane treated juvenile males (n = 10) had no detectable serum estradiol, whereas a minority (n = 4) had levels similar to vehicle controls. A single exemestane treated juvenile female (n=1) had no detectable serum estradiol, but some (n=3) had levels that overlapped with vehicle controls. When we consider all exemestane treated animals at both ages for both sexes, the majority of them (84%) had very low to no detectable serum estradiol levels when compared to vehicle treated animals. We surmise that the minority of animals where we did not see a decrease to undetectable levels was due to the treatment regimen not being optimized for complete aromatase suppression in all animals. Juveniles treated daily with estradiol had ∼380-fold elevated serum estradiol levels compared to vehicle controls. (**Fig. 3C, Supplemental Table 1**). These data demonstrate that chronic long-term treatment of exemestane or estradiol results in dramatic changes in estrogen levels. We note a high degree of variability is seen in estradiol levels from both adult and juvenile vehicle treated control samples using two independent assay methods, suggesting that the variability is biological as opposed to technical.

### 3.2 Estrogen is necessary for normal male plumage development

We found that males treated with exemestane showed impaired male-specific plumage development after their first molt around PHD60, displaying patterns that were intermediate between normal (or vehicle treated) males and females (**Fig. 4A-C)**. This includes weak to no orange cheek patches, and minimal zebra striped throat feathers. By PHD90, all long-term exemestane treated males had distinctly less vibrant plumage than either vehicle or estradiol treated counterparts (**Fig. 4D-F**). To our knowledge, this is the first observation linking estrogen to zebra finch male plumage, and is consistent with a hypothesis that estrogens interact with some feather proteins in birds (Somes and Smyth, 1967) (Ralph, 1969). This finding increases our confidence that our daily dosing protocol of exemestane exerted a strong endocrine disrupting effect within the zebra finch.

**Fig. 4.**
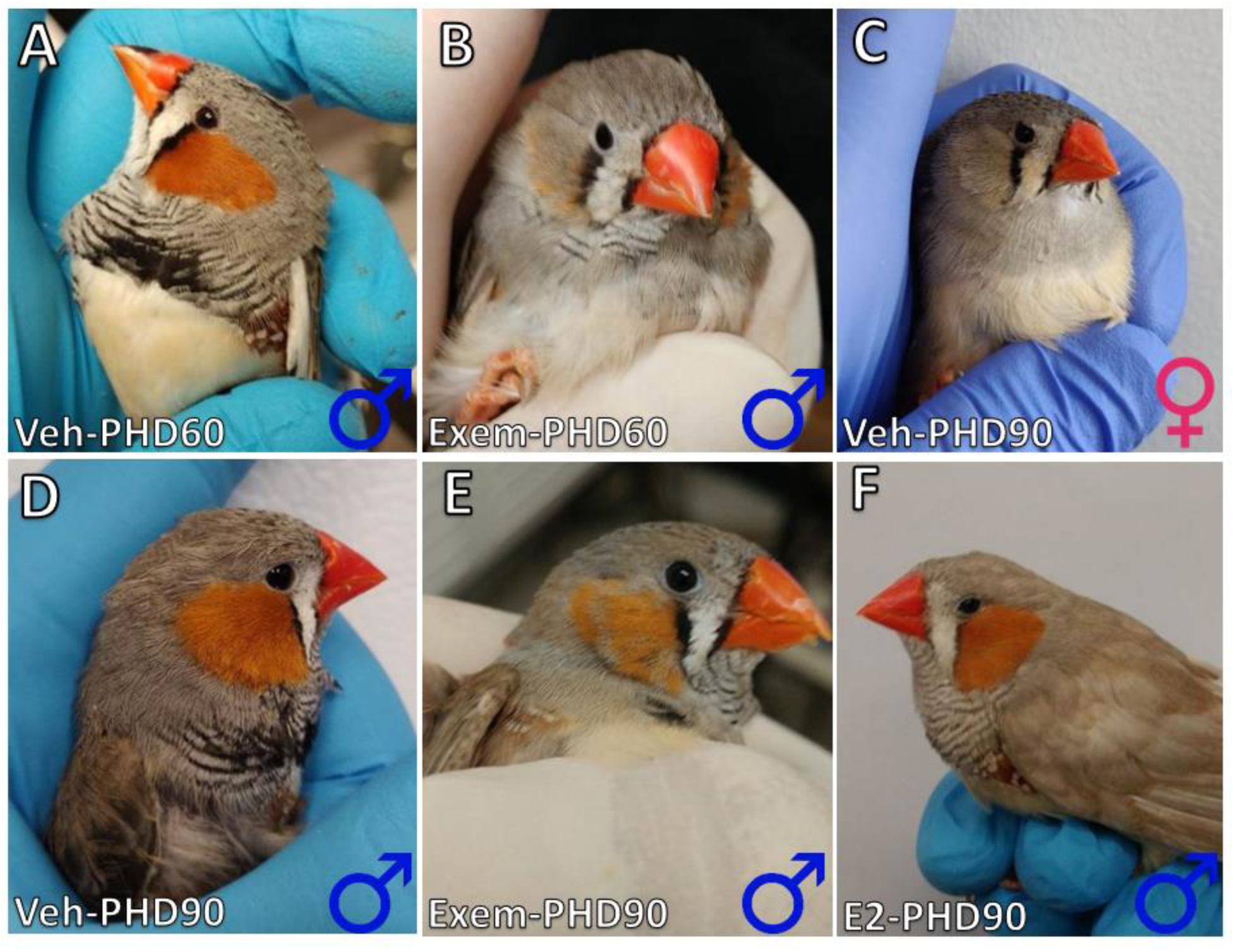
Estrogen manipulation effects on male plumage. (A) PHD60 vehicle treated male following his first molt with normal plumage, and left-over black juvenile coloration on the beak. (B) PHD60 exemestane treated male showing weak male plumage. (C) PHD90 vehicle treated female. (D) PHD90 vehicle treated male. (E) PHD90 exemestane treated male, still showing weaker male plumage. (F) Estradiol treated male, showing typical male plumage.

### 3.3 Estrogen is required for normal song development in males

Under our experimental conditions (**Fig. 1D**), males treated with vehicle over their 90-day development period produced normal songs with stereotyped syllables arranged into repeating, well-structured motifs (**Fig. 5A**). However, males treated with exemestane produced fewer unique syllable types, and thus simpler songs (**Fig. 5B, Supplemental Audios 1-9**). The songs of males treated with estradiol were similar to vehicle treated males (**Fig. 5C**). Vehicle treated females did not produce song, but only innate calls as expected (**Fig. 5D**). Interestingly we found that some females treated with exemestane showed repeated production of 1 or 2 apparently innate syllables similar to this affect for song seen in males treated with exemestane (**Fig. 5C,E**). All females treated with estradiol produced songs that resembled males (**Fig. 5F**), which they sometimes directed to other females (**Supplemental Movie 1**).

**Fig. 5.**
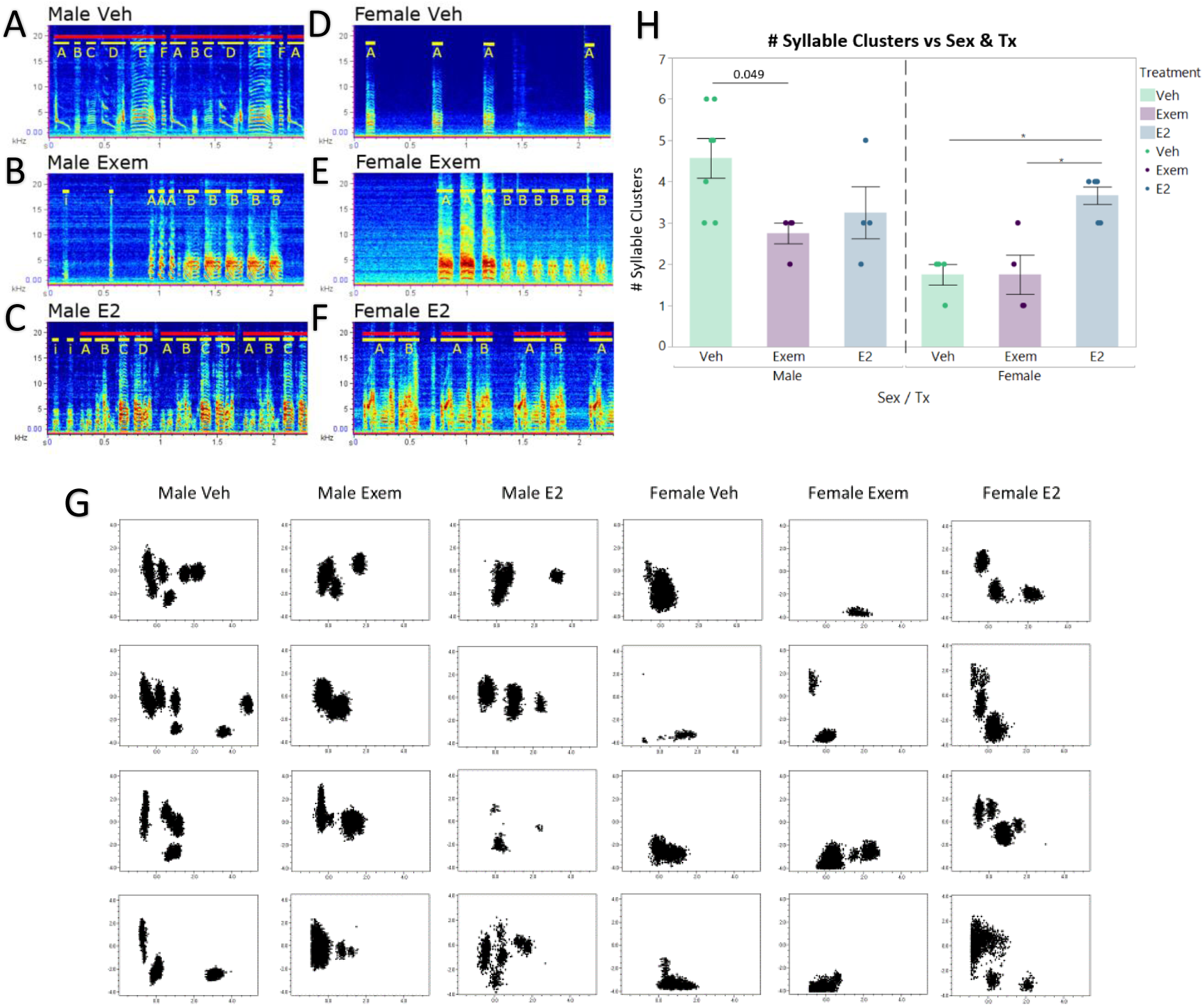
Estrogen manipulation effects on song behavior. (A) Sonogram of vehicle treated male with structured repeating motifs. Distinct syllable types are highlighted and named in yellow, and motifs are highlighted in red. (B) Exemestane treated male with simpler song. (C) Estrogen treated males with apparent normal song. (D) Vehicle treated female producing innate calls. (E) Exemestane treated females producing a series of simple syllables. (F) Estrogen treated female with song similar to that seen in males. (G) Example clusters showing song syllable repertoires of four birds for each group. Syllables clustered based on k-nearest neighbor hierarchical clustering within SAP2011, plotted by duration x frequency modulation, and converted to a binary image. (H) Cluster number was compared between sex and treatment. 2-way ANOVA and Tukey’s HSD post hoc. Sex: df=1, SS=7.041667, F=8.5932, p=0.0089 Treatment: df=2, SS=6.58333, F=4.0169, p=0.0361 Sex*Treatment: df=2, SS=10.583333, F=6.4576, p=0.0077

To quantify these observations, we performed nearest neighbor hierarchical clustering of syllables (built into Sound Analysis Pro 2011) using different acoustic features. Based on these clusters, we found that males chronically treated with exemestane (n = 4) had 40% fewer unique syllable types compared to males treated with vehicle (n = 7) (**Fig. 5G,H**). Males treated with estradiol (n = 4) had no significant difference with vehicle treated males. Estradiol treated females (n = 6) produced songs with comparable unique syllable types as vehicle or estradiol treated males (**Fig. 5G,H**). The number of syllables produced by exemestane treated females (n = 4) was not significantly different from that produced by vehicle treated females (n = 4). Sonograms and cluster plots for all birds are available in **Supplemental Fig. 3**. These findings suggest that estrogen over the entire post hatch development period is not required for males to acquire the ability to sing, but it is necessary for males to sing normally. Our findings also validate previous conclusions, that increased estrogen is sufficient for females to develop the ability to sing.

### 3.4 Estrogen induces early changes in specialized gene expression and size of song nuclei

Differences in behavior presumably reflect differences in the brain. To examine the brain, we focused on PHD30 animals (**Fig. 1C**), as this is the beginning of the sensorimotor learning stage when song nuclei have established their connections and are beginning to be utilized to produce sub-song (Mooney and Rao, 1994). Also at this stage, clear differences in song nuclei between males and females start to become apparent (Bottjer et al., 1985). We used cresyl violet staining to identify song nuclei. Area X was difficult to identify by cresyl violet at this age, and thus we used *in-situ* hybridization for mRNA of the Calcium Dependent Secretion Activator 2 (*CADPS2*), which identifies Area X and HVC in adults (Jarvis et al., 2013).

We found that in PHD30 vehicle treated controls, there was clear *CADPS2* signal in male Area X and HVC, but only in female HVC (**Fig. 6A**), supporting the proposal that Area X may not be present in females at this age (Garcia-Calero and Scharff, 2013). Treatment with exemestane did not change this pattern. In contrast, treatment of females with estradiol resulted in the appearance of an Area X with specialized *CADP2* gene expression, similar to males (**Fig. 6A**). Adjacent cresyl violet stained sections support these results (**Fig. 6B**), where in all males, regardless of treatment, we could identify pallial song nuclei (HVC, RA, and LMAN) and Area X, but among females we could identify an Area X-like region only in estradiol-treated animals.

**Fig. 6.**
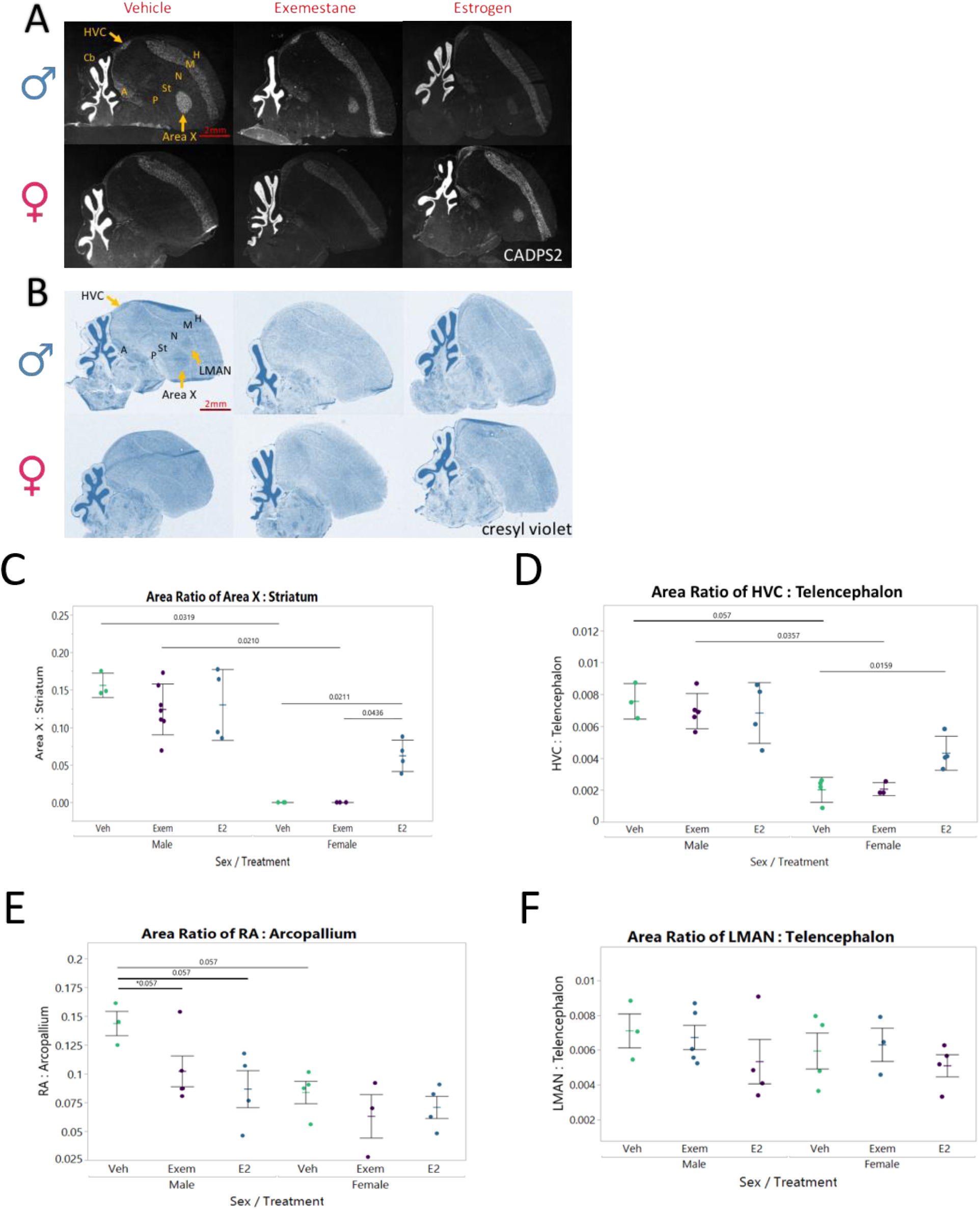
Effects of estrogen manipulation on *CADPS2* expression and sizes of song nuclei. (A) *CADPS2* mRNA expression (white) in the brains of example animals from each treatment group. Anterior is right, dorsal is up. (B) Adjacent cresyl violet stained sections to A. (C-F) Relative song nuclei sizes for HVC, Area X, RA, and LMAN respectively. Aligned ranks transformation ANOVA with post-hoc Wilcoxon rank sum. Statistics in **Supplemental Table 4**.

For HVC, RA and Area X, we noted clear visible size differences between PHD30 males and females. We quantified the areas of these song nuclei relative to individual brain subdivisions or to the whole telencephalon in the same sections, using *CADPS2* signal and cresyl violet staining to identify nuclear/regional boundaries. In vehicle controls, besides no detectable Area X in females (**Fig. 6C**), HVC and RA was larger in males (**Fig. 6D,E**), but LMAN was the same size in both sexes (**Fig. 6F; Supplemental Fig. 1**). Exemestane had no effect on male Area X, male or female HVC sizes, but did result in a trend for decrease in male RA size that became near significant after an outlier was removed (**Fig. 6E**). In contrast, estradiol treatment also caused a small decrease in male RA size (**Fig. 6E**); but in females, it caused a notably enlarged HVC (**Fig. 6D**) and the appearance of Area X (**Fig. 6C**) with sizes that approached those of males. There were no effects of sex or treatment on the sizes of the striatum, mesopallium or arcopallium relative to the telencephalon (**Supplemental Fig. 4**), indicating that the differences seen were specific to the song nuclei and not their respective brain subdivisions.

### 3.5 Specialized transcriptomes of song nuclei are differentially influenced by sex and hormones

The above findings revealed that one gene, *CADPS2*, which is specialized in juvenile male Area X is also impacted by estrogen manipulation in juvenile females. However, many genes in adults have specialized expression in different song nuclei (Lovell et al., 2008; Lovell et al., 2018; Olson et al., 2015; Pfenning et al., 2014). Prior work from our lab has shown that *SLIT1*, an axon guidance ligand, has specialized down regulation in RA beginning at PHD20-35 in males and females, but its receptor ROBO1 has specialized upregulation in RA around PHD35, followed by down regulation in adulthood in males with no changes in females (Wang et al., 2015). We wondered if exemestane or estradiol treatments impacted many genes, and whether there were differences between males and females, and among song nuclei. We therefore laser micro-dissected HVC, RA, LMAN, Area X (or the region where Area X would be in vehicle and exemestane treated females) and their immediate surrounding areas (**Supplemental Fig. 1**), and then performed RNA-seq transcriptomic profiling, with three PHD30 animals per group (a total of 144 samples; **Methods**).

PCA of expression of all 18,618 annotated genes assessed (16,882 expressed with 100 or more reads in at least one brain region), and that overlapped in the Kallisto and STAR mapped alignments, showed that samples from all 8 brain regions first clustered closely into 3 main groups that reflected their specific brain subdivision (striatum, nidopallium, and arcopallium), and then by song nucleus and surround (**Fig. 7A**). A Euclidian distance matrix plot of gene expression in all 144 samples supported this conclusion, but with LMAN showing more specialized grouping outside of the nidopallium (**Supplemental Fig. 5**). Area X showed the biggest separation by sex and treatment, where samples from males and estradiol treated females separated from all surrounding MSt samples and samples where Area X would have been located in vehicle and exemestane treated female (**Fig. 7B; Supplemental Fig. 5**). This is also consistent with high Spearman correlation values for ratios of all genes in each song nucleus versus their surround for each bird (ρ = ∼0.7-0.9; **Supplemental Table 2**), except for the Area X-analogous samples from vehicle and exemestane treated females (ρ = ∼0.1-0.5); the low correlations of the later are presumably due to low expression ratios of adjacent parts of the striatum that are not song nuclei.

**Fig. 7.**
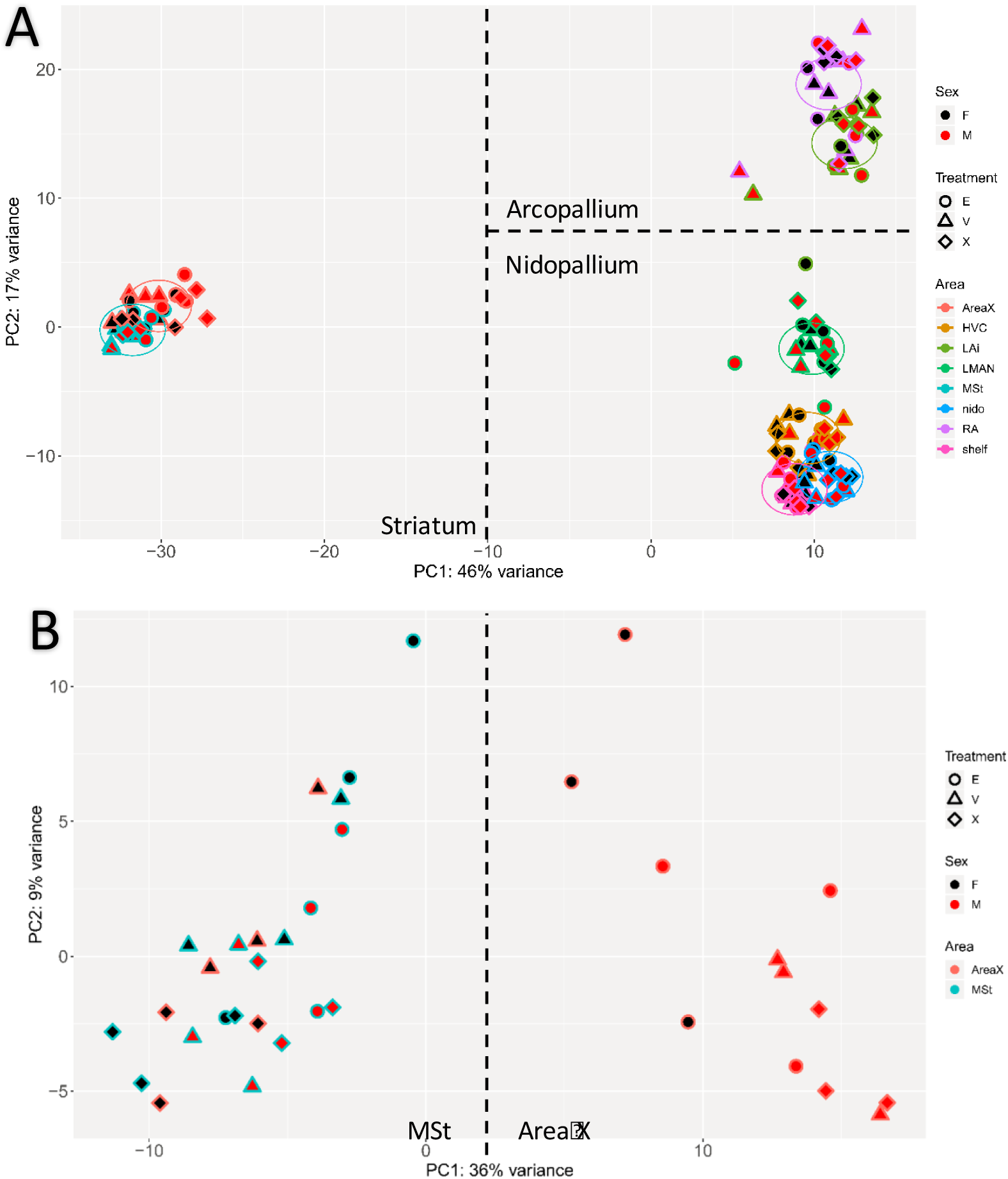
Differential gene expression in song nuclei in response to estrogen manipulation. (A) PCA plot of all RNA-Seq samples. Sex, fill color; Treatment, symbol shape; Brain region, symbol outline color. (B) PCA plot of all MSt and Area X RNA-seq samples. Female estradiol Area X samples (black filled salmon circles) cluster more closely with all male Area X samples to the right of the plot.

Unpaired statistical analysis for differentially expressed genes (DEG) over- or under-expressed in each song nucleus relative to their surrounding brain subdivision revealed that, as in adults (Pfenning et al., 2014), vehicle treated males at PHD30 already had hundreds of such DEGs (**Fig. 8, purple;** heatmap of individuals in **Supplemental Fig. 6**); overlap with DEGs in adults will be the subject of a separate study. This included *CADPS2*, which was among the top 3 genes with the greatest magnitude of specialization in Area X (**Fig. 9A-F**). Vehicle control females had no detectable DEGs of those seen in vehicle control males (**Fig. 8A**) or of any other genes (**Fig. 9B**) in the region where Area X would normally be located; the small and still present HVC had some DEGs (11 genes), but far fewer at FDR <0.5 than control males (818 genes; **Fig. 8B**); in RA, there were fewer differences between males (202 genes) and females (437 genes; **Fig. 8C**); and in LMAN there were very few differences (1023 vs 1099, respectively, **Fig. 8D**).

**Fig. 8.**
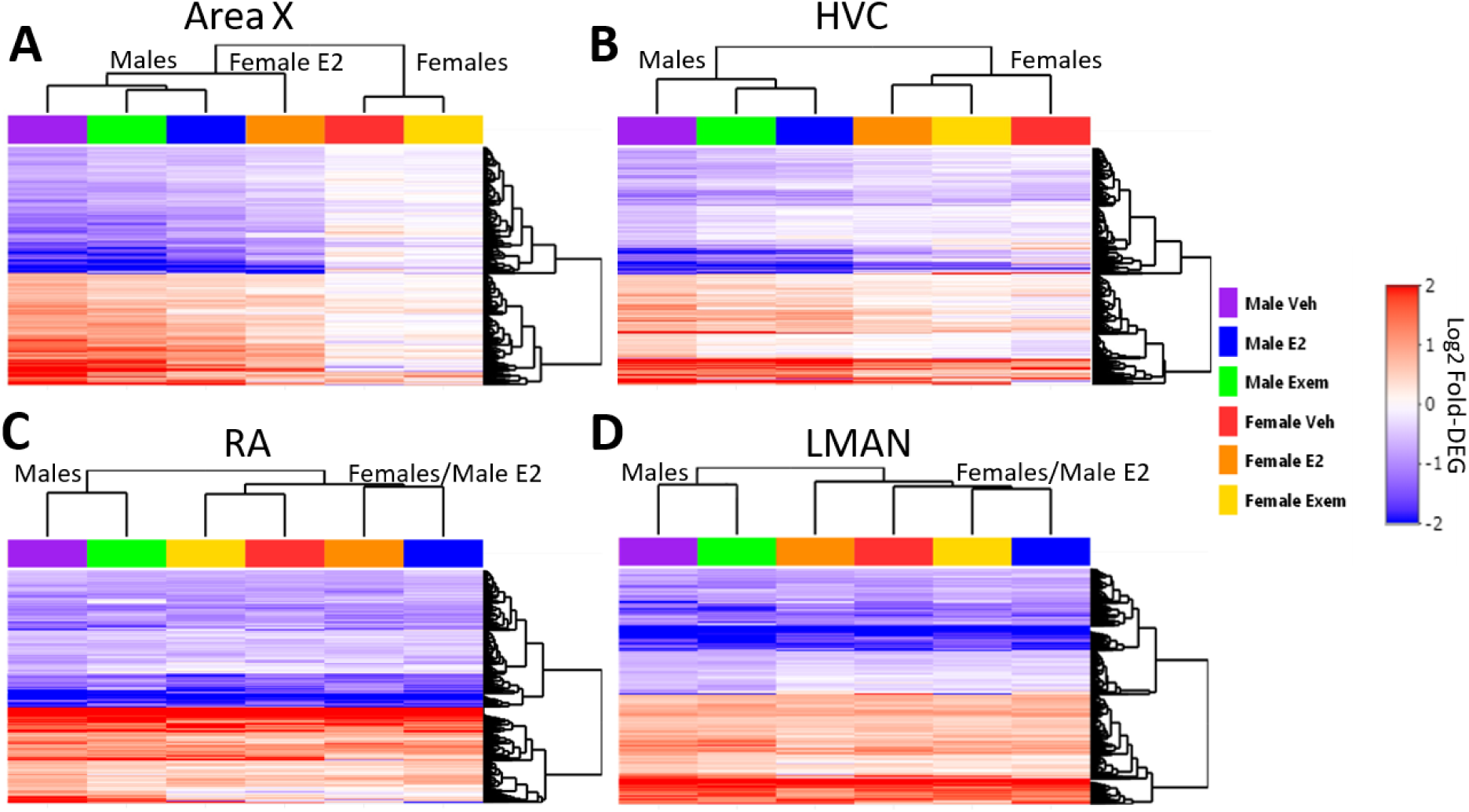
Heatmap of all song nuclei gene expression specializations for all treatment groups. Shown are genes with significant differential expression in male song nuclei of PHD30 animals, and their profiles in all other groups. (A) Vehicle male Area X had 326 DEGs. Note vehicle and exemestane treated females had hardly any specialized gene expression of the genes found in males. (B) Vehicle male HVC had 818 DEGs. Regardless of treatment, DEG patterns separated based on sex with males showing far greater fold changes for the 818 genes than females. (C) Vehicle male RA had 202 DEGs. Vehicle treated animals clustered with exemestane treated animals within sex, however estradiol treatment animals were most similar to each other and then more similar to females than males. (D) Vehicle male LMAN has 1023 DEGs. Vehicle and exemestane treated males were most similar to each other. Heatmaps show differential expression in log2 fold change (log2FC) values for each gene (row) by experimental group (column). Values with log2FC greater than |2| were binned at |2|. Red, increased expression in song nuclei relative to the surround. Blue, decreased expression.

**Fig. 9.**
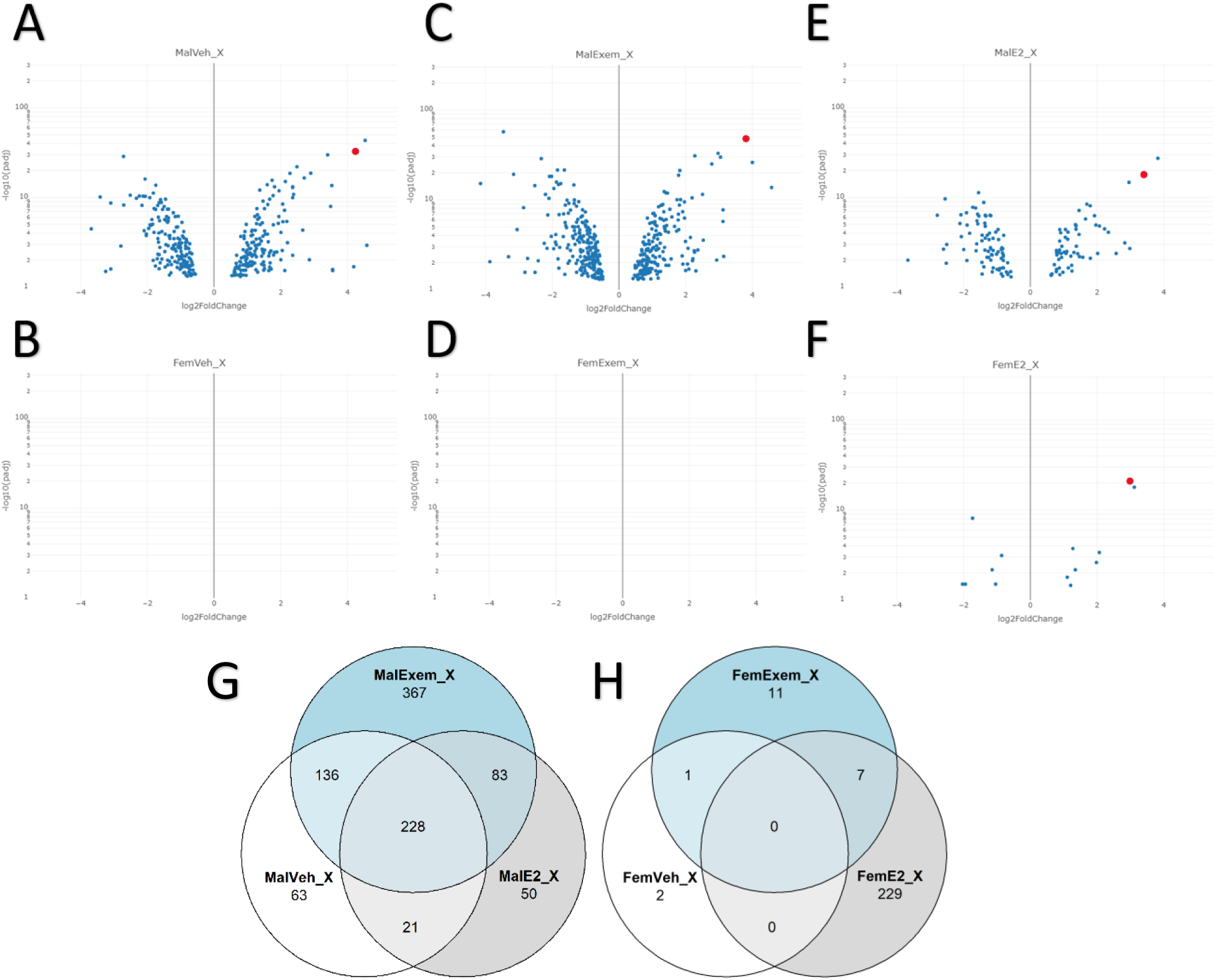
Fold changes and overlaps in DEGs between groups for Area X. (A-F) Volcano plots of differentially expressed genes in Area X as compared to MSt from male and female finches treated with vehicle, estradiol, or exemestane. Only DEGs with FDR <0.05 were plotted. *CADPS2* is colored in red. X-axis is log2 fold change values with MSt enriched genes appearing to the left, and Area X enriched genes appearing to the right. Y-axis is the -log10 transformed FDR values on a log10 scale. (G) Venn diagram of pairwise DEGs in Area X in males between treatment groups. (H) Venn diagram of pairwise DEGs in Area X in females between treatment groups. Venn diagrams for all other song nuclei are shown in **Supplemental Fig. 9**.

Exemestane treatment had little impact on the specialized gene expression in male song nuclei (**Fig. 8, 9C**), indicating that blocking estrogen did not demasculinize male song nuclei gene expression. There were also no notable large specialized transcriptome changes in females treated with exemestane (**Fig. 8, 9D**). Estradiol treatment, however, resulted in a dramatic change of female Area X to have a DEG profile similar to males, albeit at lower magnitude of differential expression for most genes (**Fig. 8A, orange**) with only the highest magnitude differences passing an unpaired FDR < 0.05 (**Fig. 9F**). This conclusion was supported by a more sensitive pairwise statistical analyses of song nuclei and surround for each bird, which revealed many more of these genes in females that pass a paired FDR <0.05 (**Supplemental Fig. 7A-F**). HVC showed the next most notable differences, where unlike Area X, treating females with estradiol only marginally made female HVC to be more male-like in its specialized gene expression profile compared to vehicle control females (**Fig. 8B**). In RA and LMAN, although they showed much fewer differences between control males and control females, estradiol induced male specializations to be more female-like (**Fig. 8C,D**). Full genelist of overlapping and unique DEG for each nucleus and group are available in **Supplemental Table 3**.

We evaluated a subset of genes by *in-situ* hybridization (36 genes across 4 song nuclei) on PHD30 animals performed in our lab, or analyzed from the online zebra finch gene expression atlas on control adult males (http://www.zebrafinchatlas.org) and our past adult studies (Jarvis et al., 2013). We found that 100/140 song nuclei patterns (71%) of the top DEGs examined from our RNA-seq data were congruent with the *in-situ* hybridization data (**Table 1**). This validation rate is presumably lower than what it really is, as we know differences exist between DEGs in juveniles (this study) and adults (past studies). Overall, these findings indicate that the molecular specializations that define each song learning nucleus are on different developmental trajectories in their growth in males and atrophy in females. This trajectory in females is altered in response to estrogen, but long-term blockage of estrogen synthesis minimally changes these specializations in males.

**Table 1.** DEGs with localization data from the zebra finch gene expression atlas and other sources.

| Gene Name | Area X |  | HVC |  | RA |  | LMAN |  |
| --- | --- | --- | --- | --- | --- | --- | --- | --- |
|  | RNA-Seq | ISH Atlas | RNA-Seq | ISH Atlas | RNA-Seq | ISH Atlas | RNA-Seq | ISH Atlas |
| KCNT2 | UP | ✓ | - | ✓ | - | ✓ | - | ✓ |
| TAC1 | UP | ✓ | DOWN | ✗ | - | ✓ | DOWN | ✓ |
| FLRT2 | UP | ✓ | UP | ✓ | UP | ✗ | - | ✗ |
| CACNG3 | DOWN | ✓ | DOWN | ✗ | - | ✓ | DOWN | ✓ |
| ALDH1A2 | - | ✓ | UP | ✓ | UP | ✗ | UP | ✓ |
| PVALB | - | ✓ | UP | ✓ | UP | ✓ | UP | ✓ |
| IGF1 | UP | ✗ | - | ✓ | UP | ✓ | DOWN | ✗ |
| ADCYAP1 | - | ✓ | DOWN | ✓ | DOWN | ✓ | - | ✓ |
| GRIN3A | DOWN | ✗ | - | ✓ | DOWN | ✓ | DOWN | ✓ |
| CLMN | - | ✗ | - | ✓ | DOWN | ✗ | - | ✓ |
| CBLN2 | UP | ✗ | UP | ✓ | - | ✓ | UP | ✓ |
| SEMA3E | UP | ✓ | UP | ✓ | - | ✓ | UP | ✓ |
| ADRA2A | - | ✓ | - | ✓ | UP | ✗ | UP | ✓ |
| SCN4B | - | ✓ | UP | ✓ | - | ✓ | UP | ✓ |
| PLXNC1 | - | ✓ | DOWN | ✓ | - | ✗ | DOWN | ✓ |
| HPCAL1 | - | ✗ | DOWN | ✓ | - | ✓ | DOWN | ✓ |
| SLC6A7 | - | ✓ | - | ✓ | - | ✗ | DOWN | ✗ |
| CCK | - | ✓ | - | ✓ | DOWN | ✗ | DOWN | ✓ |
| CACNG4 | DOWN | ✗ | DOWN | ✗ | - | ✓ | DOWN | ✗ |
| CPE | - | ✓ | - | ✓ | - | ✓ | DOWN | ✓ |
| SEMA3A | DOWN | ✗ | DOWN | ✗ | - | ✓ | DOWN | ✗ |
| DACH1 | - | ✓ | DOWN | ✗ | - | ✓ | DOWN | ✗ |
| NR0B1 | - | ✓ | UP | ✓ | - | ✓ | UP | ✗ |
| ROBO2 | - | ✗ | UP | ✓ | - | ✓ | - | ✓ |
| ZEB2 | - | ✓ | UP | ✓ | - | ✓ | DOWN | ✗ |
| FOSL2 | - | ✓ | UP | ✓ | - | ✓ | - | ✓ |
| POMC | - | ✓ | UP | ✗ | - | ✓ | - | ✓ |
| VIP | DOWN | ✓ | UP | ✗ | DOWN | ✓ | - | ✓ |
| GRIA4 | DOWN | ✗ | DOWN | ✓ | - | ✓ | DOWN | ✗ |
| HPCAL1 | - | ✗ | DOWN | ✗ | - | ✓ | DOWN | ✓ |
| ADCYAP1 | - | ✓ | DOWN | ✓ | DOWN | ✓ | - | ✓ |
| GLRA4 | - | ✗ | DOWN | ✓ | - | ✓ | DOWN | ✗ |
| AMIGO2 | DOWN | ✗ | DOWN | ✗ | - | ✓ | DOWN | ✗ |
| MAPK1 | - | ✓ | DOWN | ✗ | - | ✓ | - | ✓ |
| AKR1D1 | - | ✓ | DOWN | ✓ | - | ✓ | - | ✓ |

RNA-seq differential analysis data (left subcolumn) from post hatch day 30 vehicle/control males compared to RNA *in-situ hybridization* data (right subcolumn) from adult control males. Congruent DEGs are highlighted in blue and incongruent DEGs are highlighted in salmon.

**Table 2.**
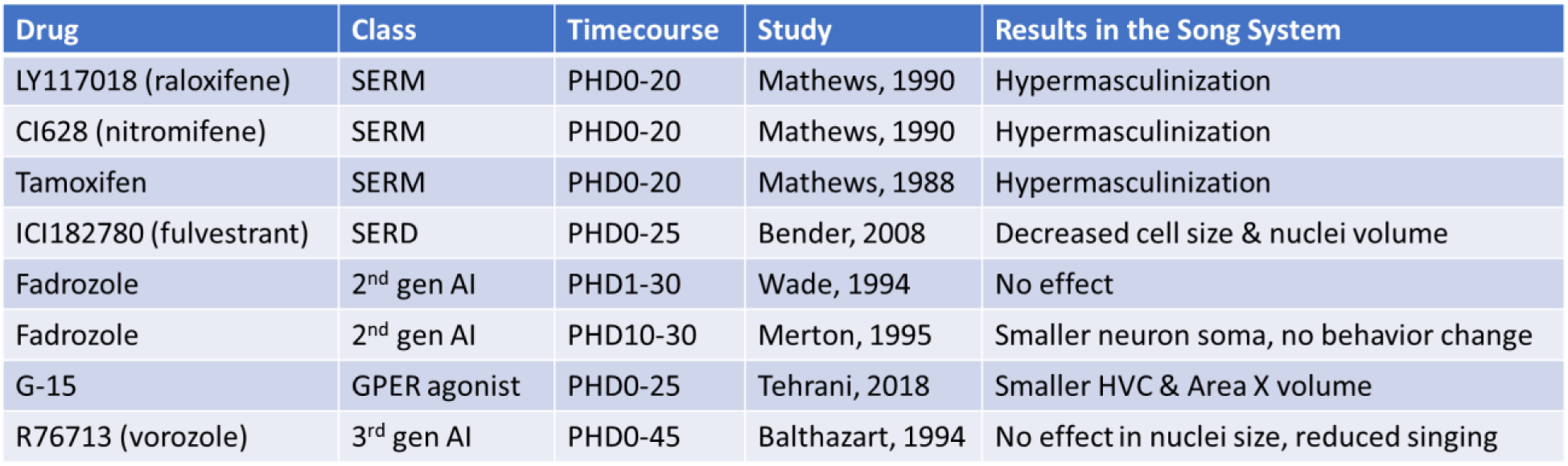
Summary of prior studies modulating estrogen in zebra finches

### 3.6 Each song learning nucleus has specific functional molecular specializations

The DEGs we found for each song nucleus suggest that they contribute to differences in song nuclei functions (Pfenning et al., 2014; Whitney et al., 2014). We performed gene ontology (GO) analyses on the specialized DEGs identified in the unpaired test from control males at PHD30, and found that the top biological categories in Area X were associated with synapse transmission and signaling, neurotransmitter release, and neuron development (**Fig. 10A**); HVC and LMAN were specialized for neuron development, synaptic development and transynaptic signaling (**Fig. 10B,D**); and RA was specialized also for neurotransmitter release, and synaptic transmission and signaling (**Fig. 10C**). In terms of specific molecular function specializations, top GO terms were ion and cation channels in Area X, HVC, and RA, but receptor-ligand activity in RA (**Fig. 10**). Additional genes with lower magnitude differences in the paired analyses did not change these results for the top GO categories, except channel activity was more specialized than receptor-ligand activity for RA (**Supplemental Fig. 8**).

**Fig. 10.**
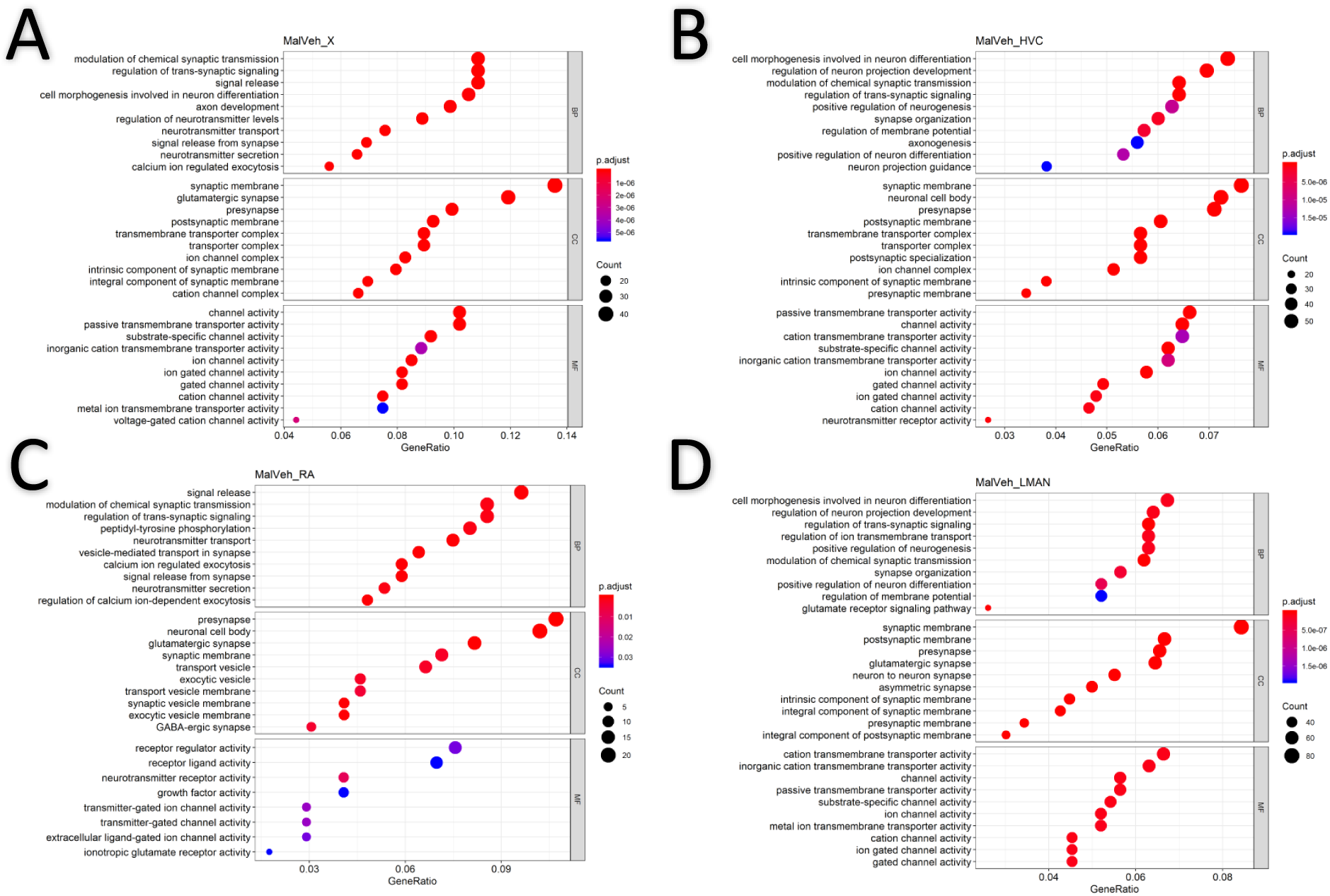
Top GO terms for DEGs in each song nucleus in vehicle males. (A-D) Area X, HVC, RA, and LMAN respectively. X-axis indicates the ratio of genes with specialized expression out of the total list of genes, which contribute to each value. Count, number of DEGs that contributed to each category. BP, Biological Process; CC, Cell Compartment; MF, Molecular Function. Genelist is from all vehicle male DEGs for each region with FDR <0.05. GO enrichment results include p.adjusted value <0.1. Heatmap scale, level of significance.

**Fig. 11.**
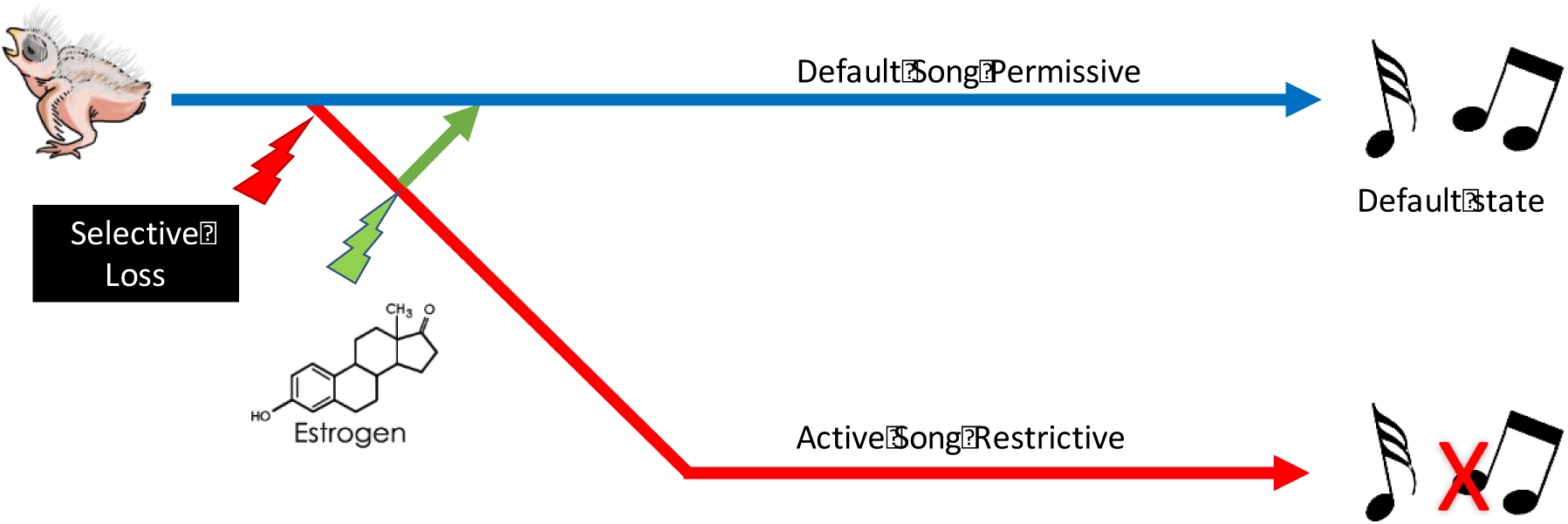
Alternative hypothesis of female song loss in sexually dimorphic songbirds. The song permissive state is the ancestral trait of songbirds (blue line), the rapid degeneration of the W chromosome either alone or in tandem with rapid evolution of the Z chromosome may contribute to the active restriction of the development of the song system in sexually dimorphic songbirds (red line). Estrogen may work downstream of this induced atrophy to rescue the nascent song system (green line).

We next examined GO enrichment of genes that changed in response to estrogen or exemestane treatment. To do this, we examined genes that were uniquely differentially expressed in estradiol treated females when compared to vehicle or exemestane treated females, and exemestane treated males when compared to vehicle or estradiol treated males, for each song nucleus, from the paired statistical analyses to obtain more of the lower magnitude differences (**Supplemental Fig. 9**). Top biological GO terms in estradiol treated females for Area X included ion transport, ATP metabolic processes, and respiratory/oxidation processes; HVC and RA included cellular differentiation, axonogenesis, and development; and LMAN included synapse organization/regulation. For exemestane treated males, top biological GO terms for Area X DEGs included signaling, action potential regulation/processes, and ion transport; and RA had terms for cellular respiration (**Supplemental Fig. 10**). Overall, the GO analyses indicate that the varied specializations may lead diverse functional outcomes, among the different song nuclei, in the different sexes, in response to estrogen modulation.

## 4. Discussion

Through dose-controlled treatment with exemestane or estradiol, we were able to examine the effects of chronic estrogen manipulation on song behavior and the transcriptome of the song system during the onset of the sensorimotor vocal learning period in zebra finches. In males, we found an unexpected interaction between estrogen and male specific plumage development. We also found that male song learning was diminished by estrogen depletion, although there was little impact on the gross anatomy of the song learning circuit and its molecular specializations. In contrast, in females, we recapitulated prior studies establishing the need for artificially high levels of estrogen for female song learning and discovered that the development of the different song learning nuclei have different responses to estrogen manipulation. Notably Area X and its molecular specializations were the most dependent on high levels of estrogen in females at the start of the sensorimotor learning period. These findings support the hypothesis that vocal learning loss in female zebra finches evolved alongside a sex specific dependence on estrogen for vocal learning, suggesting that vocal learning evolved in both sexes first independent of estrogen and was later lost in the females. This behavioral loss is associated with an uneven neural loss of the different song nuclei. We suggest that Area X is the first to atrophy or not even appear, followed by HVC, then RA, and finally LMAN with limited atrophy.

Our results with estrogen inhibition agree with some past publications but not others (**Table 4**). Some of the differences could be due to the pharmacological agent we used. Exemestane is more specific for aromatase inhibition with limited extra-target effects and without the “rebound” effects seen with other aromatase inhibitors (Harada and Hatano, 1998; Lonning and Eikesdal, 2013). Other differences may be due to our delivering the agents chronically until the time of sacrifice. The requirement of estrogen for sexually dimorphic male plumage was unexpected for us, and not seen in prior studies that we are aware of. The highly reduced male plumage in the presence of still well-developed song learning nuclei with generally preserved gene expression specializations in exemestane treated animals suggest that estrogenic influence on these two anatomical systems may be independent.

However, our findings suggest that estrogen still plays a modulatory role in song learning, since all males dosed with exemestane had impoverished singing ability. We were also surprised to see that chronic exemestane treatment in females caused some females to produce 1-2 repeated syllables in a song-like manner, even though they had atrophied song nuclei. Prior studies have shown that acute aromatase inhibition can suppress the rate of singing in zebra finches and canaries (Alward et al., 2016; Vahaba et al., 2019; Walters and Harding, 1988), but no changes in song learning acuity have been reported (Merten and Stocker-Buschina, 1995; Vahaba et al., 2019). Perhaps the more potent and long-term application of the estrogen inhibitor we used revealed an unknown involvement of estrogen in vocal learning. This could be through estrogenic modulation of a subset of genes with specialized expression in the song nuclei detected here that change with exemestane treatment in males. Alternatively, the mechanism could be through the auditory pathway, as recent work has shown that acute aromatase inhibition with fadrozole alters hearing-induced expression of activity-dependent genes in the zebra finch auditory pallium, and more so in males than in females (Krentzel et al., 2019).

Although the estrogen-induced “masculinization” of the female song system and behavior we observed is considerable, it is still incomplete with regards to song nuclei size and molecular specializations. This indicates that there may be other factors at play besides estrogen in suppressing the development of the song learning system in females, or our estrogen administration protocol was not optimal, which is a possibility as we had to use far less estradiol (200ug pellets at PHD20 instead of 1500ug) than was published previously (Simpson and Vicario, 1991a; Simpson and Vicario, 1991b). This was due to the toxic effects of long-term elevated estrogen on bone integrity in birds (Whitehead, 2004). The finding that most HVC gene expression specializations are not sensitive to estrogen treatment in females, whereas specializations in Area X are extremely sensitive, was unexpected as HVC is the only song nucleus known to express the classical NR (nuclear receptor) ESR1 (estrogen receptor α), and this ESR1 expression is not yet sexually dimorphic at ∼PHD30 (Gahr, 1996; Gahr and Konishi, 1988; Jacobs et al., 1999). Additionally, an intact HVC is necessary for RA development in males and estrogen-treated females, and for Area X in estrogen-treated females (Akutagawa and Konishi, 1994; Herrmann and Arnold, 1991). Combining findings, one interpretation could be that the ESR1 receptor in HVC may lead to specialization of some genes in HVC that influences a cue for Area X development, but only in females. The higher degree of stability of specialized gene expression in RA and LMAN regardless of sex or estrogen manipulation suggest that their specialized functions during development are not strongly estrogen dependent. Additionally, there are other estrogen receptors, ESR2 (Estrogen receptor β) (Kuiper et al., 1996; Mosselman et al., 1996) and GPER (Maggiolini and Picard, 2010) that maybe involved. Selective receptor agonist/antagonist studies will need to be conducted in the future to determine their individual effects.

Our findings help shed light on the hypotheses of the origin of vocal learning systems in songbirds and their sex differences. The song learning system of songbirds is inferred to have evolved ∼30 million years ago with the emergence of the split between oscine and suboscine (songbird) Passeriformes (Jarvis et al., 2014). For many years it was assumed that sex differences in song learning could be an ancestral trait, consistent with the findings that many songbird species use their learned vocalizations for sexually dimorphic behaviors such as mate attraction and territorial defense. This begged the question as to why the female song system appeared to be estrogen dependent, while the male song system was estrogen independent. A fact underappreciated by many scientists in temperate regions, is that in the vast majority of songbirds, both males and females, sing, and they are concentrated more in equatorial regions (Jarvis, 2004). An analysis of many songbird species globally and phylogenetically suggest that female-selective loss occurred multiple independent times among songbirds (Odom et al., 2014), indicating that there may be positive selection for song repression in females rather than a gain/expansion only in males. One explanation for the geographic difference in sex-specific song learning, is that after songbird song learning evolved ∼30 million years ago, species in temperate zones faced a more demanding environment that selected for more division of labor between the sexes and/or increased competition for limited resources (Jarvis, 2004). Here we further propose that sex specific division of labor for vocal learning in zebra finches was selected for by a sex hormone-dependent mechanism in females. This would also allow for evolution to reverse this loss in females should environmental factors reselect for it. These hypotheses can be further tested in future studies using species with and without vocal learning sexual dimorphism.

Our findings are also informative for understanding variations in sex determination on behavior. In mammals, females are homogametic and generally regarded as the default sex. In mammals, the genetic signals from the male Y chromosome induce development of male reproductive organs and the second X chromosome in females become silenced to prevent overdosage of X chromosomal genes. In birds, males are the homogametic sex, carrying two Z chromosomes. Either signals from reduced Z chromosome dosage or the female specific W chromosome may influence development of female reproductive organs. Whether the brain has a default sex state in birds or mammals is still open to debate, but it is well known that estrogen plays an organizational role in the sexual differentiation of the central nervous system (McCarthy, 2008). Sexually dimorphic structures within the hypothalamus can be found in all vertebrates, indicating that these structures are ancient, present long before the synapsid/sauropsid split >300 million years ago (Bruce, 2009; Godwin and Crews, 1997; Kumar and Hedges, 1998). The sexually dimorphic hypothalamic medial preoptic nucleus in chicken and quail are sensitive to estrogen modulation during pre-hatching in a manner that is opposite to that seen in mammals, where estrogen administration feminizes and aromatase inhibition masculinizes this nucleus and associated behaviors (Balthazart and Ball, 1995; Balthazart et al., 1992; Kurian et al., 2010; Lambeth et al., 2016; McEwen et al., 1977; Panzica et al., 2001; Panzica et al., 1998). Further, in many amniotic vertebrate systems, estrogenic activity during an early critical period sets up the bipotential brain to develop in a masculine or feminine manner (Juntti et al., 2010; Kurian et al., 2010). Taking all this into consideration, we propose that vocal learning in songbirds is the default state, not linked to sex chromosome determination genes nor ancient brain pathways that exist in a sex-specific state (**Fig. 12**, blue). After vocal learning evolved, we suggest that selective loss/suppression of vocal learning in females of some species became linked to sex determination genes (**Fig. 12**, red), which can then be overturned or otherwise modulated through the actions of estrogen (**Fig.12**, green).

If this hypothesis is correct, it would mean that the molecular specializations of the vocal learning systems, particularly Area X in the striatum, became subsequently linked to the estrogen regulatory pathway and the W chromosome in females. Finding that link may help towards understanding the mechanism of specialized gene regulation in vocal learning systems in songbirds, but also the convergently evolved specializations with human speech brain regions (Pfenning et al., 2014). Although humans have very small sex differences in vocal learning and spoken language behaviors (Ross et al., 2015; Weis et al., 2019) relative to sexually dimorphic vocal learning birds, some language deficits and other learning deficits are strongly linked to sex (Ferri et al., 2018; Werling and Geschwind, 2013), and a subset of the specialized genes in songbirds is convergent with those found in human spoken language brain regions (Pfenning et al., 2014). Future comparative studies in genomics, behavior, and sex hormone regulation in humans as well as other vocal learning species should help shed light on these hypotheses.

## Supporting information

Supplemental Audios 1-9

## Acknowledgements

We thank: Matthew Biegler and Lindsey Cantin for optimizing the *in situ hybridization* protocol for use in zebra finch sections. Gregory Gedman and Matthew Davenport for their instruction on laser capture microscopy. Bettina Haase and Olivier Fedrigo for guidance during RNA-Seq library preparation; Gregory Gedman and James Cahill for helpful discussion regarding transcriptomic analysis. Claire de March for her guidance on *in silico* protein-modeling. We also thank Robyn Donahue for animal husbandry, and members of the Duke metabolomics core for the mass spec steroid assays: Lisa St. John-Williams, Will Thompson, and Arthur Moseley. Lastly, we thank Claire de March, Aashutosh Vihani, Tatjana Abaffy, and Veronica Choe for helpful feedback on the manuscript.

## References

ZEBrA. Oregon Health & Science University, Portland, OR 97239.

Adam, I., Scharff, C., Honarmand, M., 2014. Who is who? Non-invasive methods to individually sex and mark altricial chicks. J Vis Exp, 51429.

Akutagawa, E., Konishi, M., 1994. Two separate areas of the brain differentially guide the development of a song control nucleus in the zebra finch. Proceedings of the National Academy of Sciences 91, 12413.

Alward, B.A., de Bournonville, C., Chan, T.T., Balthazart, J., Cornil, C.A., Ball, G.F., 2016. Aromatase inhibition rapidly affects in a reversible manner distinct features of birdsong. Scientific Reports 6, 32344.

Balthazart, J., Ball, G.F., 1995. Sexual differentiation of brain and behavior in birds. Trends Endocrinol Metab 6, 21–29.

Balthazart, J., De Clerck, A., Foidart, A., 1992. Behavioral demasculinization of female quail is induced by estrogens: Studies with the new aromatase inhibitor, R76713. Hormones and Behavior 26, 179–203.

Bender, A.T., Veney, S.L., 2008. Treatment with the specific estrogen receptor antagonist ICI 182,780 demasculinizes neuron soma size in the developing zebra finch brain. Brain Res 1246, 47–53.

Bottjer, S.W., Glaessner, S.L., Arnold, A.P., 1985. Ontogeny of brain nuclei controlling song learning and behavior in zebra finches. The Journal of Neuroscience 5, 1556.

Bray, N.L., Pimentel, H., Melsted, P., Pachter, L., 2016. Near-optimal probabilistic RNA-seq quantification. Nature Biotechnology 34, 525–527.

Breed, M.D., Moore, J., 2010. Encyclopedia of animal behavior. Elsevier B.V., Amsterdam.

Bruce, L.L., 2009. Evolution of the Hypothalamus in Amniotes, in: Binder, M.D., Hirokawa, N., Windhorst, U. (Eds.), Encyclopedia of Neuroscience. Springer Berlin Heidelberg, Berlin, Heidelberg, pp. 1363–1367.

Carrer, H.F., Cambiasso, M.J., 2009. Sexual Differentiation of the Brain: Genetic, Hormonal and Trophic Factors in: Janigro, D. (Ed.), Mammalian Brain Development 1ed. Springer, pp. 1–15.

Çiftci, H.B., 2012. Effect of estradiol-17β on follicle-stimulating hormone secretion and egg-laying performance of Japanese quail. animal 6, 1955–1960.

Dobin, A., Davis, C.A., Schlesinger, F., Drenkow, J., Zaleski, C., Jha, S., Batut, P., Chaisson, M., Gingeras, T.R., 2013. STAR: ultrafast universal RNA-seq aligner. Bioinformatics 29, 15–21.

Feenders, G., Liedvogel, M., Rivas, M.V., Zapka, M., Horita, H., Hara, E., Wada, K., Mouritsen, H., Jarvis, E.D., 2008. Molecular Mapping of Movement-Associated Areas in the Avian Brain: A Motor Theory for Vocal Learning Origin. PLoS ONE 3, e1768.

Ferri, S.L., Abel, T., Brodkin, E.S., 2018. Sex Differences in Autism Spectrum Disorder: a Review. Current Psychiatry Reports 20, 9.

Gahr, M., 1996. Developmental changes in the distribution of oestrogen receptor mRNA expressing cells in the forebrain of female, male and masculinized female zebra finches. Neuroreport 7, 2469–2473.

Gahr, M., Konishi, M., 1988. Developmental changes in estrogen-sensitive neurons in the forebrain of the zebra finch. Proc Natl Acad Sci U S A 85, 7380–7383.

Garcia-Calero, E., Scharff, C., 2013. Calbindin expression in developing striatum of zebra finches and its relation to the formation of area X. The Journal of comparative neurology 521, 326–341.

Gobes, S.M.H., Jennings, R.B., Maeda, R.K., 2017. The sensitive period for auditory-vocal learning in the zebra finch: Consequences of limited-model availability and multiple-tutor paradigms on song imitation. Behav Processes, S0376-6357(0317)30163-30168.

Godwin, J., Crews, D., 1997. Sex Differences in the Nervous System of Reptiles. Cellular and Molecular Neurobiology 17, 649–669.

Grisham, W., Arnold, A.P., 1995. A direct comparison of the masculinizing effects of testosterone, androstenedione, estrogen, and progesterone on the development of the zebra finch song system. Journal of neurobiology 26, 163–170.

Gurney, M.E., 1982. Behavioral correlates of sexual differentiation in the zebra finch song system. Brain Research 231, 153–172.

Harada, N., Hatano, O., 1998. Inhibitors of aromatase prevent degradation of the enzyme in cultured human tumour cells. British journal of cancer 77, 567–572.

Herrmann, K., Arnold, A.P., 1991. Lesions of HVc block the developmental masculinizing effects of estradiol in the female zebra finch song system. Journal of neurobiology 22, 29–39.

Holloway, C.C., Clayton, D.F., 2001. Estrogen synthesis in the male brain triggers development of the avian song control pathway *in vitro*. Nat Neurosci 4, 170–175.

Jacobs, E.C., Arnold, A.P., Campagnoni, A.T., 1999. Developmental regulation of the distribution of aromatase- and estrogen-receptor-mRNA-expressing cells in the zebra finch brain. Dev Neurosci 21, 453–472.

Jarvis, E.D., 2004. Chapter 8 - Brains and birdsong, in: Marler, P., Slabbekoorn, H. (Eds.), Nature’s Music. Academic Press, San Diego, pp. 226–271.

Jarvis, E.D., 2019. Evolution of vocal learning and spoken language. Science 366, 50.

Jarvis, E.D., Mirarab, S., Aberer, A.J., Li, B., Houde, P., Li, C., Ho, S.Y.W., Faircloth, B.C., Nabholz, B., Howard, J.T., Suh, A., Weber, C.C., da Fonseca, R.R., Li, J., Zhang, F., Li, H., Zhou, L., Narula, N., Liu, L., Ganapathy, G., Boussau, B., Bayzid, M.S., Zavidovych, V., Subramanian, S., Gabaldón, T., Capella-Gutiérrez, S., Huerta-Cepas, J., Rekepalli, B., Munch, K., Schierup, M., Lindow, B., Warren, W.C., Ray, D., Green, R.E., Bruford, M.W., Zhan, X., Dixon, A., Li, S., Li, N., Huang, Y., Derryberry, E.P., Bertelsen, M.F., Sheldon, F.H., Brumfield, R.T., Mello, C.V., Lovell, P.V., Wirthlin, M., Schneider, M.P.C., Prosdocimi, F., Samaniego, J.A., Velazquez, A.M.V., Alfaro-Núñez, A., Campos, P.F., Petersen, B., Sicheritz-Ponten, T., Pas, A., Bailey, T., Scofield, P., Bunce, M., Lambert, D.M., Zhou, Q., Perelman, P., Driskell, A.C., Shapiro, B., Xiong, Z., Zeng, Y., Liu, S., Li, Z., Liu, B., Wu, K., Xiao, J., Yinqi, X., Zheng, Q., Zhang, Y., Yang, H., Wang, J., Smeds, L., Rheindt, F.E., Braun, M., Fjeldsa, J., Orlando, L., Barker, F.K., Jønsson, K.A., Johnson, W., Koepfli, K.-P., O’Brien, S., Haussler, D., Ryder, O.A., Rahbek, C., Willerslev, E., Graves, G.R., Glenn, T.C., McCormack, J., Burt, D., Ellegren, H., Alström, P., Edwards, S.V., Stamatakis, A., Mindell, D.P., Cracraft, J., Braun, E.L., Warnow, T., Jun, W., Gilbert, M.T.P., Zhang, G., 2014. Whole-genome analyses resolve early branches in the tree of life of modern birds. Science 346, 1320.

Jarvis, E.D., Nottebohm, F., 1997. Motor-driven gene?expression. Proceedings of the National Academy of Sciences 94, 4097–4102.

Jarvis, E.D., Yu, J., Rivas, M.V., Horita, H., Feenders, G., Whitney, O., Jarvis, S.C., Jarvis, E.R., Kubikova, L., Puck, A.E.P., Siang-Bakshi, C., Martin, S., McElroy, M., Hara, E., Howard, J., Pfenning, A., Mouritsen, H., Chen, C.-C., Wada, K., 2013. Global View of the Functional Molecular Organization of the Avian Cerebrum: Mirror Images and Functional Columns. Journal of Comparative Neurology 521, 3614–3665.

Juntti, S.A., Tollkuhn, J., Wu, M.V., Fraser, E.J., Soderborg, T., Tan, S., Honda, S.-I., Harada, N., Shah, N.M., 2010. The Androgen Receptor Governs the Execution, but Not Programming, of Male Sexual and Territorial Behaviors. Neuron 66, 260–272.

Kojima, S., Doupe, A.J., 2007. Song selectivity in the pallial-basal ganglia song circuit of zebra finches raised without tutor song exposure. J Neurophysiol 98, 2099–2109.

Konishi, M., Akutagawa, E., 1985. Neuronal growth, atrophy and death in a sexually dimorphic song nucleus in the zebra finch brain. Nature 315, 145–147.

Konishi, M., Akutagawa, E., 1988. A critical period for estrogen action on neurons of the song control system in the zebra finch. Proceedings of the National Academy of Sciences of the United States of America 85, 7006–7007.

Krentzel, A.A., Ikeda, M.Z., Oliver, T.J., Koroveshi, E., Remage-Healey, L., 2019. Acute neuroestrogen blockade attenuates song-induced immediate early gene expression in auditory regions of male and female zebra finches. Journal of comparative physiology. A, Neuroethology, sensory, neural, and behavioral physiology.

Kuiper, G.G., Enmark, E., Pelto-Huikko, M., Nilsson, S., Gustafsson, J.A., 1996. Cloning of a novel receptor expressed in rat prostate and ovary. Proceedings of the National Academy of Sciences 93, 5925.

Kumar, S., Hedges, S.B., 1998. A molecular timescale for vertebrate evolution. Nature 392, 917–920.

Kurian, J.R., Olesen, K.M., Auger, A.P., 2010. Sex Differences in Epigenetic Regulation of the Estrogen Receptor-alpha Promoter within the Developing Preoptic Area. Endocrinology 151, 2297-2305.

Lambeth, L.S., Morris, K.R., Wise, T.G., Cummins, D.M., O’Neil, T.E., Cao, Y., Sinclair, A.H., Doran, T.J., Smith, C.A., 2016. Transgenic Chickens Overexpressing Aromatase Have High Estrogen Levels but Maintain a Predominantly Male Phenotype. Endocrinology 157, 83–90.

Lonning, P.E., Eikesdal, H.P., 2013. Aromatase inhibition 2013: clinical state of the art and questions that remain to be solved. Endocrine-Related Cancer 20, R183–R201.

Love, M.I., Huber, W., Anders, S., 2014. Moderated estimation of fold change and dispersion for RNA-seq data with DESeq2. Genome Biology 15, 550.

Lovell, P.V., Clayton, D.F., Replogle, K.L., Mello, C.V., 2008. Birdsong “Transcriptomics”: Neurochemical Specializations of the Oscine Song System. PLOS ONE 3, e3440.

Lovell, P.V., Huizinga, N.A., Friedrich, S.R., Wirthlin, M., Mello, C.V., 2018. The constitutive differential transcriptome of a brain circuit for vocal learning. BMC Genomics 19, 231.

Maggiolini, M., Picard, D., 2010. The unfolding stories of GPR30, a new membrane-bound estrogen receptor. J. Endocrinol 204, 104–114.

Martinkovich, S., Shah, D., Planey, S.L., 2014. Selective estrogen receptor modulators: tissue specificity and clinical utility. Clin Interv Aging 9, 1437–1452.

Mathews, G.A., Arnold, A.P., 1990. Antiestrogens Fail to Prevent the Masculine Ontogeny of the Zebra Finch Song System. Gen Comp Endocrinol 80, 48–58.

Mathews, G.A., Arnold, A.P., 1991. Tamoxifen’s Effects on the Zebra Finch Song System Are Estrogenic, not Antiestrogenic. J Neurobiol 22, 957–969.

Mathews, G.A., Brenowitz, E.A., Arnold, A.P., 1988. Paradoxical hypermasculinization of the zebra finch song system by an antiestrogen. Horm Behav 22, 540–551.

McCarthy, M.M., 2008. Estradiol and the developing brain. Physiological reviews 88, 91–124.

McCarthy, M.M., Arnold, A.P., 2011. Reframing sexual differentiation of the brain. Nat Neurosci 14, 677–683.

McDonnell, D.P., 2005. The Molecular Pharmacology of Estrogen Receptor Modulators: Implications for the Treatment of Breast Cancer. Clin Cancer Res 11, 871–877.

McEwen, B.S., Lieberburg, I., Chaptal, C., Krey, L.C., 1977. Aromatization: Important for sexual differentiation of the neonatal rat brain. Hormones and Behavior 9, 249–263.

Merten, M.D.P., Stocker-Buschina, S., 1995. Fadrozole induces delayed effects on neurons in the zebra finch song system. Brain Research 671, 317–320.

Mooney, R., Rao, M., 1994. Waiting periods versus early innervation: the development of axonal connections in the zebra finch song system. The Journal of Neuroscience 14, 6532.

Mosselman, S., Polman, J., Dijkema, R., 1996. ERβ: Identification and characterization of a novel human estrogen receptor. FEBS Letters 392, 49–53.

Nordeen, E.J., Nordeen, K.W., 1988. Sex and regional differences in the incorporation of neurons born during song learning in zebra finches. The Journal of Neuroscience 8, 2869.

Nottebohm, F., Arnold, A.P., 1976. Sexual dimorphism in vocal control areas of the songbird brain. Science 194, 211.

Odom, K.J., Hall, M.L., Riebel, K., Omland, K.E., Langmore, N.E., 2014. Female song is widespread and ancestral in songbirds. Nature Communications 5, 3379.

Olson, C.R., Hodges, L.K., Mello, C.V., 2015. Dynamic gene expression in the song system of zebra finches during the song learning period. Developmental Neurobiology 75, 1315–1338.

Panzica, G., Viglietti-Panzica, C., Balthazart, J., 2001. Sexual dimorphism in the neuronal circuits of the quail preoptic and limbic regions. Microscopy Research and Technique 54, 364–374.

Panzica, G.C., Castagna, C., Viglietti - Panzica, C., Russo, C., Tlemçani, O., Balthazart, J., 1998. Organizational effects of estrogens on brain vasotocin and sexual behavior in quail. Journal of neurobiology 37, 684–699.

Pfenning, A.R., Hara, E., Whitney, O., Rivas, M.V., Wang, R., Roulhac, P.L., Howard, J.T., Wirthlin, M., Lovell, P.V., Ganapathy, G., Mouncastle, J., Moseley, M.A., Thompson, J.W., Soderblom, E.J., Iriki, A., Kato, M., Gilbert, M.T.P., Zhang, G., Bakken, T., Bongaarts, A., Bernard, A., Lein, E., Mello, C.V., Hartemink, A.J., Jarvis, E.D., 2014. Convergent transcriptional specializations in the brains of humans and song-learning birds. Science 346, 1256846.

Pfizer, 2018. AROMASIN (exemestane) Product Monograph Kirkland, Quebec.

Pohl-Apel, G., Sossinka, R., 1984. Hormonal Determination of Song Capacity in Females of the Zebra Finch: Critical Phase of Treatment1. Zeitschrift für Tierpsychologie 64, 330–336.

Ralph, C.L., 1969. The Control of Color in Birds. American Zoologist 9, 521–530.

Riebel, K., 2009. Chapter 6 Song and Female Mate Choice in Zebra Finches: A Review, Advances in the Study of Behavior. Academic Press, pp. 197–238.

Ross, L.A., Del Bene, V.A., Molholm, S., Frey, H.P., Foxe, J.J., 2015. Sex differences in multisensory speech processing in both typically developing children and those on the autism spectrum. Frontiers in neuroscience 9, 185.

Sahores, A., Luque, G.M., Wargon, V., May, M., Molinolo, A., Becu-Villalobos, D., Lanari, C., Lamb, C.A., 2013. Novel, Low Cost, Highly Effective, Handmade Steroid Pellets for Experimental Studies. PLOS ONE 8, e64049.

Schlinger, B.A., Arnold, A.P., 1991. Androgen effects on the development of the zebra finch song system. Brain Research 561, 99–105.

Shaughnessy, D., Hyson, R., Bertram, R., Wu, W., Johnson, F., 2018. Female zebra finches do not sing yet share neural pathways necessary for singing in males. Journal of Comparative Neurology 527.

Simpson, H.B., Vicario, D.S., 1991a. Early estrogen treatment alone causes female zebra finches to produce learned, male-like vocalizations. J Neurobiol 22, 755–776.

Simpson, H.B., Vicario, D.S., 1991b. Early estrogen treatment of female zebra finches masculinizes the brain pathway for learned vocalizations. J Neurobiol 22, 777–793.

Somes, R.G., Smyth, J.R., 1967. Effects of Estrogen on Feather Phaeomelanin Intensity in the Fowl1,2. Poultry science 46, 26–32.

Takatoh, J., Nelson, A., Zhou, X., Bolton, M.M., Ehlers, M.D., Arenkiel, B.R., Mooney, R., Wang, F., 2013. New modules are added to vibrissal premotor circuitry with the emergence of exploratory whisking. Neuron 77, 346–360.

Tchernichovski, O., Nottebohm, F., Ho, C.E., Pesaran, B., Mitra, P.P., 2000. A procedure for an automated measurement of song similarity. Animal behaviour 59, 1167–1176.

Tehrani, M.A., Veney, S.L., 2018. Intracranial administration of the G-protein coupled estrogen receptor 1 antagonist, G-15, selectively affects dimorphic characteristics of the song system in zebra finches (Taeniopygia guttata). Developmental Neurobiology 78, 775–784.

Theodorsson, A., Hilke, S., Rugarn, O., Linghammar, D., Theodorsson, E., 2005. Serum concentrations of 17β-estradiol in ovariectomized rats during two times six weeks crossover treatment by daily injections in comparison with slow - release pellets. Scandinavian Journal of Clinical and Laboratory Investigation 65, 699–706.

Vahaba, D.M., Hecsh, A., Remage-Healey, L., 2019. Blocking neuroestrogen synthesis modifies neural representations of learned song without altering vocal imitation accuracy in developing songbirds. bioRxiv, 702704.

Vedder, L.C., Bredemann, T.M., McMahon, L.L., 2014. Estradiol replacement extends the window of opportunity for hippocampal function. Neurobiology of aging 35, 2183–2192.

Wade, J., Arnold, A.P., 1994. Post-hatching inhibition of aromatase activity does not alter sexual differentiation of the zebra finch song system. Brain Res 639, 347–350.

Walters, M.J., Harding, C.F., 1988. The effects of an aromatization inhibitor on the reproductive behavior of male zebra finches. Hormones and Behavior 22, 207–218.

Wang, R., Chen, C.-C., Hara, E., Rivas, M.V., Roulhac, P.L., Howard, J.T., Chakraborty, M., Audet, J.-N., Jarvis, E.D., 2015. Convergent differential regulation of SLIT-ROBO axon guidance genes in the brains of vocal learners. The Journal of comparative neurology 523, 892–906.

Wang, X., Chen, S., 2006. Aromatase Destabilizer: Novel Action of Exemestane, a Food and Drug Administration–Approved Aromatase Inhibitor. Cancer Research 66, 10281.

Wardell, S.E., Kazmin, D., McDonnell, D.P., 2012. Research resource: Transcriptional profiling in a cellular model of breast cancer reveals functional and mechanistic differences between clinically relevant SERM and between SERM/estrogen complexes. Mol Endocrinol 26, 1235–1248.

Wardell, S.E., Nelson, E.R., McDonnell, D.P., 2014. From empirical to mechanism-based discovery of clinically useful Selective Estrogen Receptor Modulators (SERMs). Steroids 90, 30–38.

Warren, W.C., Clayton, D.F., Ellegren, H., Arnold, A.P., Hillier, L.W., Kunstner, A., Searle, S., White, S., Vilella, A.J., Fairley, S., Heger, A., Kong, L., Ponting, C.P., Jarvis, E.D., Mello, C.V., Minx, P., Lovell, P., Velho, T.A., Ferris, M., Balakrishnan, C.N., Sinha, S., Blatti, C., London, S.E., Li, Y., Lin, Y.C., George, J., Sweedler, J., Southey, B., Gunaratne, P., Watson, M., Nam, K., Backstrom, N., Smeds, L., Nabholz, B., Itoh, Y., Whitney, O., Pfenning, A.R., Howard, J., Volker, M., Skinner, B.M., Griffin, D.K., Ye, L., McLaren, W.M., Flicek, P., Quesada, V., Velasco, G., Lopez-Otin, C., Puente, X.S., Olender, T., Lancet, D., Smit, A.F., Hubley, R., Konkel, M.K., Walker, J.A., Batzer, M.A., Gu, W., Pollock, D.D., Chen, L., Cheng, Z., Eichler, E.E., Stapley, J., Slate, J., Ekblom, R., Birkhead, T., Burke, T., Burt, D., Scharff, C., Adam, I., Richard, H., Sultan, M., Soldatov, A., Lehrach, H., Edwards, S.V., Yang, S.P., Li, X., Graves, T., Fulton, L., Nelson, J., Chinwalla, A., Hou, S., Mardis, E.R., Wilson, R.K., 2010. The genome of a songbird. Nature 464, 757–762.

Weis, S., Patil, K., Hoffstaedter, F., Nostro, A., Yeo, B.T.T., Eickhoff, S.B., 2019. Sex classification by resting state brain connectivity. bioRxiv, 627711.

Werling, D.M., Geschwind, D.H., 2013. Sex differences in autism spectrum disorders. Curr Opin Neurol 26, 146–153.

Whitehead, C.C., 2004. Overview of bone biology in the egg-laying hen. Poultry science 83, 193-199.

Whitney, O., Pfenning, A.R., Howard, J.T., Blatti, C.A., Liu, F., Ward, J.M., Wang, R., Audet, J.-N., Kellis, M., Mukherjee, S., Sinha, S., Hartemink, A.J., West, A.E., Jarvis, E.D., 2014. Core and region-enriched networks of behaviorally regulated genes and the singing genome. Science 346, 1256780.

Wobbrock, J., Findlater, L., Gergle, D., Higgins, J., 2011. The Aligned Rank Transform for Nonparametric Factorial Analyses Using Only ANOVA Procedures.

Wu, M.V., Manoli, D.S., Fraser, E.J., Coats, J.K., Tollkuhn, J., Honda, S.-I., Harada, N., Shah, N.M., 2009. Estrogen Masculinizes Neural Pathways and Sex-Specific Behaviors. Cell 139, 61–72.

