## Supplement Figures and Legends for "Estrogen and sex-dependent loss of the vocal learning system in female zebra finches"

__**Supplemental Material**__

**Supplemental Figures**

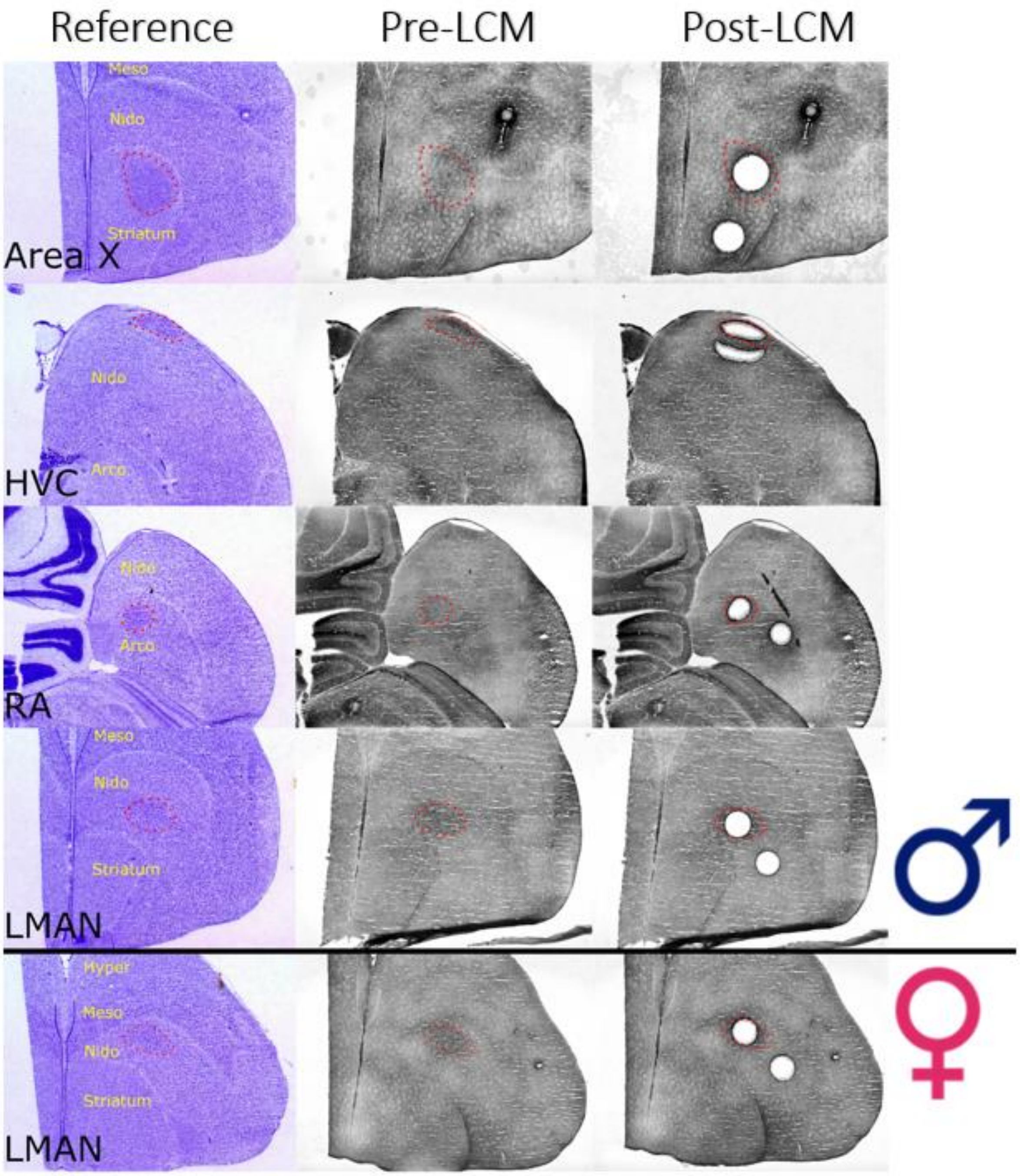


**Supplemental Fig. 1.** Example images of laser captured sections from vehicle PHD30 animals, ordered from top to bottom: male Area X and surrounding MSt, HVC and shelf, RA and LAi, and LMAN and LANido; and female LMAN and LANido. The sizes of the regions captured were made as similar as possible between the core of the song nucleus and the surrounding non-song area. Left: Brightfield cresyl violet stained reference slide. Middle: Before LCM. Right: after LCM.

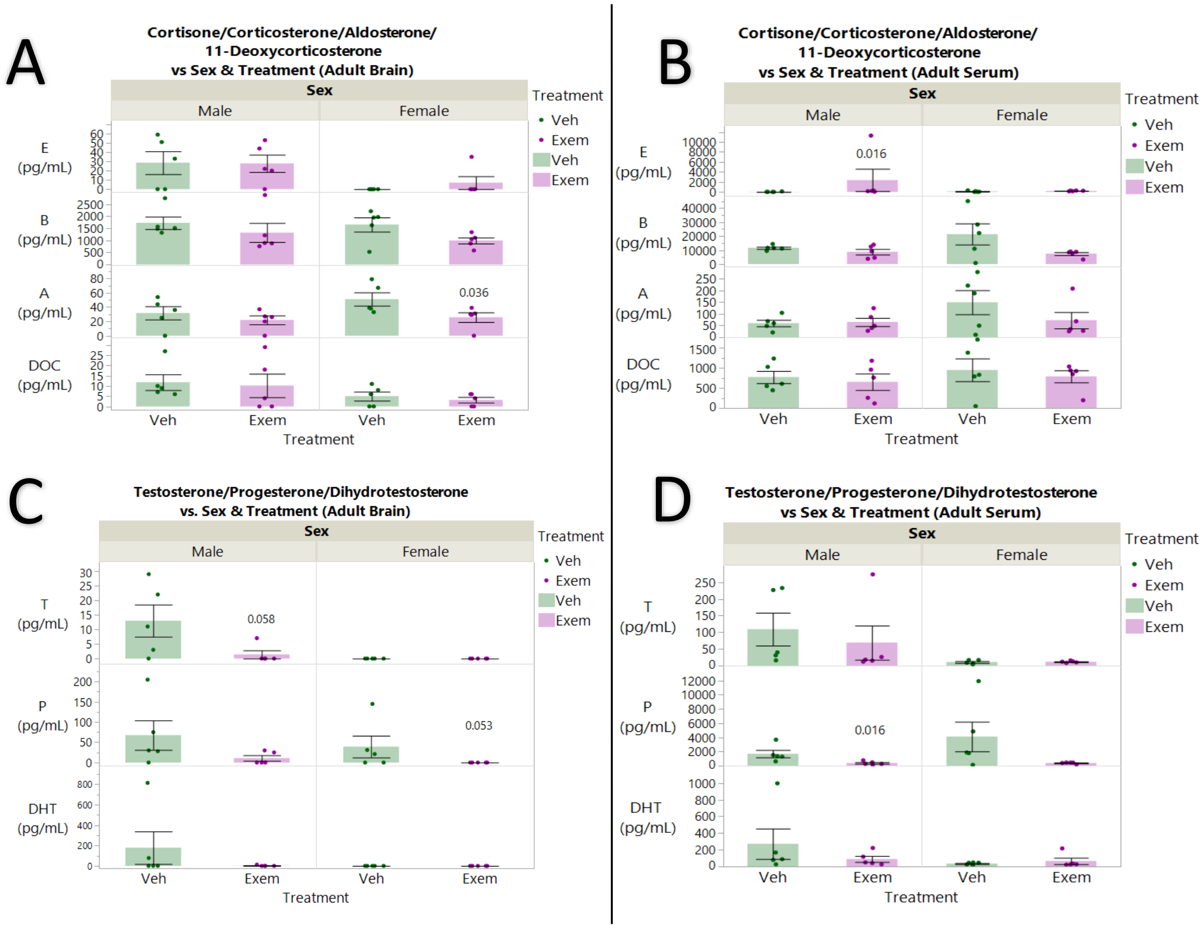


**Supplemental Fig. 2.** Complete steroid panel uHPLC-MS/MS for adult finches treated with vehicle or exemestane. (A) Adults Brain metabolic hormones. (B) Adult Serum metabolic hormones. (C) Adult Brain sex hormones. (D) Adult Serum sex hormones. Significant differences were determined with Wilcoxon test (n=5) followed by Steel-Dwass post-hoc tests. E: Cortisone, B: Corticosterone, A: Aldosterone, DOC: 11-Deoxycorticosterone, T: Testosterone, P: Progesterone, DHT: Dihydrotestosterone.

| Animal ID | # Clusters | SAP Plot | Sonogram (2.5-3.0 seconds) |
|---|---|---|---|
| Black532 Male Vehicle | 5 | 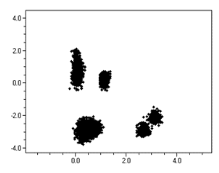 | 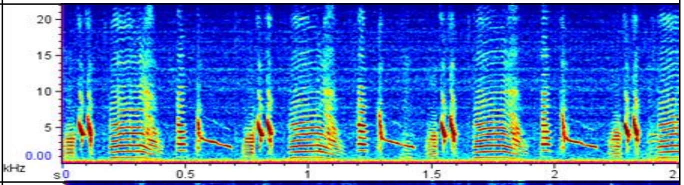 |
| Black533 Male Vehicle | 6 | 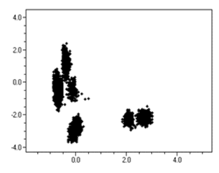 | 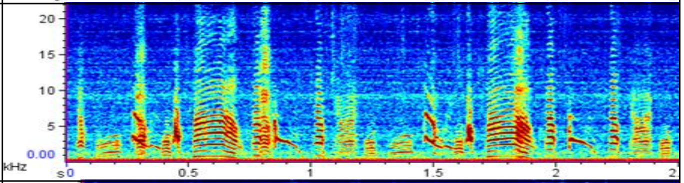 |
| Black534 Male Vehicle | 3 | 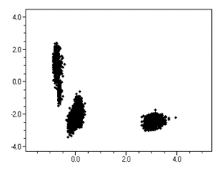 | 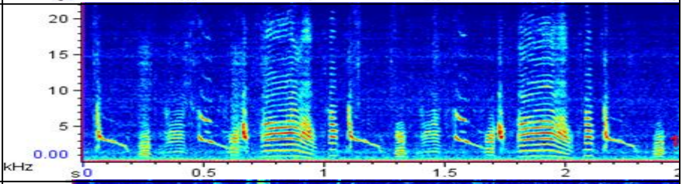 |
| Black514 Male Vehicle | 4 | 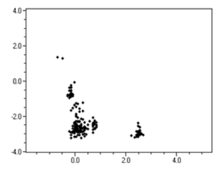 | 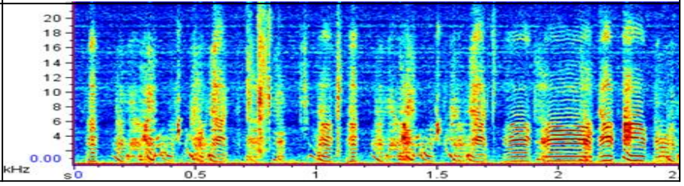 |

| Animal ID | # Clusters | SAP Plot | Sonogram (2.5-3.0 seconds) |
|---|---|---|---|
| Blue224 Male Vehicle | 5 | 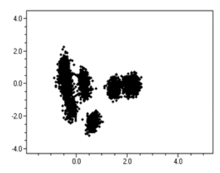 | 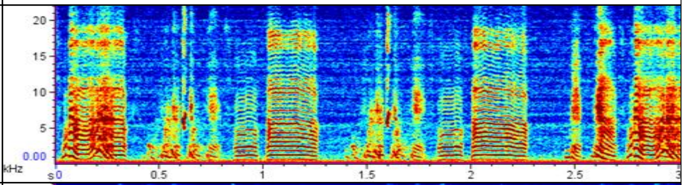 |
| Blue226 Male Vehicle | 6 | 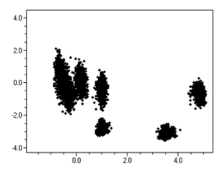 | 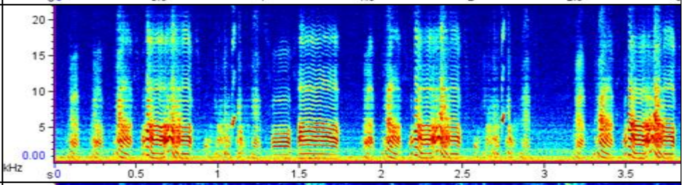 |
| Blue228 Male Vehicle | 3 | 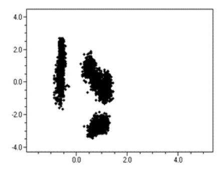 | 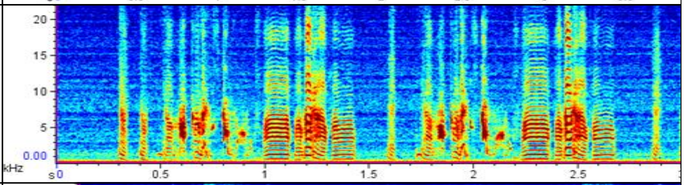 |
| Black283 Female Vehicle | 2 | 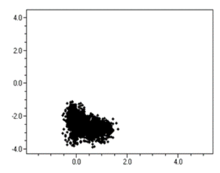 | 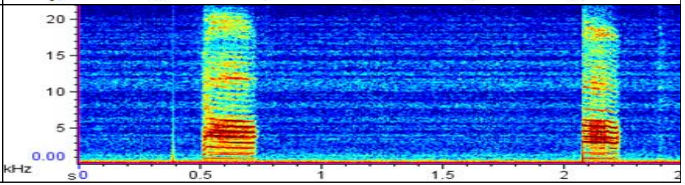 |

| Animal ID | # Clusters | SAP Plot | Sonogram (2.5-3.0 seconds) |
|---|---|---|---|
| Gold104 Female Vehicle | 2 | 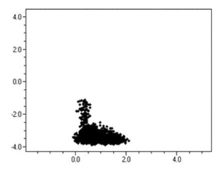 | 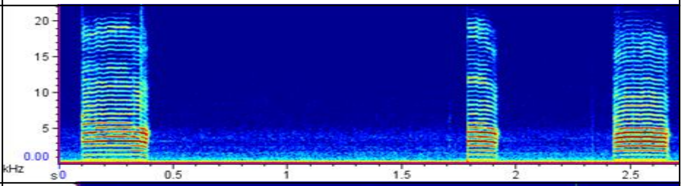 |
| Gold230 Female Vehicle | 2 | 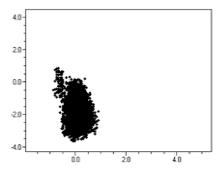 | 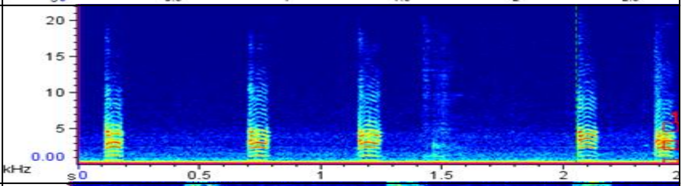 |
| Purple 348 Female Vehicle | 2 | 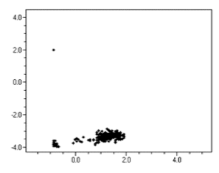 | 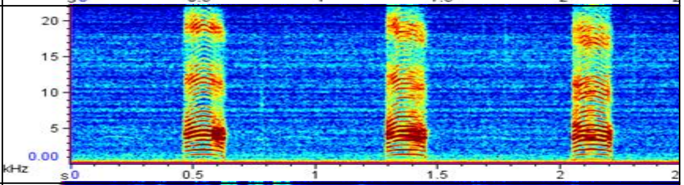 |
| Violet647 Male Exem | 3 | 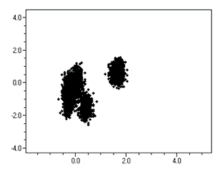 | 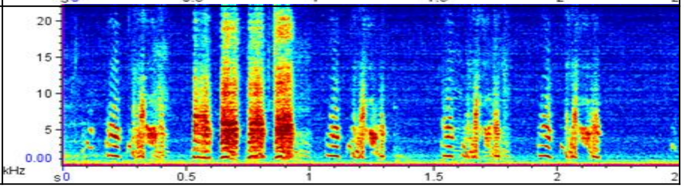 |

| Animal ID | # Clusters | SAP Plot | Sonogram (2.5-3.0 seconds) |
|---|---|---|---|
| violet648 Male Exem | 3 | 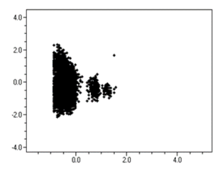 | 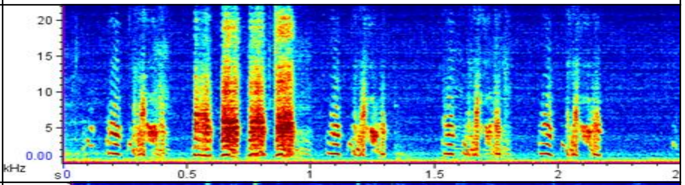 |
| violet653 Male Exem | 2 | 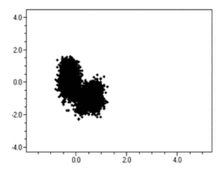 | 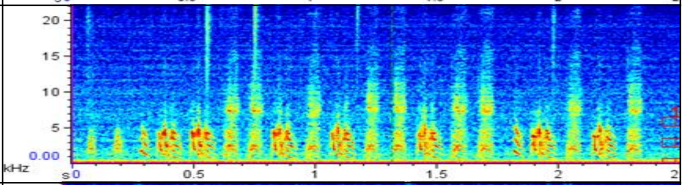 |
| violet655 Male Exem | 3 | 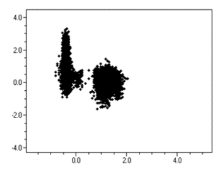 | 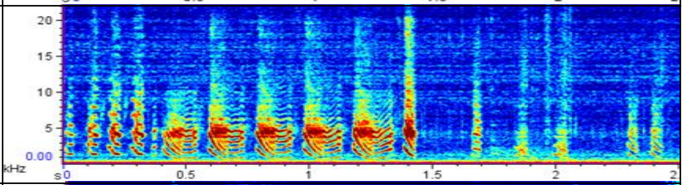 |
| Violet650 Female Exem | 1 | 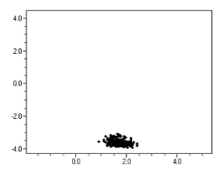 | 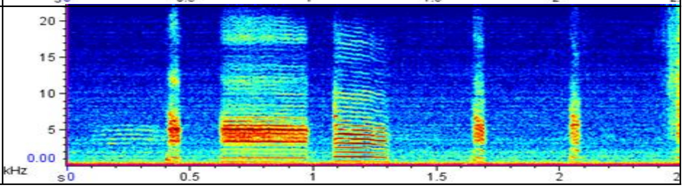 |

| Animal ID | # Clusters | SAP Plot | Sonogram (2.5-3.0 seconds) |
|---|---|---|---|
| Violet641<br>Female<br>Exem | 2 | 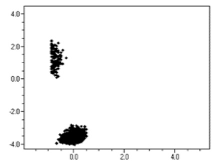 | 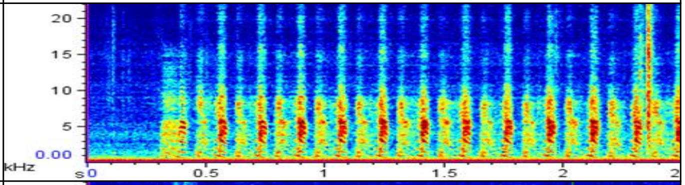 |
| Violet652<br>Female<br>Exem | 3 | 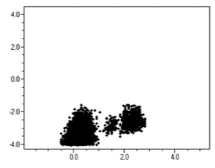 | 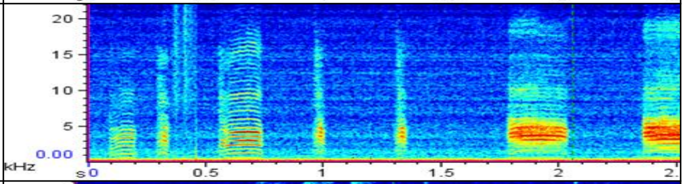 |
| Violet654<br>Female<br>Exem | 1 | 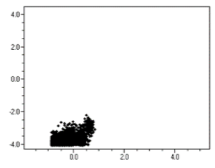 | 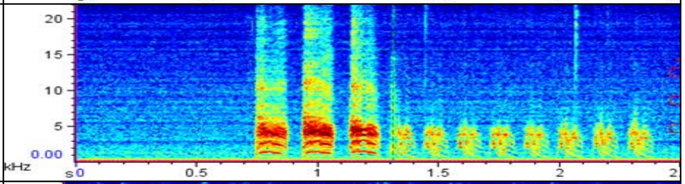 |
| Violet661<br>Male<br>E2 | 2 | 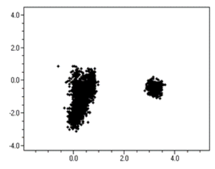 | 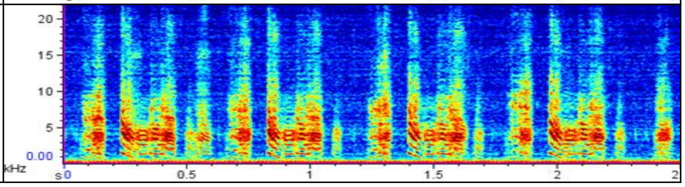 |

| Animal ID | # Clusters | SAP Plot | Sonogram (2.5-3.0 seconds) |
|---|---|---|---|
| Violet670<br>Male<br>E2 | 3 | 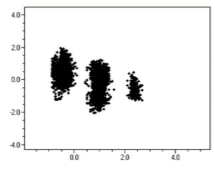 | 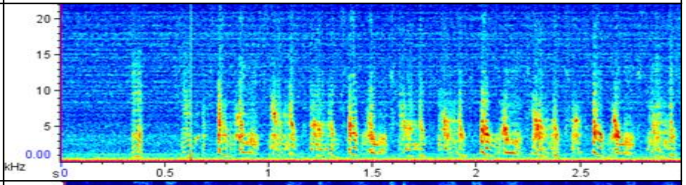 |
| Violet662<br>Male<br>E2 | 3 | 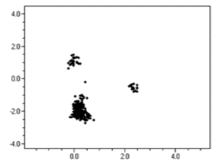 | 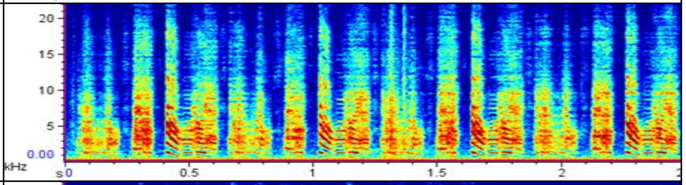 |
| Blue235<br>Male<br>E2 | 5 | 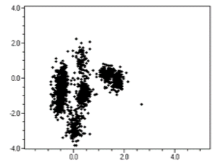 | 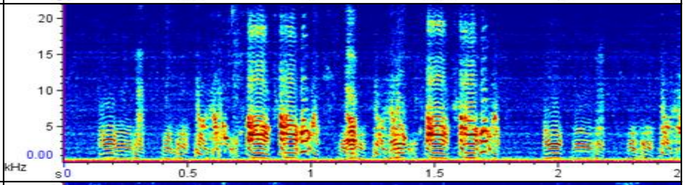 |
| Blue236<br>Female<br>E2 | 4 | 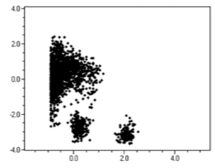 | 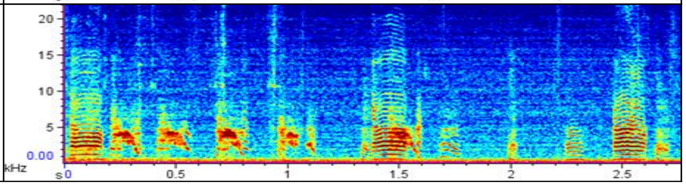 |

| Animal ID | # Clusters | SAP Plot | Sonogram (2.5-3.0 seconds) |
|---|---|---|---|
| Blue237<br>Female<br>E2 | 3 | 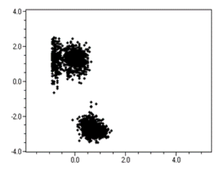 | 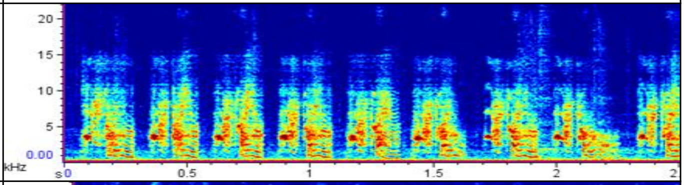 |
| Black324<br>Female<br>E2 | 4 | 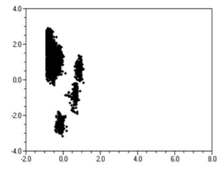 | 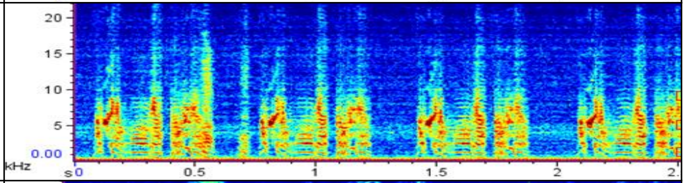 |
| Alpha1<br>Female<br>E2 | 3 | 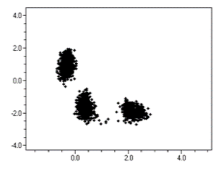 | 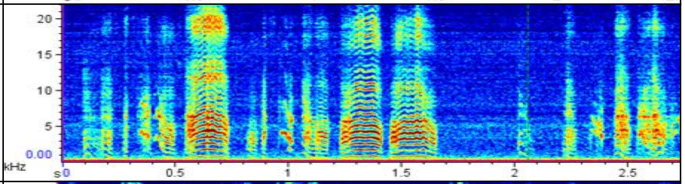 |
| Violet665<br>Female<br>E2 | 4 | 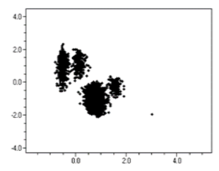 | 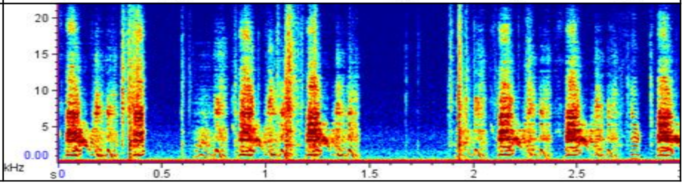 |

| Animal ID | # Clusters | SAP Plot | Sonogram (2.5-3.0 seconds) |
|---|---|---|---|
| Blue238<br>Female<br>E2<br>Implant lost | 4 | 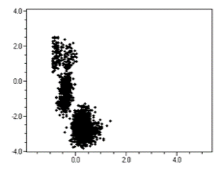 | 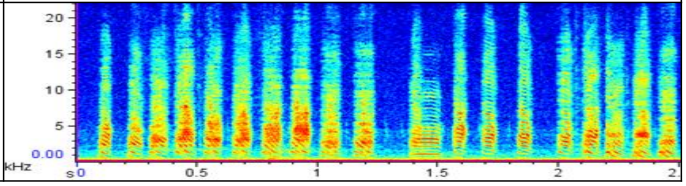 |

**Supplemental Fig. 3.** Sonograms and SAP2011 cluster plots of adult songs from control and estrogen manipulated animals during juvenile to adult development. Listed are individual animal identification, sex and treatment. All sonograms are from vocalizations produced in the presence of a novel female (directed song).

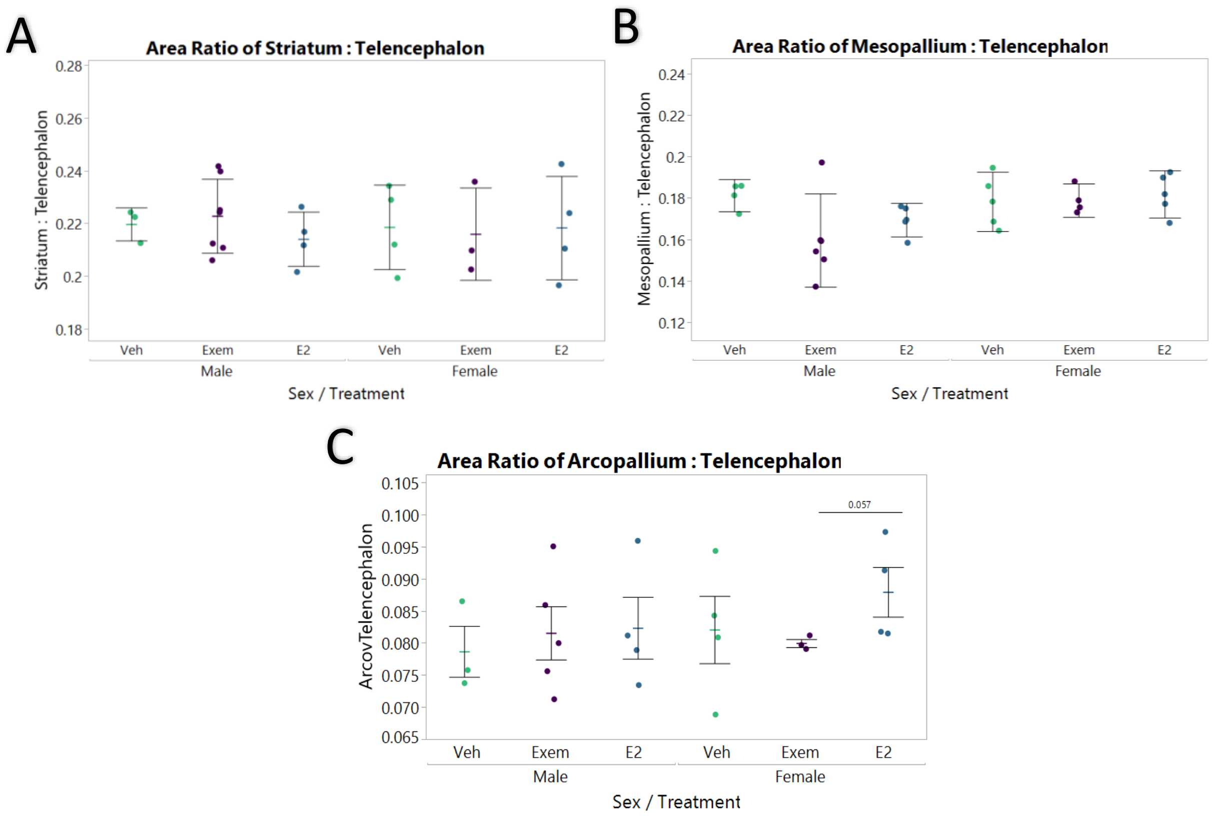


**Supplemental Fig. 4.** Relative area comparisons for different brain subdivisions. No differences were seen in these brain subdivisions relative to telencephalon size of the same brain section. There was a small difference of relatively smaller or less varied arcopallium size in exemestane treated females that approached significance (p = 0.057, Aligned ranks transformation ANOVA with post-hoc Wilcoxon comparison).

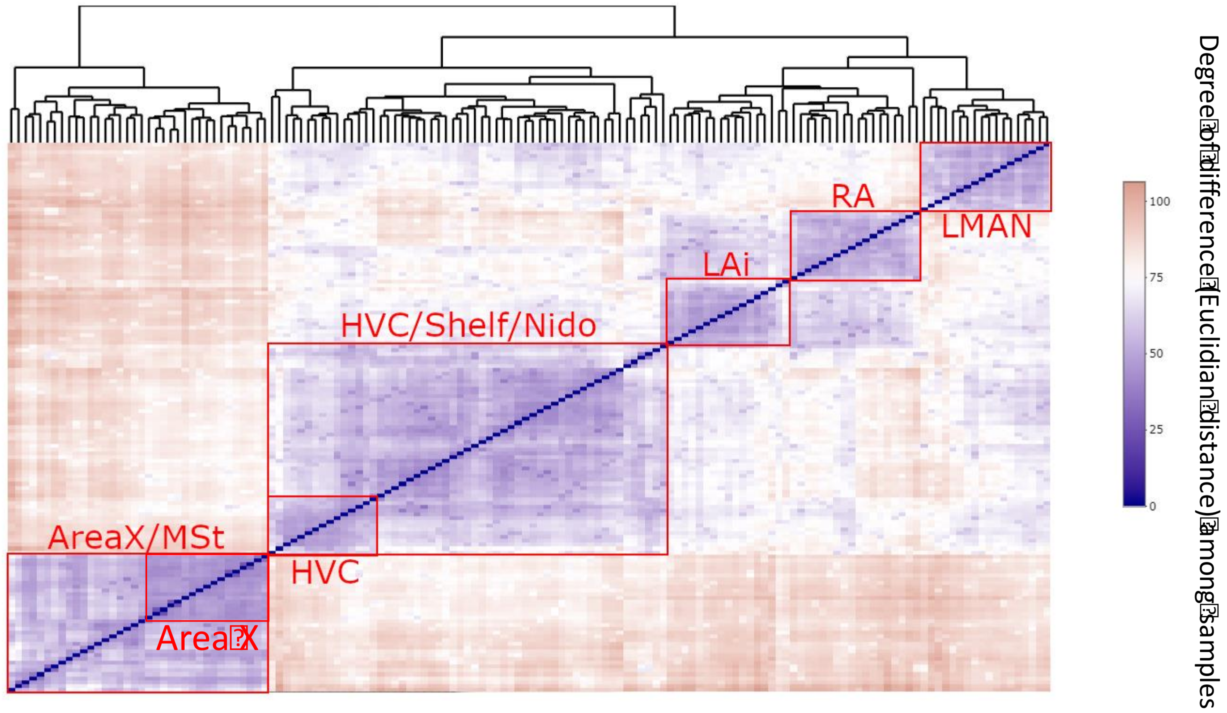


**Supplemental Fig. 5.** Sample to sample distance matrix of pharmacologically manipulated PHD30 finches. Euclidean distance is plotted for variance stabilized reads for each sequenced sample. 144 samples: 2 sexes, 3 treatments, 8 LCM collected regions, biological triplicates. Of the nidopallial samples, LMAN is most distant.

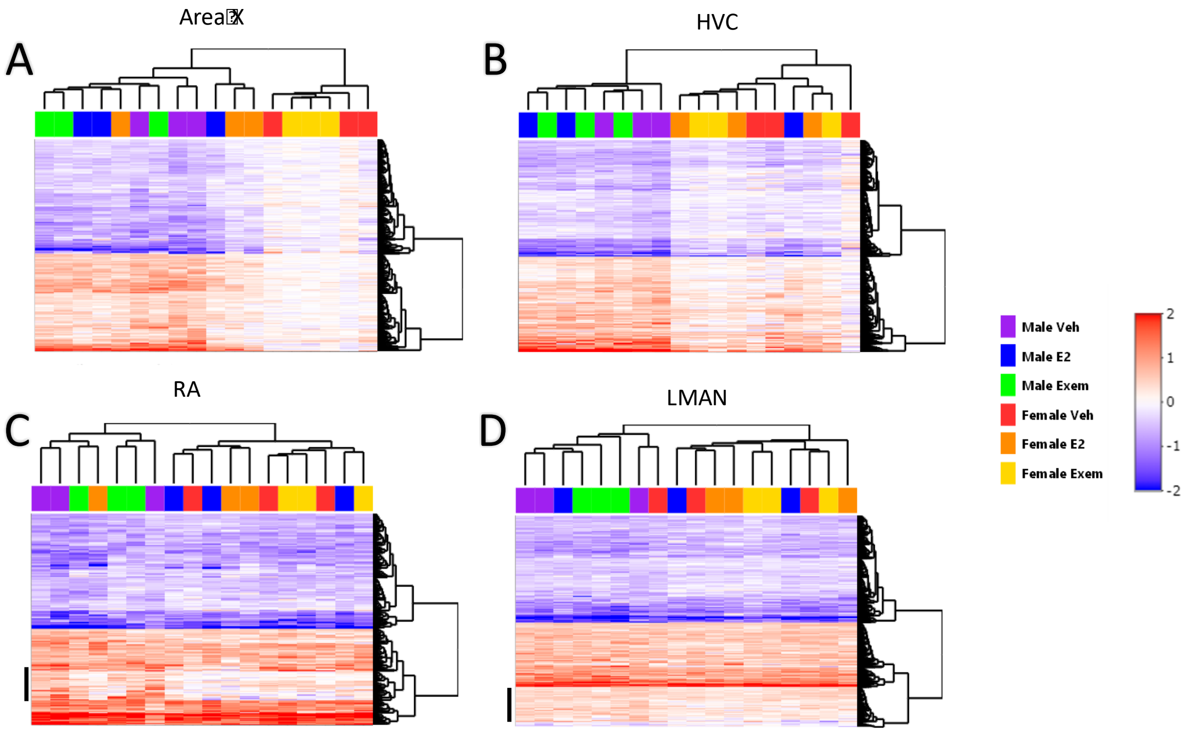


**Supplemental Fig. 6.** Hierarchically clustered gene expression heatmaps of individual animals. Genelist is from DEGs in vehicle males with FDR <0.05. (A) Vehicle male Area X has 326 DEGs. Hierarchical clustering shows a divide between individuals males + females treated with estradiol versus all other females, seen with nearly all genes. (B) Vehicle male HVC has 818 DEGs. Hierarchical clustering shows a divide between males and females regardless of treatment, save for one male treated with estrogen [blue]. One of the female vehicle samples appears that it might be different from other females [red on right]. (C) Vehicle male RA has 202 DEGs. Hierarchical clustering shows a weaker divide (small basal branch length) between males treated with vehicle and exemestane versus males treated with estradiol and all females but one treated with estradiol. The difference appears to be mostly due to a cluster of genes that is upregulated in males, but not in females or males treated with estradiol (black bar). (D) Vehicle male LMAN has 1023 DEGs. Hierarchical clustering shows a weaker divide (small basal branch length) similar to RA, but without as distinct a separation of groups. The heatmap pattern also is less distinct between groups than in the other three song nuclei. Heatmaps show differential expression in log2 fold change (log2FC) values for each gene (row) by individual (column). Values with log2FC greater than |2| were binned at |2|. Red, increased expression in song nuclei relative to the surround. Blue, decreased expression.

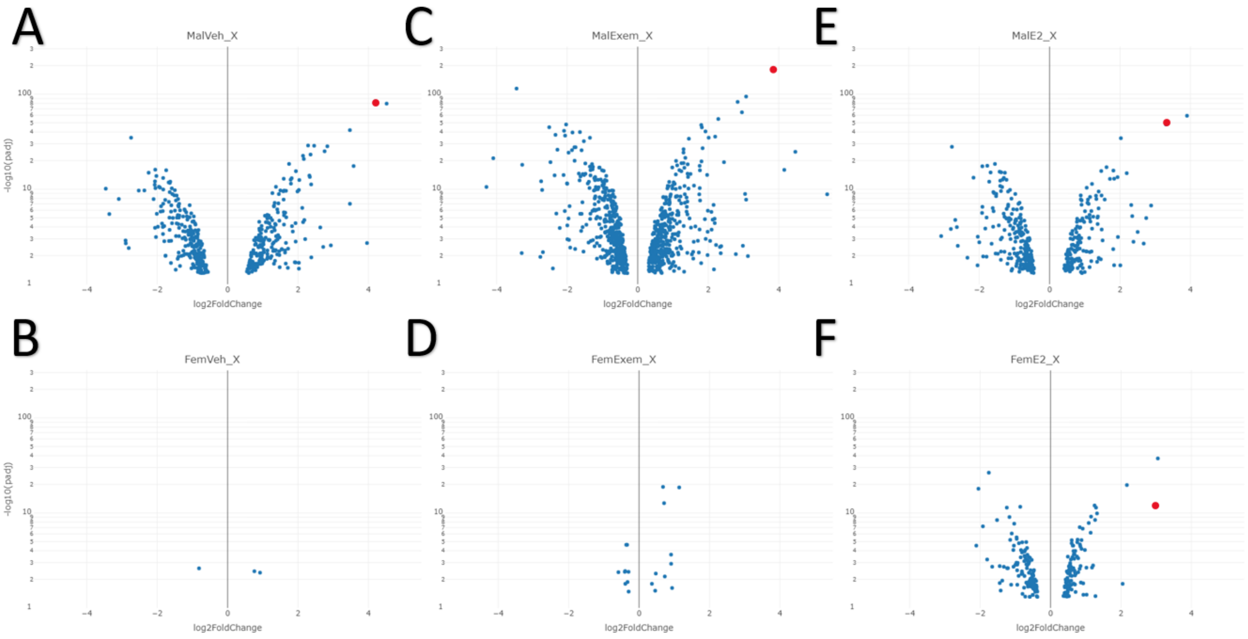


**Supplementary Fig. 7.** Fold changes and significance in DEGs between groups for Area X using pairwise statistical analyses in song nuclei versus the surrounding MSt. (A-F) Volcano plots of differentially expressed genes in Area X as compared to MSt from male and female finches treated with vehicle, estradiol, or exemestane. Only DEGs with FDR <0.05 were plotted. *CADPS2* is colored in red. X-axis is log2 fold change values with MSt enriched genes appearing to the left, and Area X enriched genes appearing to the right. Y-axis is the -log10 transformed FDR values on a log10 scale.

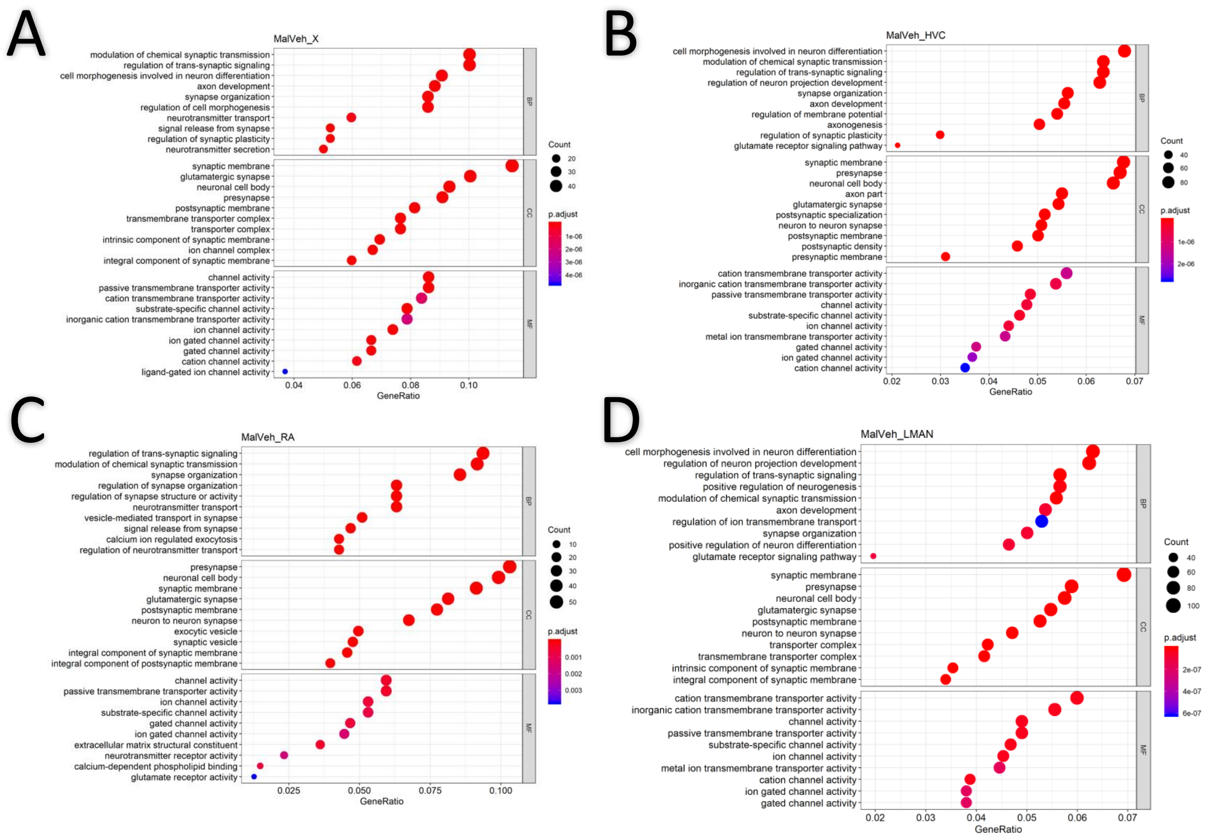

**Supplemental Fig. 8.** Top GO terms for DEGs in each song nucleus in vehicle males using pairwise statistical analyses in song nuclei versus the surrounding area. (A-D) Area X, HVC, RA, and LMAN respectively. X-axis indicates the ratio of genes with specialized expression out of the total list of genes, which contribute to each value. Count, number of DEGs that contributed to each category. BP, Biological Process; CC, Cell Compartment; MF, Molecular Function. Genelist is from all vehicle male DEGs for each region with FDR <0.05. GO enrichment results include p.adjusted value <0.1. Heatmap scale, level of significance.

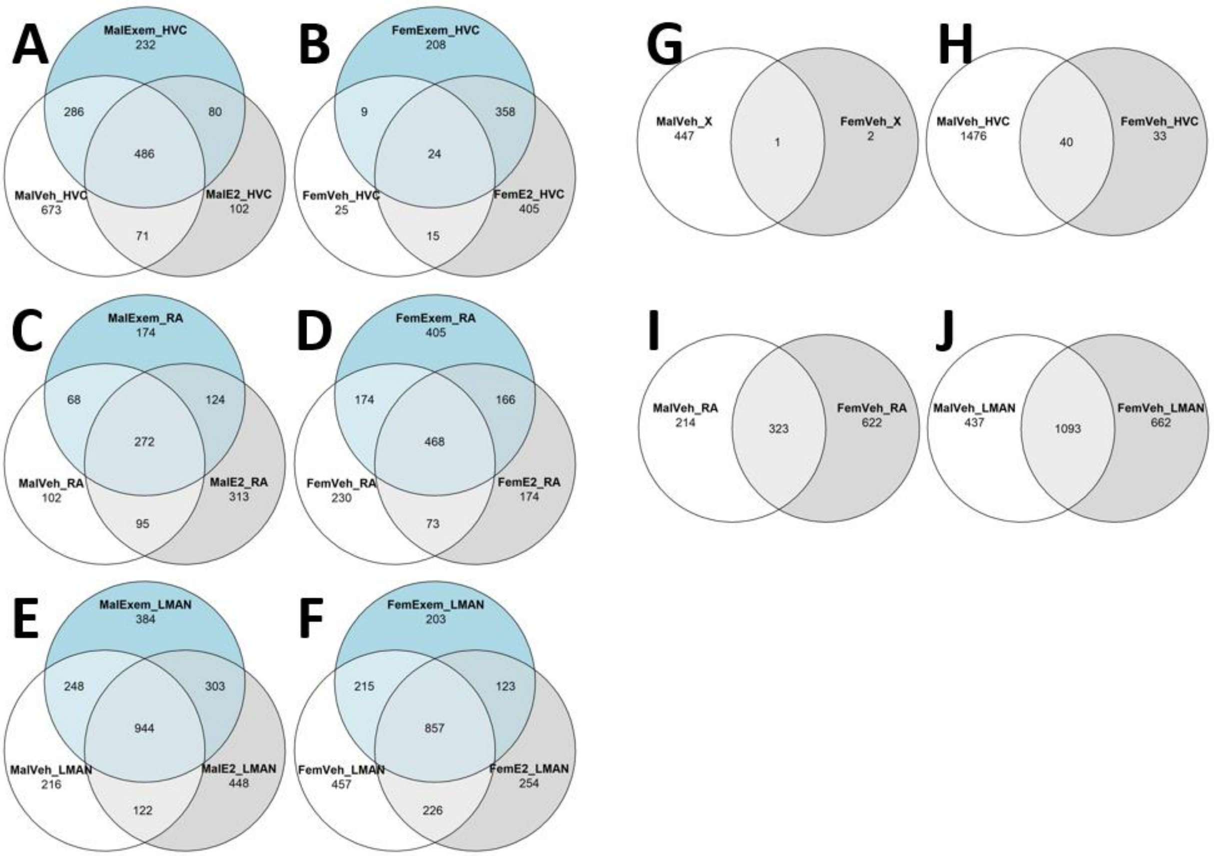


**Supplemental Fig. 9.** Venn diagram of DEGs by sex, region and phenotype using pairwise statistical analyses in song nuclei versus the surrounding area. Shown are the number of genes with specialized expression that overlap in (A) Male HVC, (B) Female HVC, (C) Male RA, (D) Female RA, (E) Male LMAN, and (F) Female LMAN. Number of overlaps between sexes in (G) vehicle treated Area X/Area X-analogous region, (H) vehicle treated HVC, (I) vehicle treated RA, and (J) vehicle treated LMAN. Genelist for each group is from all DEGs with FDR <0.05 for each nucleus/area per subject.

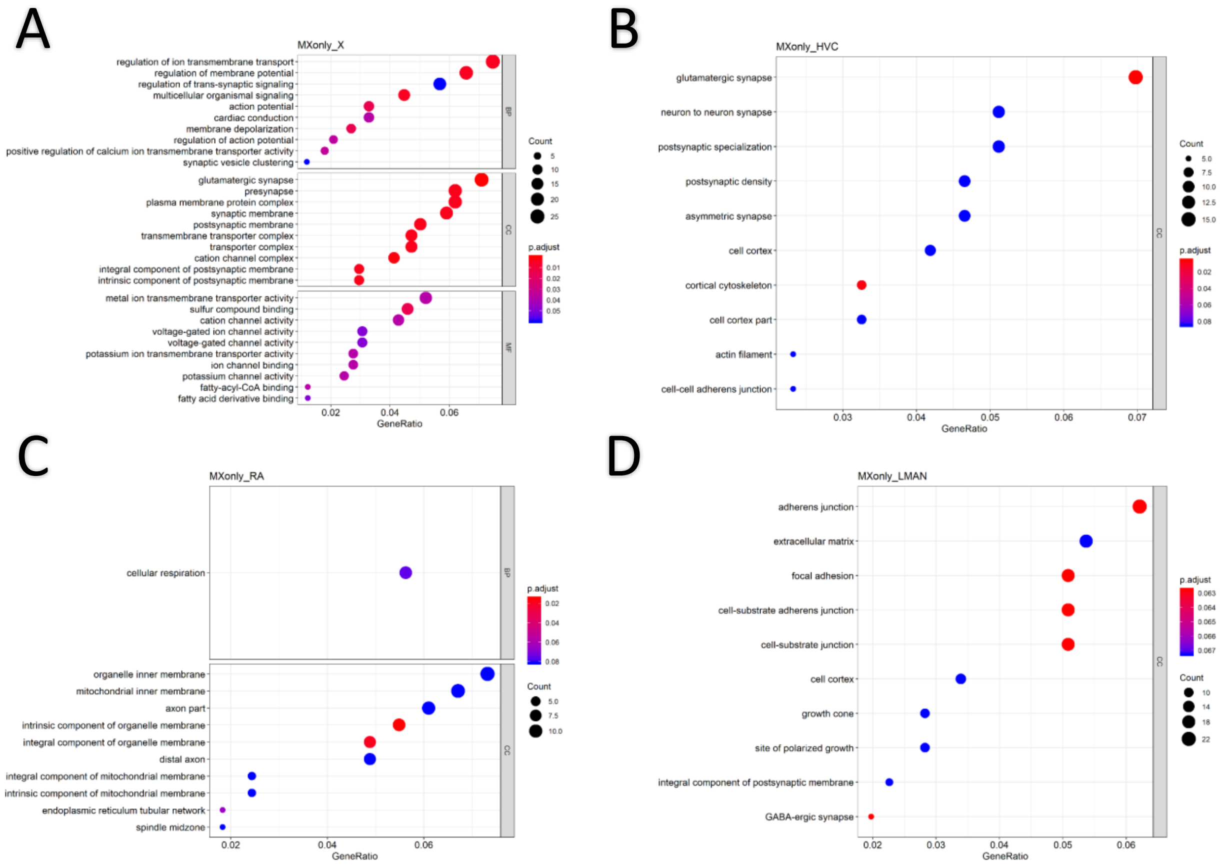
A
MXonly_X
regulation of ion transmembrane transport
regulation of membrane potential
regulation of trans-synaptic signaling
multicellular organismal signaling
action potential
cardiac conduction
membrane depolarization
regulation of action potential
positive regulation of calcium ion transmembrane transporter activity
synaptic vesicle clustering
glutamatergic synapse
presynapse
plasma membrane protein complex
synaptic membrane
postsynaptic membrane
transmembrane transporter complex
transporter complex
cation channel complex
integral component of postsynaptic membrane
intrinsic component of postsynaptic membrane
metal ion transmembrane transporter activity
sulfur compound binding
cation channel activity
voltage-gated ion channel activity
voltage-gated channel activity
potassium ion transmembrane transporter activity
ion channel binding
potassium channel activity
fatty-acyl-CoA binding
fatty acid derivative binding
BP
CC
MF
Count
5
10
15
20
25
p.adjust
0.01
0.02
0.03
0.04
0.05
0.02
0.04
0.06
GeneRatio
B
MXonly_HVC
glutamatergic synapse
neuron to neuron synapse
postsynaptic specialization
postsynaptic density
asymmetric synapse
cell cortex
cortical cytoskeleton
cell cortex part
actin filament
cell-cell adherens junction
CC
Count
5.0
7.5
10.0
12.5
15.0
p.adjust
0.02
0.04
0.06
0.08
0.03
0.04
0.05
0.06
0.07
GeneRatio
C
MXonly_RA
cellular respiration
organelle inner membrane
mitochondrial inner membrane
axon part
intrinsic component of organelle membrane
integral component of organelle membrane
distal axon
integral component of mitochondrial membrane
intrinsic component of mitochondrial membrane
endoplasmic reticulum tubular network
spindle midzone
BP
CC
p.adjust
0.02
0.04
0.06
0.08
Count
5.0
7.5
10.0
0.02
0.04
0.06
GeneRatio
D
MXonly_LMAN
adherens junction
extracellular matrix
focal adhesion
cell-substrate adherens junction
cell-substrate junction
cell cortex
growth cone
site of polarized growth
integral component of postsynaptic membrane
GABA-ergic synapse
CC
p.adjust
0.063
0.064
0.065
0.066
0.067
Count
10
14
18
22
0.02
0.03
0.04
0.05
0.06
GeneRatio

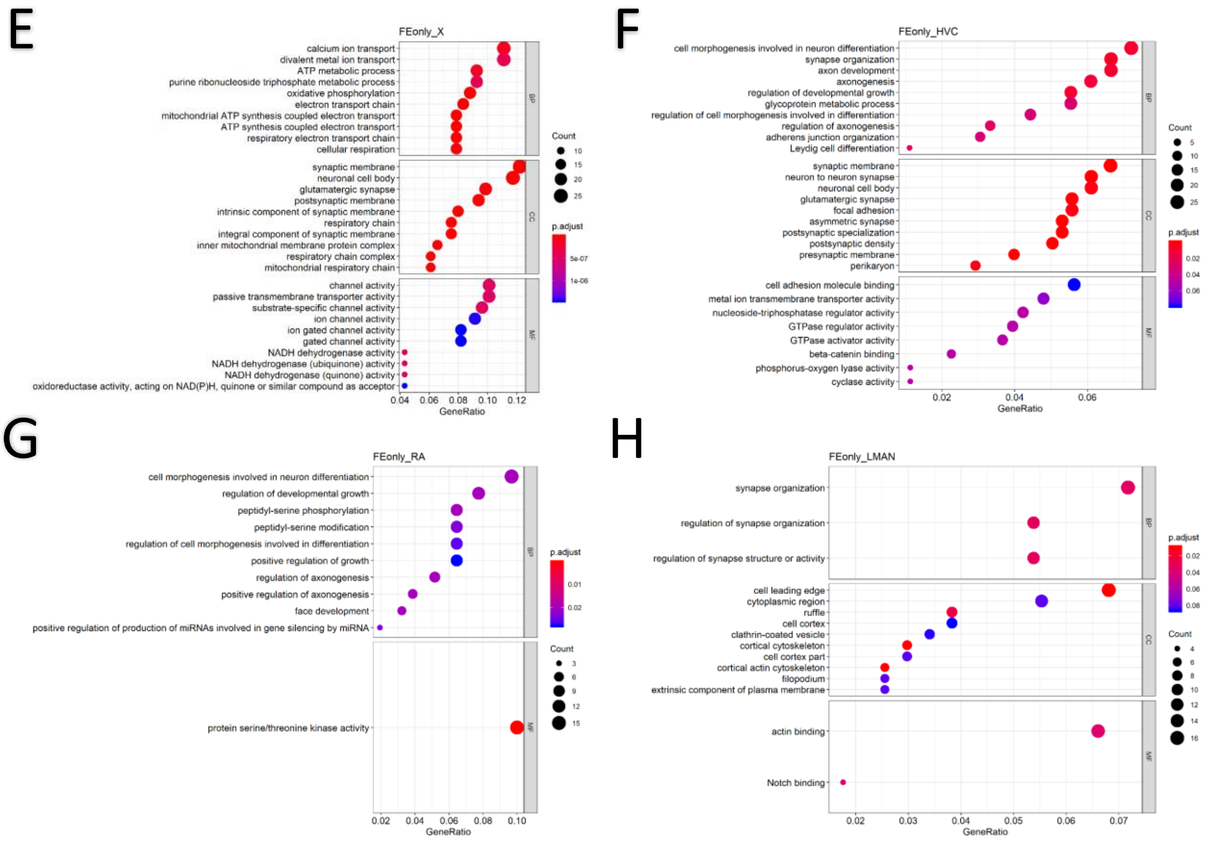


**Supplemental Fig. 10.** Top enriched GO terms for DEGs unique to exemestane treated males (A-D) and estradiol treated females (E-H) in each song nucleus. Genelist is from all exemestane male DEGs, excluding those shared with vehicle males and E2 males for the same region; or all estradiol female DEGs, excluding those shared with vehicle females and exemestane females within the same region. Genelist for each region was selected from DEGs with FDR <0.05. GO enrichment results include p.adjusted value <0.1.

**Supplemental Tables**

**Supplemental Table 1.** Statistics for all steroid levels in brain and serum of treated animals.

**Supplemental Table 2.** Subject to subject Spearman's correlation matrix of pharmacologically manipulated PHD30 finches. Each cell is the correlation of gene expression ratios of all genes from the respective nucleus in relation to the surrounding region, between pairs of animals. Red (1) – Blue (-1); High correlation to low correlation. Same sample comparison = 1. Female E2 HVC sample appears to be outlier in this analysis, although it is not in the PCA (**Fig. 7**). We believe this sample includes a proportion of ependymal cells overlying the ventricle space above HVC, since we found high levels of ependymal markers. Nevertheless, it did not prevent finding true positive specialized expression differences in males versus females in group analyses.

**Supplemental Table 3.** List of genes unique and overlapping between treatment groups within sex, within phenotype, or across all treatments and sex.

**Supplement Table 4.** Statistics for all song nucleus and brain region size analysis.

**Supplement Audio 1-9.** Sample audio clips from: Black324 (Female E2), Black532 (Male veh), Black534 (Male veh), Blue226 (Male veh), Blue237 (Female E2), Violet647 (Male exem), Violet648 (Male exem), Violet653 (Male exem), and Violet665 (Female E2).

**Supplement Movie 1.** Blue233 (Female E2) singing directed songs at a novel female behind electrochromic glass.

### Supplemental Note 1

#### UPLC/MS/MS Analysis of Zebra Finch Brain and Plasma Samples using the Biocrates Absolute*IDQ*® Stero17 Kit

Report from Duke Proteomics Core: Lisa St. John-Williams (sample preparation, data collection, data analysis, report writing), Will Thompson (study design, scientific oversight, report writing), and Arthur Moseley (scientific oversight, report writing).

The Absolute*IDQ* Stero17 assay quantifies 17 steroid hormones. The Stero17 kit includes all requisite calibration standards, internal standards, and QC samples. The use of these standards according to the detailed analysis protocol, which was validated in Biocrates' lab in Austria, assures assay harmonization and standardization within a project, across projects, and across laboratories. Selective analyte detection is accomplished by use of a triple quadrupole tandem mass spectrometer operated in Multiple Reaction Monitoring (MRM) mode in which specific precursor to product ion transitions are measured for every analyte and stable isotope labeled internal standard. There are two separate tandem mass spectrometric analyses of each sample. Sample analysis is performed by a UPLC (ultra-high pressure liquid chromatography) tandem MS method using a reversed phase analytical column for analyte separation (LC-MS/MS, **Figure 1**). A second UPLC/MS-MS method is used to acquire data for DHEAS (chromatogram not shown).

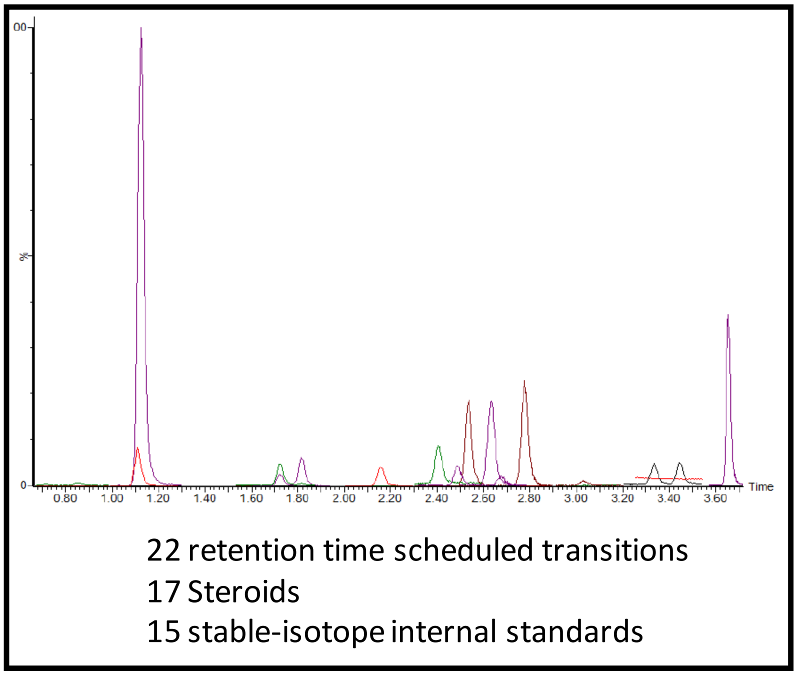


**Figure 1.** LC-MS/MS chromatogram showing a base-peak ion chromatogram of the analysis of steroids using the Biocrates Stero17 kit.

The calibration standards provided in the Biocrates Absolute*IDQ* kit were used for quantitation. Seven calibration standards are used for highly accurate and reproducible quantitation of the steroids as shown in **Figure 2**. Calibration standards were fit with a linear regression using $1/x^2$ weighting. **Figure 2** also shows a schematic and representative example of how the calibration curve would be used to back-calculate QC and sample concentrations for the different analytes.

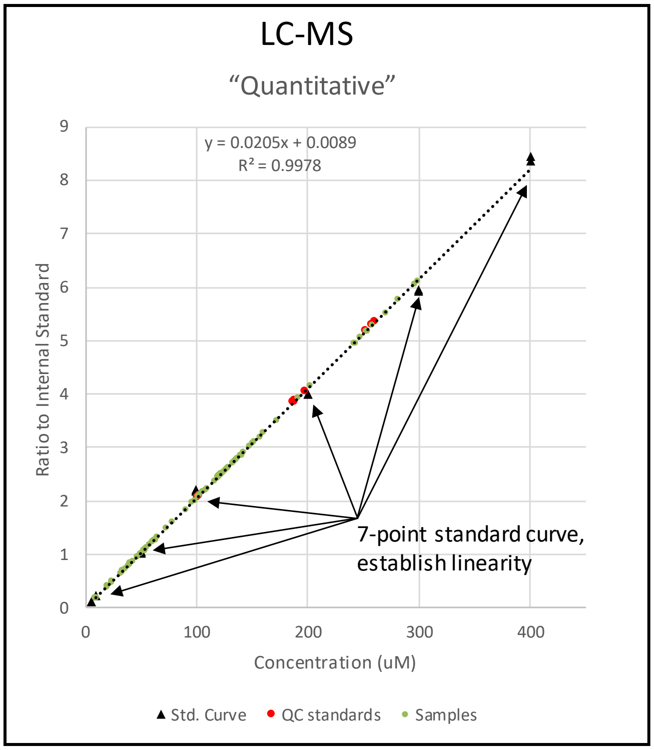


**Figure 2.** Schematic depicting the quantitative calibration methodology used in the Biocrates Stero17 kit for LC-MS/MS analysis

The samples were prepared in a 96-well plate format using the layout shown in **Figure 3**.

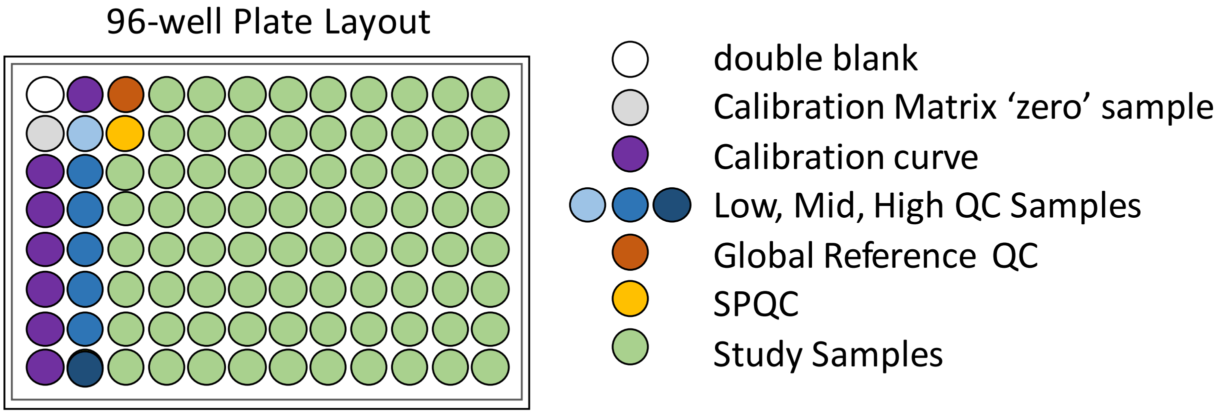


**Figure 3.** Schematic depicting the 96-well plate layout for the analysis of study samples including: blanks, calibration standards, and QC samples from Biocrates. Two additional QC samples were analyzed: the DPMCF Global Reference QC and the brain study sample pool QC (SPQC).

Brain Sample Preparation: Brain tissue samples were transferred into Precellys soft tissue homogenizing CK14 tubes (Bertin Technologies, Montigny-le-Bretonneux, France) and weighed. Each sample was diluted with 3 volumes of tissue extraction buffer (85:15 Ethanol:10mM Phosphate Buffer, v:v). For example, use 90µL tissue extraction buffer for 30mg brain tissue. Samples were then homogenized using 3x10-second pulses in the Precellys Evolution, between which samples were cooled for 60 seconds using the Cryolys function. The samples were then sonicated in an ice bath for 5 minutes and were stored at -80C until the day of sample extraction.

Plasma Sample Preparation: The typical volume of plasma extracted for steroid analysis is 500µL to allow the detection low abundance steroids. The volume of the plasma samples for this study ranged from 82µL to 377µL. Therefore, the sensitivity of the assay is as much as 6-fold less than expect due to sample volume limitation.

Sample Extraction: On the day of Stero17 sample extraction the plasma samples were thawed and vortexed. The brain homogenates were subjected to a single 10 second burst in the Precellys followed by centrifugation at 4°C for 10 minutes at 10,000rpm. The samples were then stored on ice until addition to the kit plate.

Samples were prepared using the Absolute*IDQ*® Stero17 kit (Biocrates Innsbruck, Austria) in accordance with their detailed protocol. A proprietary 96-well solid phase extraction (SPE) plate was provided with the kit. The SPE plate was washed with 1mL dichloromethane, 1 mL acetonitrile, 1mL methanol, and 1mL water. Blank Calibration Matrix, Biocrates calibration standards, Biocrates QC samples, brain study samples, and the SPQC sample were added to an empty 2mL 96-well plate in 500µL aliquots. Plasma samples were added to the plate in their entirety since they were all provided in aliquots of less than 500µL. After the addition of 10uL of the supplied Stero17 internal standard to the appropriate wells 400µL water was added to each well. Samples in the 2mL plate were mixed with 3 aspirate/dispense cycles using an 8-channel pipette then transferred to the appropriate wells of the SPE plate. Samples were allowed to elute by gravity for 10 minutes after which a gentle vacuum was applied. After the SPE plate was washed with 500µL water, it was dried under a stream of nitrogen while full vacuum was applied for one hour to ensure complete drying. Steroids were eluted from the SPE plate with 500µL dichloromethane into a 1mL 96-well collection plate. The eluent was dried under a stream of nitrogen at 50°C for 10 minutes. The dichloromethane elution and drying was repeated into the same wells of the 1mL plate. The dried dichloromethane eluents were reconstituted in 50µL 25% methanol in water, capped, sonicated in a water bath for 1 minute, and then shaken at 600rpm for 5 minutes.

Elution of DHEAS was then accomplished by adding two aliquots 200µL acetonitrile to the SPE plate and collecting the extracts in a second 96-well plate. The acetonitrile samples were diluted with 200µL water then shaken at 400rpm for 5 minutes.

A pool of equal volumes of all 20 brain homogenate samples analyzed was created (5226 SPQC). The pooled sample was prepared and analyzed in the same way as the study samples. From the plate, this sample was injected once before and once after the study samples in order to measure the performance of the assay across the sample cohort. The analyses of this pool can be used to assess potential batch effects. The order of injection of the samples is shown in **Figure 4**. There was insufficient volume of the plasma samples to prepare a plasma study pool.

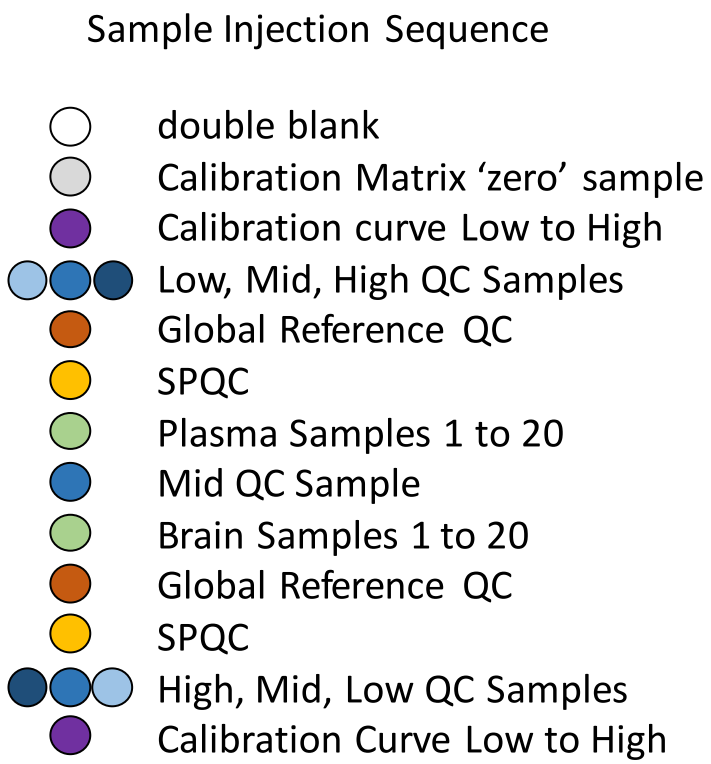


**Figure 4.** Schematic depicting the injection order of the samples for UPLC analysis. Dichloromethane extracts of each sample were injected alternately with the acetonitrile extracts of the same sample. Note that the Global Reference QC was prepared once and analyzed two times giving a measure of the analytical variability of the MS/MS analyses. The SPQC sample was also prepared one time and analyzed two times giving an additional measure of analytical variability.

Sample Analysis: UPLC separation of steroids (except for DHEAS) in the dichloromethane extracts was performed using a Waters (Milford, MA) Acquity UPLC with a proprietary column fitted with a proprietary guard column, both supplied by Biocrates. Analytes were separated using a gradient from a 30% proprietary aqueous solution to 85:10:5 Acetonitrile:Methanol:Water (v:v:v). Total UPLC analysis time was approximately 6 minutes per sample. DHEAS was analyzed by UPLC with total analysis time of approximately 6 minutes per sample using the same column and mobile phases. Using electrospray ionization in positive mode, samples were introduced directly into a Xevo TQ-S triple quadrupole mass spectrometer (Waters) operating in the Multiple Reaction Monitoring (MRM) mode. MRM transitions (compound-specific precursor to product ion transitions) for each analyte and internal standard were collected over the appropriate retention time. The UPLC-MS/MS data were imported into Waters application TargetLynx™ for peak integration, calibration, and concentration calculations. The UPLC-MS/MS data from TargetLynx™ were analyzed using Biocrates Met*IDQ*™ software.

<u>**Supplemental Note 2**</u>

#### R scripts for analysis after alignment with STAR/Kallisto

```
#Make column data matrix
#Made a much larger version for all samples when creating PCA plot/distance matrix. Instead of one Sex, one Tx, 2 areas, 3 subjects (6 samples), used two sex, three Tx, 8 areas, 3 subjects (144 samples).
diffExpression_labels_SexTx <- data.frame(
  FileName = c("Sex1_Tx_Surr","Sex2_Tx_Surr","Sex3_Tx_Surr",
        "Sex1_Tx_Nuc","Sex2_Tx_Nuc","Sex3_Tx_Nuc"),
  Subject = c("A","B","C","A","B","C"),
  Area = c(0,0,0,1,1,1))

#Run DESeq2 on the matrix of count data to output results as a data frame and a dds object
#Ran two sets of DE comparisons:
#Non-pairwise comparisons of area for average comparisons. Generates more stringent list for comparisons across group averages
#       design = ~Area
#Pairwise comparisons of area within subject for individual comparisons. Generates less stringent list for between group comparisons
#       design = ~Subject + Area
DiffExp2 <- function(Labels, Output_of_aggregation)
  dds <- DESeq2::DESeqDataSetFromMatrix(Output_of_aggregation, Labels, design = ~Subject + Area);
  dds <- DESeq(dds)
  res <- results(dds)
  tabl <- as.data.frame(res)
  tabl <- rownames_to_column(tabl, var = "id")
  result <- merge(Output_of_aggregation, tabl, by.x=0, by.y="id",sort=F)
  colnames(result)[1] <- "id"
  return(result)
}
DDS <- function(Labels, Output_of_aggregation) {
  dds <- DESeq2::DESeqDataSetFromMatrix(Output_of_aggregation, Labels, design = ~Subject + Area);
  dds <- DESeq(dds)
  return(dds)
}

#Get gene names from BioMart
mart <- useMart('ensembl')
TaeGutEnsembl <- useMart("ensembl", dataset = "tguttata_gene_ensembl")
genelist <- getBM(attributes = c("ensembl_gene_id", "external_gene_name"), mart = TaeGutEnsembl)

#Run DESEQ2 vst pipeline (see original documentation) on all of (144) the samples, and then run the PCA data with ggplot to see if samples cluster
#Repeat for Area X & MSt only by using Area X and MSt samples (36 samples)
DEGall <- DESeq2::DESeqDataSetFromMatrix(MergedCounts_byGroup, diffExpression_labels, design = ~Subject+Area)
```

```
DEGall <- DESeq(DEGall)
vDEGall <- vst(DEGall, blind=FALSE)
pcaAll <- plotPCA(vDEGall, intgroup=c("Area", "Sex","Treatment"), returnData=TRUE)
percentVar <- round(100 * attr(pcaAll, "percentVar"))
PCA<- ggplot(pcaAll, aes(PC1, PC2, stroke=1.5)) +
  geom_jitter(aes(shape=Treatment, color=Area, fill=Sex, size=Treatment)) +
  scale_shape_manual(values = c(21,24,23)) +
  scale_size_manual(values = c(4,4,4)) +
  scale_fill_manual(values = c("Black","Red")) +
  guides(fill = guide_legend(override.aes=list(size=3, color=c("Black","Red"))),
         shape = guide_legend(override.aes = list(size=3, stroke=1.5)),
         color = guide_legend(override.aes = list(size=1, stroke=2))) +
  xlab(paste0("PC1: ",percentVar[1],"% variance")) +
  ylab(paste0("PC2: ",percentVar[2],"% variance")) +
  stat_ellipse(aes(color=Area), type = "euclid", level = 2) +
  theme(legend.title = element_text(size=12),
        legend.text = element_text(size=10),
        axis.title.x=element_text(size=15),
        axis.title.y=element_text(size=15),
        axis.text.x=element_text(size=15),
        axis.text.y=element_text(size=15)) +
  theme(panel.background = element_rect(fill = 'gray95'))
coord_fixed()

#Each list of DEGs entered for generating the common gene list between STAR and Kallisto is combined,
will include single example:
STAR_Combined_X <- list(
  "MalVeh_X" = DEG_MV_mstX_paird,
  "MalE2_X" = DEG_ME_mstX_paird,
  "MalExem_X" = DEG_MX_mstX_paird,
  "FemVeh_X" = DEG_FV_mstX_paird,
  "FemE2_X" = DEG_FE_mstX_paird,
  "FemExem_X" = DEG_FX_mstX_paird)

#Function to filter and only keep DEG results with unique gene names (not just ensembl gene names)
#This version deletes duplicated entries
STAR_Filter_List_DEG <- function(List_Combined_DEG) {
  Filtered_list <- list()
  for (i in 1:length(List_Combined_DEG)) {
    current_tbl <- List_Combined_DEG[[i]]
    current_tbl <- subset(current_tbl, !duplicated(external_gene_name))
    filtered_tbl <- dplyr::filter(current_tbl, external_gene_name !="") %>%
      dplyr::select(external_gene_name, log2FoldChange, padj)
    Filtered_list[[i]] <- filtered_tbl
```

```
  }
  names(Filtered_list) <- names(List_Combined_DEG)
  return(Filtered_list)
}
Kallisto_Filter_List_DEG <- function(List_Combined_DEG) {
  Filtered_list <- list()
  for (i in 1:length(List_Combined_DEG)) {
    current_tbl <- List_Combined_DEG[[i]]
    current_tbl <- subset(current_tbl, !duplicated(id))
    filtered_tbl <- dplyr::filter(current_tbl, id !="") %>%
      dplyr::select(id, log2FoldChange, padj)
    Filtered_list[[i]] <- filtered_tbl
  }
  names(Filtered_list) <- names(List_Combined_DEG)
  return(Filtered_list)
}
#Can be used to filter and keep genes that are FDR <0.05 in each Sex/Tx/Brain region to create list of genes that are uniquely DE for each sex/tx/region.
#Will also pull the log2FC and pAdj from the STAR aligned data. These are not useful for making the common genes list, but used later for volcano plots.
Sig_genes_Only <- function(List_Filtered_list){
  Sig_list <- list()
  for (i in 1:length(List_Filtered_list)) {
    current_tbl <- List_Filtered_list[[i]]
    current_tbl <- dplyr::filter_at(current_tbl, vars(starts_with("pad")), any_vars(. < 0.05))
    Sig_list[[i]] <- current_tbl
  }
  names(Sig_list) <- names(List_Filtered_list)
  return(Sig_list)
}
#Only pull out genes that are common between the STAR and Kallisto lists
Pull_common <- function(STAR_list, Kallisto_list) {
  STAR_kallisto <- list()
  for (i in 1:6) {
    current_STAR <- STAR_list[[i]]
    current_Kallisto <- Kallisto_list[[i]]
    current_STAR_kallisto <- semi_join(current_STAR, current_Kallisto, by = c("external_gene_name" = "id"))
    STAR_kallisto[[i]] <- current_STAR_kallisto
  }
  names(STAR_kallisto) <- names(STAR_list)
  return(STAR_kallisto)
}
```

```
#Each item(dataframe) in the list will have genes that are DE with FDR<0.05 for each sex, tx and brain region. Male Veh DE genes are used as the standard for all downstream analysis, except for those that require DE genes of specific sex/tx/region.
#Example output
```

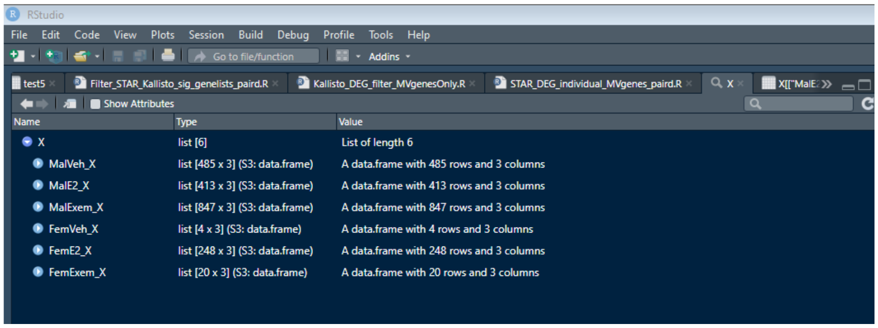


```
#Remove activity dependent genes (ADGs) identified in the Whitney et Al 2014 paper
removeADG <- function(common_list, ADG){
  no_ADG <- list()
  for (i in 1:6) {
    current <- common_list[[i]]
    current <- current[!current$external_gene_name %in% ADG$gene, ]
    no_ADG[[i]] <- current
  }
  names(no_ADG) <- names(common_list)
  return(no_ADG)
}

#DDS_rlog is for normalizing reads for individual DEG analysis
#This function will output the normalized reads for each region (both nucleus and surround) for each animal in the Sex/Tx group. Will only keep genes that are in the provided “GeneList”. Use any prior DEG results to bridge the ensembl symbols used in the STAR output (“Output_of_aggregation”) to the external_gene_names.
DDS_rlog <- function(Labels, Output_of_aggregation, GeneList, priorDEG) {
  Area <- factor(Labels$Area);#DESEQ2
  dds <- DESeq2::DESeqDataSetFromMatrix(Output_of_aggregation, Labels, design = ~Subject + Area);
  dds <- DESeq(dds)
  lds <- rlog(dds)
  sa <- SummarizedExperiment::assay(lds)
  ensembl_symbol <- left_join(GeneList, priorDEG, by= "external_gene_name")
  ensembl_DEGs <- as.data.frame(dplyr::select(ensembl_symbol, external_gene_name,
ensembl_gene_id))
  sa <- as.data.frame(sa)
  sa <- rownames_to_column(sa, "ensembl_gene_id")
  msa <- left_join(ensembl_DEGs, sa, by = "ensembl_gene_id")
```

```
  return(msa)
}

#LOG2(A/B) = LOG2(A) - LOG2(B)
#LOG2FC (nucleus vs Surround) = LOG2(nucleus) - LOG2(surround)
#Subtract matching columns (6$3, 7&4, 8&5) and write to new table with just the subject name
#Order of samples from individual areas are always the same, as defined by the diffExpression_labels made above
doMath <- function(DDS_results){
  DDS_results$Sub1 <- DDS_results[,6] - DDS_results[,3]
  DDS_results$Sub2 <- DDS_results[,7] - DDS_results[,4]
  DDS_results$Sub3 <- DDS_results[,8] - DDS_results[,5]
  Subnames <- substr(colnames(DDS_results[1,]), start = 1, stop = 8)
  Subonly <- DDS_results %>% dplyr::select(external_gene_name, Sub1, Sub2, Sub3)
  colnames(Subonly) <- Subnames[c(1, 3:5)]
  return(Subonly)
}

#Merge the LOG2FC generated with the doMath function
#This will give a large dataframe of columns (subjects). Made of 2 sex, 3 subjects, 3 Tx
mergeMath <- function(list_of_Mathed_subregion) {
  FC_individuals <- doMath(list_of_Mathed_subregion[[1]])
  for (i in 2:length(list_of_Mathed_subregion)) {
    FC_next <- doMath(list_of_Mathed_subregion[[i]])
    FC_individuals <- left_join(FC_individuals, FC_next, by = " external_gene_name ")
  }
  return(FC_individuals)
}

#Make values into buckets for heatmapping.
bucket <- function(DEG) {
  rownames(DEG) <- c()
  DEG <- column_to_rownames(DEG, var = "gene")
  DEG[DEG < -2] <- -2
  DEG[DEG > 2] <- 2
  return(DEG)
}

#Generate heatmap for rlog normalized DESeq2 results with each individual's fold change values, with values higher than 2log2FC (4-fold change) binned at 2log2FC by the bucket function above.
#Turn off seriation to try and keep male samples to the left
heatmaply(Nuc_res_FC_bucket, scale_fill_gradient_fun = scale_fill_gradient2(
  low="blue", high="red", mid="white", midpoint=0, limits = c(-2,2)), seriate = "none",
  Colv = TRUE, col_side_colors = c(rep(c("FV", "FE", "FX", "MV", "ME", "MX"), each=3)),
```

```
  side_color_layers = scale_fill_manual(values = c(
    "FV"="firebrick1", "FE"="darkorange", "FX"="gold", "MV"="purple", "ME"="blue", "MX"="green")),
  file = "Nuc_res_common_individuals_paird.html")

#Generate heatmap for rlog normalized DESeq2 results with group averages fold change values. Same as individual. Also no seriation.

heatmaply(Nuc_res_bucket, scale_fill_gradient_fun = scale_fill_gradient2(
        low="blue", high="red", mid="white", midpoint=0, limits = c(-2,2)), seriate = "none",
        Colv = TRUE, col_side_colors = c("MV", "ME", "MX", "FV", "FE", "FX"),
        side_color_layers = scale_fill_manual(values = c(
        "MV"="purple", "ME"="blue", "MX"="green", "FV"="firebrick1",
        "FE"="darkorange", "FX"="gold")), file = "Nuc_res_bucket_colors.html")


#Make Volcano plots using the log2FoldChange and pAdj columns from the common gene list made above. Using all genes specifically DE to Sex/Tx/region. Not using the vehicle male as a standard.
VP_html <- function(DEG_FDR, SexTxNucleus) {
  VP <- plot_ly(DEG_FDR, x = ~log2FoldChange, y= ~-log10(padj), type = "scatter", text = ~paste("ID: ",
external_gene_name)) %>%
    layout(title = sprintf("%s", SexTxNucleus))
  htmlwidgets::saveWidget(VP, sprintf("%s_VP_paird.html", SexTxNucleus))
}

#Make VennEuler plots using the common gene list made above. Use all the genes to compare what is DE between groups.
#VE plots are useless after 4 groups, so make several by a common variable: sex/Tx/song
Export_plot <- function(List_Venn, SexNucleus){
  Venn <- plot(venn(List_Venn))
  ggsave(sprintf("%s_Venn_paird.svg", SexNucleus), plot = Venn, width = 5, height = 5)
  ggsave(sprintf("%s_Venn_paird.png", SexNucleus), plot = Venn, width = 5, height = 5)
}


#Make a list of character vectors of genes from each Sex/Tx
#This format is needed for the clusterprofiler functions
MakeGL <- function(common_list){
  GL_list <- list()
  for (i in 1:6) {
    GL_list[[i]] <- common_list[[i]][,1]
  }
  names(GL_list) <- names(common_list)
  return(GL_list)
}
```

```
#Need to load the database made from NCBI data using AnnotationForge
#Similar to loading a library package, but from a local .sqlite file
#This makes an OrgDB type object that is required to run all the functions in clusterProfiler
#If you need to make/update this .sqlite source file, then run:
# makeOrgPackageFromNCBI(version = "1.0", author = "yourname", maintainer = "yourname",
outputDir = "C:/Users/name/Documents/", tax_id = "59729", genus = "Taeniopygia", species = "guttata")
#See AnnotationForge/AnnotationHub/AnnotationDbi documentation
ZForgDB <- loadDb("./org.Tguttata.eg.sqlite")

#Did not use finch OrgDb due to incomplete annotation. Had to default to human
library(org.Hs.eg.db)


#Function to run enrichGO on each character vector in the list
#Run the enrichGO function to get the GO terms associated with the genelist given
enrich_GList <- function(GL){
  enrich_GO_list <- list()
  for (i in 1:6) {
    enrich_GO_list[[i]] <- enrichGO(gene = GL[[i]], OrgDb = org.Hs.eg.db, keyType = "SYMBOL",
            ont = "ALL", pAdjustMethod = "BH", pvalueCutoff = 0.05, qvalueCutoff = 0.1)
  }
  names(enrich_GO_list) <- names(GL)
  return(enrich_GO_list)
}


#Make dotplots for each Sex/Tx
MakeDotplot_list <- function(enrichGO_list){
  for (i in 1:6) {
    Label <- names(enrichGO_list[i])
    dotplot(enrichGO_list[[i]], split = "ONTOLOGY", title = Label, showCategory = 10) +
      facet_grid(ONTOLOGY~., scale = "free") +
      ggsave(sprintf("%s_GO_dotplot_paird.svg", Label), width = 10, height = 7) +
      ggsave(sprintf("%s_GO_dotplot_paird.png", Label), width = 10, height = 7)
    }
}
```

#### Supplemental Note 3

### R packages

| Package Name | Repository (if not CRAN) | Release number | R Citations |
|---|---|---|---|
| Mixtools | | 1.1.0 | Benaglia, 2009 |
| Seqinr | | 3.4-5 | Charif, 2007 |
| Ape | | 5.3 | Paradis, 2018 |
| DESeq2 | Bioconductor | 1.24 | Love, 2013 |
| Lattice | | 0.20-38 | Sarkar, 2008 |
| MASS | | 7.3-51.4 | Venables, 2002 |
| Gplots | | 3.0.1.1 | Warnes, 2016 |
| Calibrate | | 1.7.2 | Graffelman, 2006 |
| Plotly | | 4.9.0 | Sievert, 2018 |
| Tidyverse | | 1.2.1 | Wickham, 2019 |
| Ggrepel | | 0.8.1 | Slowikowski |
| bioMart | Bioconductor | 2.40.0 | Durinck, 2009 |
| Heatmaply | | 0.16.0 | Galili, 2017 |
| Clusterprofiler | Bioconductor | 3.12.0 | Yu, 2012 |
| enrichPlot | Bioconductor | 1.4.0 | Yu, 2019 |
| AnnotationHub | Bioconductor | 2.16.0 | Morgan, 2019 |
| AnnotationDbi | Bioconductor | 1.46.0 | Pagès, 2019 |
| AnnotationForge | Bioconductor | 1.26.0 | Carlson, 2019 |
| Rcompanion | | 2.3.7 | Magniafico |
| FSA | | 0.8.25 | Ogle, 2019 |
| ARTool | | 0.10.6 | Kay, 2019 |
| Emmeans | | 1.4.2 | Length, 2019 |
| Eulerr | | 6.0.0 | Larsson, 2019 |
| Hmisc | | 4.3-0 | Harrell, 2019 |
